# Combining Plasma Extracellular Vesicle Let-7b-5p, miR-184 and Circulating miR-22-3p Levels for NSCLC Diagnosis and for Predicting Drug Resistance

**DOI:** 10.1101/2021.08.15.456411

**Authors:** G. P. Vadla, B. Daghat, A. Garcia, G. Perez, V. Ahmad, N. Patterson, Y. Manjunath, J.T. Kaifi, G. Li, C.Y. Chabu

## Abstract

Low-dose computed tomography (LDCT) Non-Small Cell Lung (NSCLC) screening is associated with high false-positive rates, leading to unnecessary expensive and invasive follow ups. There is a need for minimally invasive approaches to improve the accuracy of NSCLC diagnosis. Plasma extracellular vesicle (EV) and circulating microRNA (miRNA) have been proposed as cancer screening biomarkers. However, the identification of highly sensitive and broadly predictive core miRNA signatures remains a challenge. Also, how these systemic and diverse miRNAs impact cancer drug response is not well understood. Using an integrative approach, we examined plasma EV and circulating miRNA isolated from NSCLC patients versus screening controls with a similar risk profile. We found that combining EV (Hsa-miR-184, Let-7b-5p) and circulating (Hsa-miR-22-3p) miRNAs abundance robustly discriminates between NSCLC patients and high-risk cancer-free controls. Diagnosed NSCLC patients harboring sensitizing mutations in epidermal growth factor receptor EGFR (T790M, L578R) are treated with Osimertinib, a potent tyrosine kinase inhibitor (TKI). Nearly all patients develop TKI resistance via complex mechanisms and progress. We found that Hsa-miR-22-3p, Hsa-miR-184, and Let-7b-5p functionally converge on WNT/βcatenin and mTOR/AKT signaling axes, known cancer therapy resistance signals. Targeting Hsa-miR-22-3p and Hsa-miR-184 desensitized EGFR-mutated (T790M, L578R) NSCLC cells to Osimertinib. These findings suggest that the expression levels of circulating hsa-miR-22-3p combined with EV hsa-miR-184 and Let-7b-5p levels potentially define a core biomarker signature for improving the accuracy of NSCLC diagnosis. Importantly, these biomarkers have the potential to enable prospective identification of patients who are at risk of responding poorly to Osimertinib alone but likely to benefit from Osimertinib/AKT blockade combination treatments.

## Introduction

Lung cancer causes the most cancer-related deaths worldwide^1^, with 85% of lung cancer patients present with non-small cell lung cancer (NSCLC) and 70% of the cases are diagnosed as late-stage disease^1–3^. The 5-year survival rate for late-stage NSCLC is 2-5% compared to 92% for early-stage disease^1^, underscoring the importance of early detection.

Current lung cancer screening involves the use of the Lung imaging reporting and data system (Lung-RADS). This classification system relies on chest low-dose computed tomography (LDCT) scans: based on the numbers, size, appearance, and location of detected lung nodules, Lung-RADS assigns a score where Lung-RADS 1 means no lung nodules, Lung-RADS 2 and 3 represent probably benign lung nodules, whereas suspicious nodules with highest risk of cancer are categorized as Lung-RADS4) ^4^ However, LDCT has low specificity and a high false-positives rate (~23%)^5^. For patients who undergo multiple rounds of LDCT screening, the cumulative false-positive rate is estimated at 38-50% ^5,6^. In addition to the unnecessary emotional distress, these over-diagnosed patients are subjected to costly and invasive follow ups before they are confirmed lung cancer free. There is a need for minimally invasive strategies that permit early and accurate detection of NSCLC.

Patients who are diagnosed with NSCLC are stratified to chemotherapy and/or immunotherapy, or targeted therapy based on the presence or absence of known NSCLC driver mutations in tissue biopsy analyses ^7,8^. Depending on patients’ ethnicity, activating mutations of epidermal growth factor receptor (EGFR) tyrosine kinase is found in 15-50% of NSCLC ^9–11^. Patients harboring drug sensitizing EGFR mutations are treated with tyrosine kinase inhibitors or TKI. Some patients initially respond to tyrosine kinase inhibitors (TKI) but ultimately develop resistance: cancers acquire TKI-desensitizing EGFR mutations or activate compensatory signals, including WNT/β-catenin, the mechanistic target of rapamycin mTOR, and AKT signaling to drive cancer recurrence ^12–20^. Screening for genetic alterations that activate these signaling pathways provides a rationale for prioritizing treatment options that maximize the probability of achieving durable outcomes. However, it is becoming increasingly evident that cancer cells develop drug resistance via complex non-mutational mechanisms that involve emergent cell-cell interactions mediated by microRNAs (miRNAs) ^21–24^. miRNAs are short (19-23) non-coding nucleotides that degrade protein transcripts and fundamentally impact signaling events ^25–27^. These miRNAs are released directly into the blood circulation (circulating miRNA) or secreted as extracellular vesicles (EV) cargo to elicit signaling events in target cells ^28,29^. The potential for EV miRNA as minimally-invasive liquid biomarkers is now widely recognized ^30,31^.

In this study, we integrated EV and circulating blood miRNA analyses to identified EV and miRNA combined features that robustly differentiate cancer patients from disease-free controls. Further, the identified miRNAs converge on WNT/β-catenin and AKT/mTOR signaling pathways, suggesting a role in drug-resistance and highlighting their potential as biomarkers for predicting therapy response and for selecting patients that will likely benefit from AKT blockade.

## Results

### CD9 and CD63, but not CD81, are enriched on NSCLC patients extracellular vesicles

There is a critical need for sensitive, robust, and minimally-invasive tools for improving the accuracy of NSCLC diagnosis. To this end, we interrogated plasma extracellular vesicles and miRNA to identify molecular signatures that can discriminate between NSCLC patients and healthy individuals with similar risk profiles. The US Preventive Services Task Force recommends LDCT screening for at-risk individuals defined as 55-80 years of age with 30 or more pack-year smoking history or have quit within the past 15 years ^32^. Individuals ranging between 42 and 62 years of age with a qualifying smoking history were divided into two groups based on their initial LDCT Lung-RADS scores 2 or 4 (Table 1, Lung-RADS2 screening controls versus Lung-RADS4, *N*=20 per group).

**Table 1.** Participants profile. Table showing each patient’s description including, age, gender, smoking history, and cancer progression status. False positive diagnoses are denoted with an asterisk (*) in the Group column. Group A and B contain individuals with lung-RADS2 or lung-RADS4 initial LDCT findings, respectively.

| Sample ID | Age (yrs, as of 2019) | Gender | Smoking History (pack - year) | Group | Lung-RAD Score |  | Death Cancer Related |
| --- | --- | --- | --- | --- | --- | --- | --- |
|  |  |  |  |  | INITIAL | LATEST |  |
| 82 | 62 | M | 40 | A | 2 | 4A | None |
| 83 | 64 | F | 40 | A | 2 | 2 | None |
| 84 | 62 | F | 45 | A | 2 | 2 | None |
| 85 | 62 | F | 30 | A | 2 | 2 | None |
| 88 | 45 | F | 80 | A | 2 | 2 | None |
| 91 | 51 | F | 40 | A | 2 | 2 | None |
| 96 | 58 | F | 45 | A | 2 | 2 | None |
| 98 | 56 | F | 45 | A | 2 | 2 | None |
| 99 | 60 | M | 35 | A | 2 | 2 | None |
| 100 | 61 | F | 30 | A | 2 | 4A | None |
| 102 | 53 | F | 40 | A | 2 | 2 | None |
| 104 | 58 | M | 50 | A | 2 | 2 | None |
| 105 | 53 | F | 80 | A | 2 | 2 | None |
| 106 | 48 | M | 45 | A | 2 | 2 | None |
| 111 | 57 | F | 60 | A | 2 | 2 | None |
| 114 | 61 | M | 30 | A | 2 | 3 | None |
| 120 | 63 | F | 40 | A | 2 | 4A | None |
| 121 | 57 | F | 90 | A | 2 | 3 | None |
| 123 | 50 | F | 40 | A | 2 | 2 | None |
| 124 | 61 | M | 50 | A | 2 | 2 | None |
| 8 | 50 | F | 100 | B | 4 | 4B | None |
| 12 | 50 | F | 55 | B | 4 | 4X | None |
| 15 | 56 | F | 45 | B | 4 | 4B | None |
| 16 | 42 | F | 40 | B | 4 | 4B | None |
| 30 | 49 | F | 55 | B | 4 | 4A | Yes |
| 40 | 55 | M | 45 | B* | 4 | 4B | None |
| 52 | 44 | F | 50 | B | 4 | 4B | None |
| 53 | 58 | M | 27 | B | 4 | 4B | None |
| 70 | 45 | M | 50 | B | 4 | 4X | None |
| 92 | 61 | M | 40 | B* | 4 | 4A | None |
| 115 | 51 | F | 40 | B | 4 | 4B | None |
| 135 | 50 | M | 25 | B* | 4 | 4B/4X | None |
| 138 | 49 | F | 100 | B | 4 | 4X | Yes |
| 139 | 52 | F | 40 | B* | 4 | 4A | None |
| 149 | 57 | M | 40 | B* | 4 | 4B | None |
| 152 | 49 | F | 25 | B | 4 | 4B | None |
| 159 | 60 | M | 60 | B | 4 | 4B | None |
| 170 | 53 | F | 40 | B | 4 | 4B | None |
| 171 | 55 | M | 80 | B* | 4 | 2 | None |
| 183 | 56 | F | 40 | B | 4 | 4X | None |

We isolated plasma EV from all 40 individuals (Figure 1a) and performed nanoparticle tracking analyses (NTA) to determine whether EV from Lung-RADS4 patients cohort exhibit physical characteristics that are distinct from control cohort EV. Initial analyses revealed that EV density was reduced in Lung-RADS4 patients compared to screening controls but these EV were larger than Lung-RADS2 EV (Figure 1b, c).

**Figure 1.**
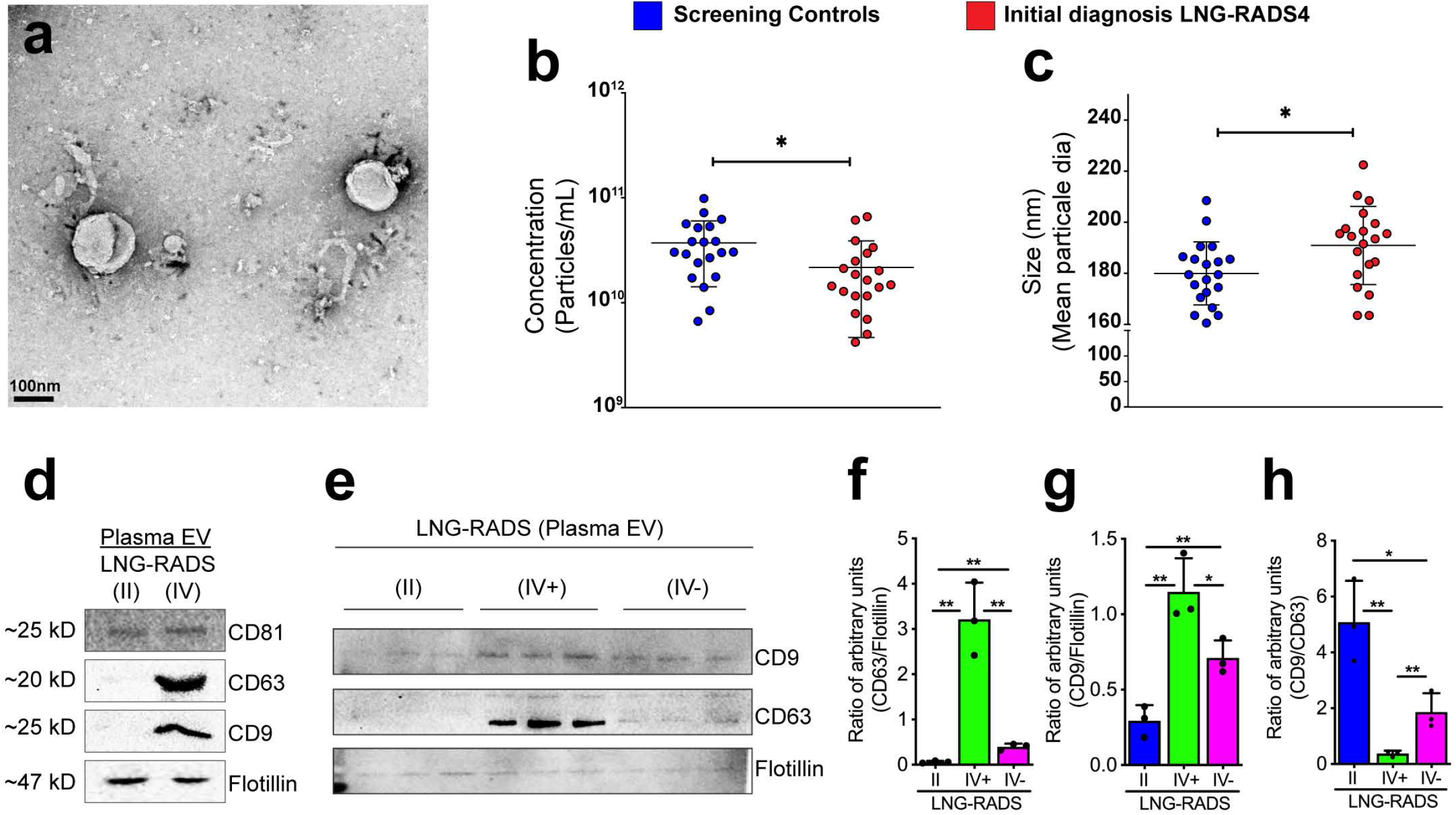
CD9 and CD63, but not CD81, are enriched on NSCLC EV. a) Representative transmission electron micrograph of the isolated extracellular vesicles (EV). b and c) Nanoparticle tracking analyses (NTA) data showing EV concentrations (particles/mL) (b) and size distribution (c). One-way ANOVA analysis was performed to determine statistical significance, *<0.05. d) Western blot image showing abundance of EV markers (CD81, CD63, CD9, and Flotillin) in EV isolated from blood samples obtained from screening controls (LNG-RADS II) or LNG-RADS IV individuals. e) Western blot images from triplicate samples showing CD9 and CD63 protein levels in EV extracted from LNG RADS II or confirmed cancer patients (IV^+^) plasma or disease-free LNG-RADS IV (IV^-^) individuals. Flotillin protein levels are used as loading controls. These experiments are shown in triplicates. (f, g) Relative quantification of CD9 and CD63 protein levels in Western blot shown in (e). Proteins levels are denoted as ratio of pixel intensities mean values quantified from each of the three replicates and normalized to Flotillin. (h) Ratio of CD9:CD63 protein band intensities presented in (e). Quantification of band intensities (pixels) was performed using ImageJ. Bars represent data as mean ± SD. P-values are derived from t-test. *p<0.05, **p<0.01, ***p<0.001.

LDCT has a false positive rate of 23-50% ^5,6,33^. Consistent with this, follow up tissue biopsies analyses revealed that 30% (6 out of 20) of the patients who were classified as Lung-RADS4 at baseline LDCT screening did not have lung cancer (Table 1; group column, asterisk). We asked whether the EV density and size characteristics can differentiate between true Lung-RADS4 patients with lung cancer and the rest of the patients with benign lung nodules without cancers (Lung-RADS2 plus the false-positive Lung-RADS4 patients). Although the trend of reduced EV density and larger EV size in true Lung-RADS4 compared to controls persisted, these differences were not statistically significant (data not shown).

We then asked whether EV membranes molecular profiles can differentiate between screening controls, over-diagnosed individuals, and confirmed cancer patients. We first examined the protein abundance of known EV membrane markers (the tetraspanins CD9, CD63, CD81, and the resident EV protein flotillin). While EV isolated from all Lung-RADS4 (initial diagnosis, including false-positive) patients and screening controls showed similar levels of CD81 and flotillin, CD63 and, to a lesser extent, CD9 were specifically enriched in Lung-RADS4 EV (CD63 high, CD9 moderate) compared to Lung-RADS2 screening controls (CD63 low and CD9 low) (Figure 1d). EV exclusively derived from confirmed NSCLC patients (IV^+^) were CD63 high and CD9 moderate) when compared to EV from false-positive patients (CD63 low and CD9 moderate) or to screening controls (II: CD63 low and CD9 low) (Figure 1e-g). Consistent with these findings, the ratio of CD9/CD63 levels distinguished confirmed cases from false-positive Lung-RADS4 or screening individuals [Figure 1h, (mean±SD): 0.37±0.1 versus 1.86±0.66 versus 5.08±1.48, respectively). Thus, EV CD9/CD63 expression ratio, but not EV physical characteristics, may help differentiate NSCLC patients from cancer-free high-risk individuals.

### Plasma Let-7b-5p, miR-184, and miR-22-3p levels differentiate NSCLC patients from high-risk individuals

Next, we used next generation sequencing (NGS) approaches to profile EV miRNA from Lung-RADS4 confirmed cancer patients or over-diagnosed Lung-RADS4 individuals or high-risk screening controls (Lung-RADS2). We included circulating plasma miRNA because combining multiple analytes from diverse biological sources have the potential to identify robust biomarkers. We identified 58 differentially expressed miRNAs, including miRNA widely known to be deregulated in cancers (Supplemental table 1). To identify a set of miRNAs that can robustly discriminate between NSCLC and cancer-free individuals, we prioritized miRNAs that were differentially expressed in at least two of the following comparisons: Lung-RADS2 versus Lung-RADS4; Lung-RADS4 false positive versus confirmed cancer patients; Lung-RADS2 combined with false-positive Lung-RADS4 patients versus confirmed cancer patients; any of the preceding groups versus patients who rapidly progressed (LDCT imaging and/or death shortly after sampling). We then focused on miRNAs showing significant performance (*P* and area under the curve/AUC values) in receiver operating characteristic (ROC) analyses for further examination. This approach led to the discovery of let-7b-5p, miR-184, and miR-22-3p as potential biomarkers for discriminating cancer patients from high-risk controls.

Let-7b-5p levels were elevated in EV from Lung-RADS4 patients compared to EV from screening patients alone (Figure 2a, e) or to screening patients plus false-positive Lung-RADS4 patients combined (Figure 2b, f). Also, miR-184 abundance was significantly reduced in Lung-RADS4 EV compared to either control group (excluding or including false-positive Lung-RADS4 patients) (Figure 2c, d g, h). Circulating miRNA analyses showed that miR-22-3p levels were reduced in Lung-RADS4 patients compared to either control group (Figure 2i, j, k, l). ROC curves showed area under the curve (AUC) values above 70% and significant p-values for all of the identified markers across comparisons (Figure 2e-h, 2k, l). The expression of these miRNAs was independently verified in quantitative polymerase chain reaction (qPCR) experiments using EV or plasma samples obtained from controls or false-positives or confirmed cancer patients (Figure 2n-p and supplemental figure 1a-c). Further, the expression of Let-7b-5p, miR-184, miR-22-3p showed no significant correlation with the gender, age, and pack-years smoking record of the participants (supplemental figure 1d and supplemental table 1), providing additional statistical rigor and indicating that the expression profile of these EV and circulating miRNAs is specifically associated with disease status.

**Figure 2.**
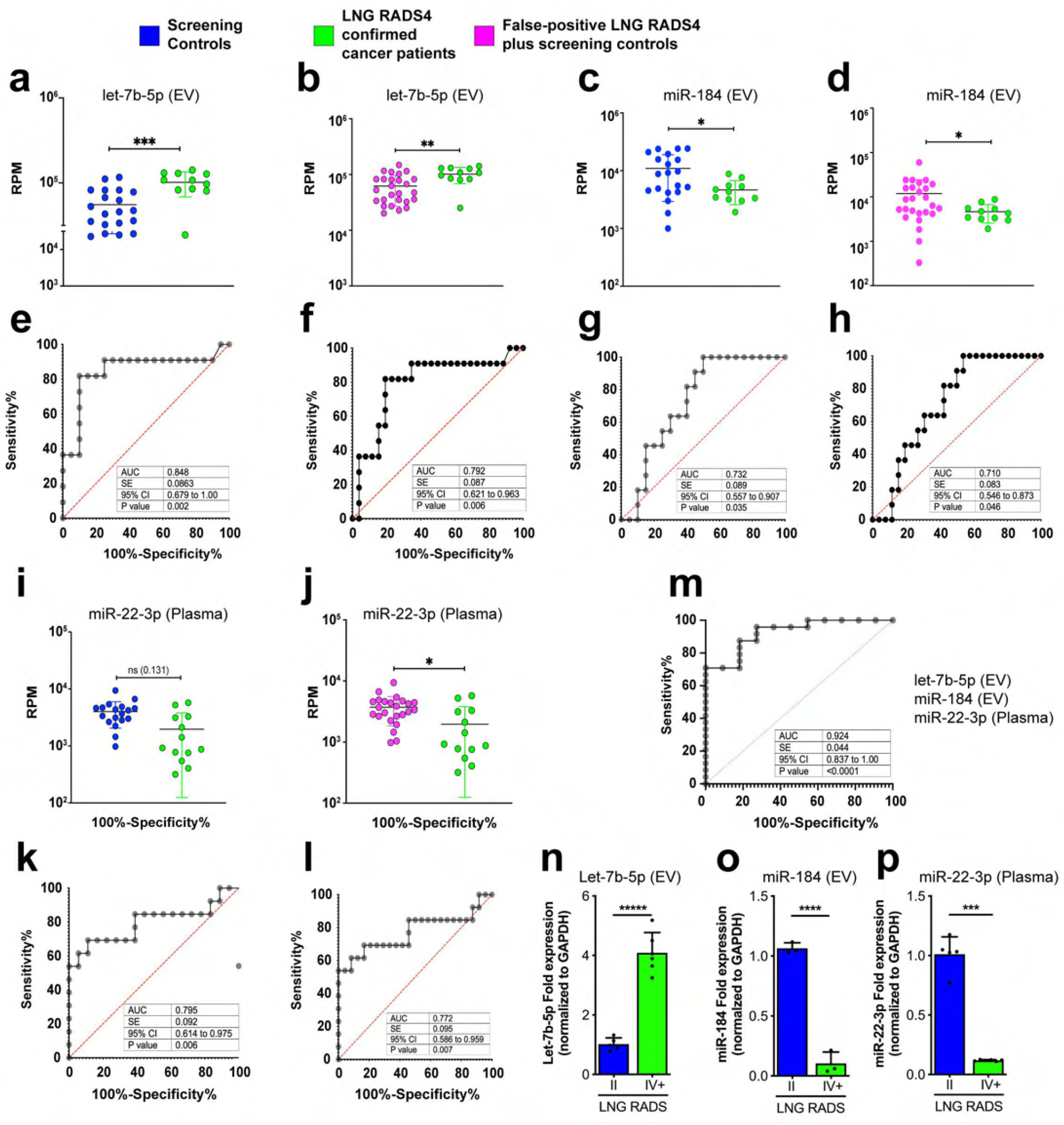
Identification of unique EV and circulating miRNA profiles Lung-RADS4 cancer patients. a-l) Next Generation Sequencing (NGS) of blood EV or circulating miRNA identified Let-7b-5p, miR-184, and miR-22-3p as differentially expressed in NSCLC patients compared to controls. Let-7b-5p (a, b) and mir-184 (c, d) are deregulated in EV and mir-22-3p in plasma (i, j). The relative abundance of the indicated miRNA in screening controls (blue) versus confirmed diseased patients only (green) or versus all lung-RADS2 plus false-positive lung-RADS4 (magenta) are shown in (a-d) and (I, j), respectively. P-values were Benjamini-Hochberg adjusted. *p<0.1, **p<0.05, ***p<0.01,****p<0.001. Note that 2-3 patient samples with undetectably low reads were excluded from the analysis. Corresponding Reads per million (RPM) were used to plot miRNA levels in confirmed cancer patients versus disease-free individuals and to perform ROC analysis shown in (e-h, k, l). EdgeR generalized linear models (GLM) were used to assess significance of miRNA regulations. Multiple logistic regression analysis was performed to determine the combined classification performance of the three miRNA biomarker candidates (m). Statistical significance of miRNA regulations was determined by EdgeR GLM, as indicated above. (n-p) Quantitative Polymerase Chain Reaction (qPCR) data showing expression fold changes (means) of let-7b-5p (n) or miRNA-184 (o) or miRNA-21-5p (n) or miRNA-22-3p (p) in EV samples obtained from confirmed cancer patients (IV+) versus screening controls (II). RNA samples were pooled from 14 cancer patients and 14 randomly selected screening individuals. Expression was normalized to *GAPDH*. Error bars denote SD values. *P* values are derived from student t-test analyses.

Furthermore, multiple logistic regression analyses of let-7b-5p, miR-184, and miR-22-3p showed a combined ROC AUC value of 92.4% (*p<0.0001*, Figure 2m and supplemental table 2), indicating remarkably high specificity and accuracy for these combined analytes to discriminate between cancer patients and high-risk Lung-RADS2 controls. Thus, the abundance of EV let-7b-5p, miR-184 combined with the levels of circulating miR-22-3p robustly discriminate between lung cancer patients and high-risk individuals.

### Let-7b-5p, miR-184, and miR-22-3p converge on treatment-resistance mechanisms

We considered the possibility that these EV and circulating plasma miRNAs (let-7b-5p, miR-184, and miR-22-3p) mediate cell-cell communication events that support NSCLC disease progression. First, we used the miRNA target proteins analysis platform MIRNET to identify experimentally validated let-7b-5p, miR-184, and miR-22-3p target proteins. We prioritized proteins that are targeted by at least two of the three miRNAs from experimental data (MIRNET miR2gene)^34^. This approach identified 43 proteins (Figure 3a and supplemental table 3), which were subsequently interrogated in Gene Ontology analyses using Kyoto Encyclopedia of Genes and Genomes (KEGG) or Reactome classifications to derive signaling pathways. Cancer was the most highly enriched KEGG term (Figure 3b, c), underscoring the robustness of our experimental pipeline and the relevance of these miRNAs to cancer disease. Interestingly, KEGG and Reactome signaling maps revealed that let-7b-5p, miR-184, and miR-22-3p converge on the activation of WNT and PI3K-AKT-mTOR signaling (Figure 3c, d), suggesting that circulating and EV miRNAs cooperatively regulate WNT and PI3K-AKT-mTOR activity in NSCLC. Activation of WNT or PI3K-AKT-mTOR signaling in NSCLC tissues is associated with aggressive and therapy resistant disease^35,36^. Considering that miR-184 and miR-22-3p are downregulated in cancer patients, this suggests that plasma from high-risk yet cancer-free individuals contain EV and circulating miRNAs that suppress WNT and the AKT signaling axis and that these mechanisms are restrained in NSCLC patients. Indeed, treatment of NSCLC cells (A549) with cancer patients EV elevated WNT and AKT signaling levels compared to the effect of EV from screening controls, as determined by β-catenin and phospho-AKT protein levels, respectively (Figure 3e, f).

**Figure 3.**
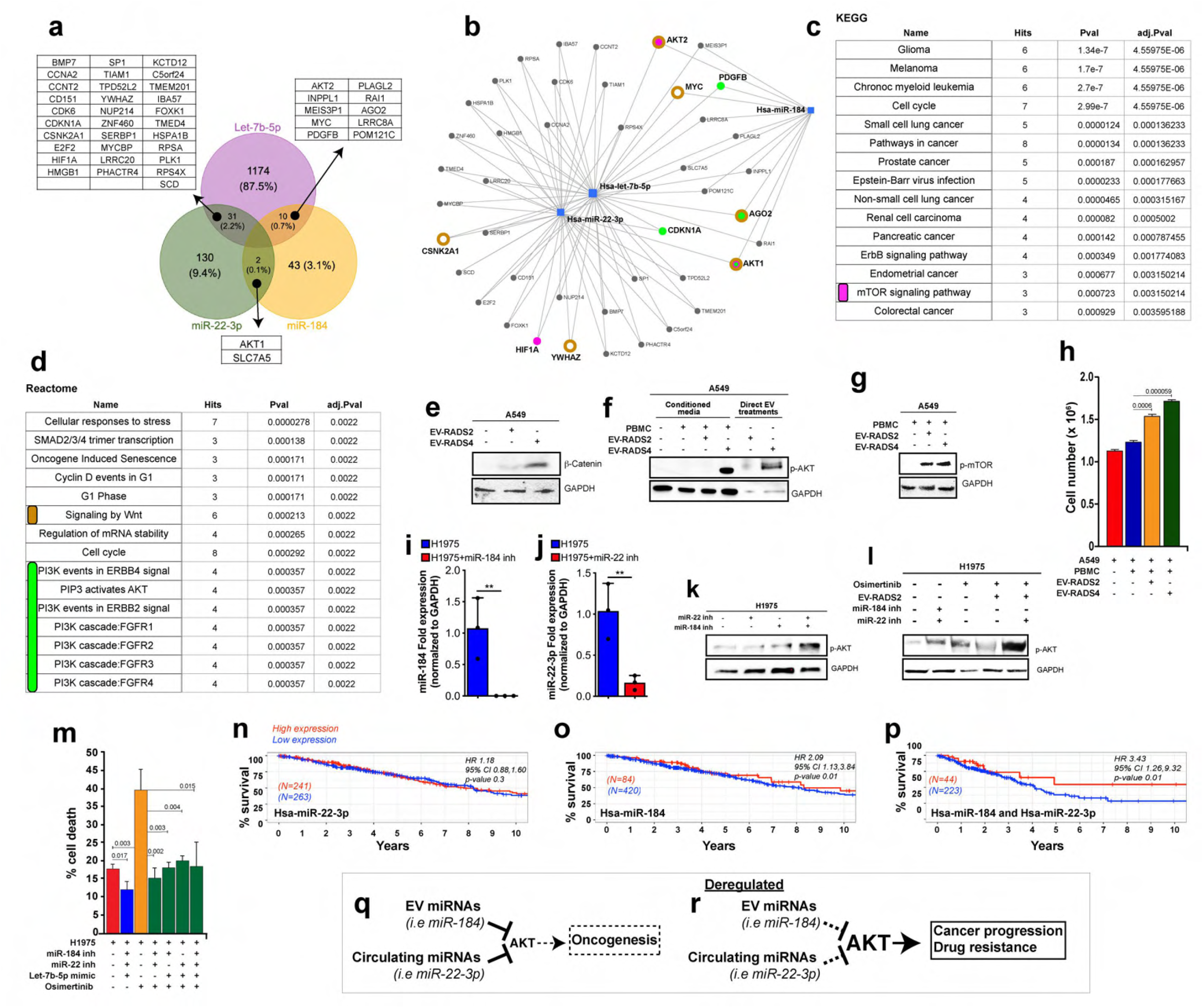
Let-7b-5p, miR-184, and miR-22-3p converge on AKT/mTOR and WNT/β-catenin therapy resistance pathways. a) Venn diagram showing the number unique of shared protein targets between the selected miRNAs. b) MIRNET star-network showing proteins that are targeted by at least two of the three miRNAs (blue squares). c, d) KEGG (c) or Reactome (d) classification analyses of the identified proteins show a convergence onto PI3K-AKT-mTOR (c, magenta label and d, green label) and WNT/β-catenin (d, brown label) signaling pathways. The underlying target genes are shown in corresponding color in “b”. e) Western blot image from A549 cells treated with equal quantity of EV from cancer patients or from high-risk screening controls. Blots were stained against β-catenin to detect WNT signaling levels or GAPDH as a loading control. f, g) Western blot images from A549 cells cultured in standard media or media conditioned with PBMC in the absence or presence of patients EV (f, lanes1-4). Also, A549 cells were treated directly with cancer patients or control EV (f, lanes 5 and 6). Blots were stained against phospho-AKT1 (f) or phospho-mTOR(g) or GAPDH as a loading control (f, g). A549 cell numbers from supernatant transfer experiments (f, lanes 1-4) are shown in “h” as average cell numbers from triplicate experiments. Error bars denote SD values. *P* values are derived from student t-test analyses. i, j) qPCR data showing mean expression fold changes of miR-184 (i) or miR-22-3p (j) in H1975 cells transfected with miR-184 (i) or miR-22-3p inhibitors (j) (blue bars) compared to untreated H1975 control cells (red bars). Expression was normalized to *GAPDH*. k) Western blot images from H1975 cells left untreated or treated with miR-22-3p or miR-184 inhibitors and blotted against phospho-AKT1 or GAPDH (loading control). l) Image of a Western blot from untreated H1975 cells or H1975 cells treated with equal portions of EV from screening controls (RADS-II) or with inhibitors against miR-22-3p and/or miR-184 inhibitors followed with Osimertinib (100nM) treatments. Western blots were stained against phospho-AKT or GAPDH (loading control). m) Graph showing the proportion of H1975 cell death across the indicated conditions using Trypan blue exclusion assays. These cell death assays were performed in triplicates and the results are shown as the average proportion (percentage) of dead cell across replicates for each treatment conditions. Error bars denote standard deviations and p-values were derived from t-tests. n-p) Survivorship comparison data using miR-184 and/or miR-22-3p expression data in TCGA-LUAD and the Bioconductor tool TCGA Biolinks RTCGA R packages. The “surv_cutpoint” function of the “survminer” R package was used to identify high versus low expressing patients’ samples for miR-22-3p (n) or miR-184 (o) or both (p) in Cox regression analyses. Survminer uses selected rank statistics to determine the optimal cut-point of a continuous variable in an unbiased manner. The related bioinformatics and statistics are presented in supplementary information 1. q, r) proposed model summarizing the role of mir-184/mir-22 in Osimertinib drug response. EV (mir-184) and circulating (mir-22) plasma miRNAs cooperatively target and modulate AKT activity. (q) Cancer-free high-risk individuals up-regulate EV mir-184 and circulating mir-22, keeping AKT levels generally low. In the context of genetic driver mutations, this low AKT activity delays oncogenesis. Similarly, cancer patients with high EV mir-184 and circulating mir-22-3p maintain AKT below an activity threshold required for AKT-mediated drug resistance, leading to positive drug response. However, AKT baseline activity is elevated in patients with low mir-184/mir-22 levels such that even a modest stimulation of AKT drives AKT above the drug-resistance activity threshold, leading to relapses and poor clinical response (r).

Further, the uptake of EV by immune cells results in paracrine signaling loops that ultimately accelerate disease progression via complex mechanisms^37–39^. We performed supernatant transfer experiments and asked whether cancer patients EV activate AKT/mTOR in A549 cells either directly or via immune cells. A549 cells were cultured in media conditioned by peripheral blood mononuclear cells (PBMC) left untreated or treated with EV either from controls or from cancer patients. Cancer patients EV/PBMC media dramatically stimulated phospho-AKT and phospho-mTOR levels in A549 cells, compared to controls (Figure 3f, g). Note that EV from the high-risk controls also stimulated mTOR signaling, possibly reflecting a NSCLC priming state (see discussion). Consistent with this AKT/mTOR activating potential, EV/PBMC conditioned media accelerated the growth of A549 cells (Figure 3h). Taken together, the above data argue that plasma EV act directly or via immune cells to activate AKT in NSCLC.

Next, we sought to determine whether let-7b-5p, miR-184, and miR-22-3p mediate the AKT activating effect of cancer EV and what implication this might have on NSCLC treatment outcomes. Activating mutations in Epidermal Growth Factor Receptor (EGFR) signaling represent one of the most known genetic alterations associated with NSCLC^40,41^. Patients harboring sensitizing EGFR mutations (exon 19 deletion and L858R) respond favorably to first- and second-generation Tyrosine Kinase Inhibitors/TKI (gefitinib, erlotinib, afatinib, and dacomitinib). However, patients acquire TKI-desensitizing EGFR mutations (T790M) and become resistant to these TKIs. Osimertinib, a third generation TKI selectively targets EGFR T790M and generates significant clinical benefits in EGFR T790M patients ^10,42–45^. Unfortunately, all patients ultimately develop resistance to Osimertinib because they acquire an Osimertinib-desensitizing mutation (C797S) or activate complex compensatory signals to resist drug-induced cell death and to promote cancer cell proliferation ^15,46–48^. Understanding the nature of these signals and how they are activated have the potential to inform new treatment strategies for re-sensitizing patients to existing TKIs.

We investigated a role for let-7b-5p, miR-184, and miR-22-3p in NSCLC response to Osimertinib using H1975 NSCLC cells, which harbor L858R and T790M EGFR mutations. First, H1975 cells were transfected with miR-184 and miR-22-3p inhibitors, mimicking their reduction in cancer patient plasma and assessed the effect of miR-184/miR-22-3p inhibition on AKT activity. The inhibitors reduced miR-184 and miR-22-3p levels in qPCR assays (Figure 3i and j, respectively) and cooperatively elevated pAKT levels in H1975 cells (Figure 3k, l).

In reciprocal experiments we found that Lung-RADS2 EV, which overexpress miR-184 (Figure 2c, o), inhibit Osimertinib-induced AKT activity in H1975 cells (Figure 3l). Importantly, inhibition of miR-22-3p in this setting was sufficient to dramatically unleash AKT (Figure 3l), further highlighting the cooperation between these two miRNAs in modulating AKT activity. Finally, we investigated the effect of miR-184/miR-22-3p inhibition on Osimertinib-induced cell death. Consistent with AKT stimulation, miR-184/miR-22-3p inhibition significantly suppressed Osimertinib-induced cell death (Figure 3m). Similar results were observed when miR-184/miR-22-3p co-inhibited cells were treated in the presence of a let-7b-5p mimic (Figure 3m).

AKT activation is associated with NSCLC resistance to TKI, leading to reduced patient survival time ^35,36^. Thus, we sought to determine to what extent reduced miR-184/miR-22-3p tumor expression correlates with reduced patient survival using the cancer genome atlas (TCGA) LUAD patient tumor miRNA expression and survival data. We did not detect any significant survival difference between patients whose tumors express low miR-22-3p (Figure 3n). However, patients with low miR-184 tumor expression had a significantly shorter survival time compared to patients with higher miR-184 tumor expression (Figure 3o, Hazard Ratio 2.09, 95% CI: 1.13, 3.84, p<0.018). Interestingly, patients with miR-184/miR-22-3p co-repressed tumors experienced even shorter survival time compared to patients with high miR-184/miR-22-3p tumor expression (Figure 3p, Hazard Ratio 3.43, 95% CI: 1.26, 9.32, p< 0.016). Note that the survivorship of miR-184/miR-22-3p tumor low patients is significantly lower than that of patients with tumor low miR-184 alone or miR-22-3p alone. This is consistent with AKT activation in miR-184/miR-22-3p co-inhibited NSCLC patients’ plasma and treatment resistance.

Thus, EV (let-7b-5p, miR-184) and circulating (miR-22-3p) plasma miRNA likely modulate NSCLC response to Osimertinib, highlighting a novel mechanism of resistance and suggesting that these biomarkers may assist in the selection of patients that will likely benefit from Osimertinib/AKT blockade combination treatments.

## Discussion

The implementation of LDCT in NSCLC screening has reduced patient deaths by ~20% ^5^. However, LDCT screening has a 23-50% false-positive rate ^5,6,33^, causing unnecessary exposure to radiation, costly and invasive follow-up studies for these mis-diagnosed patients. Additional strategies are needed to improve NSCLC screening accuracy.

Here we report the identification of a set of miRNAs (let-7-5p, miR-184 from EV and miR-22-3p from circulating miRNAs) that distinguishes NSCLC patients from high-risk controls. In addition, we found that the EV markers CD9, CD63, but not CD81 or flotillin, are differentially expressed in NSCLC patients compared to high-risk controls.

Some of the unique features of this study include the fact that reference and disease cohorts were controlled for risk profiles, reducing noise and elevating the relevance of the identified biomarkers in NSCLC. Also, circulating miRNA are gaining much interest as potential biomarkers in lung cancer ^30,31^. Different from others, our study integrates plasma and EV miRNAs to not only identify potential core molecular signatures for NSCLC risk management but to also highlight a plausible cooperation between these miRNAs in influencing patients’ drug response. Combining multiple analytes of diverse biological origins (EV surface markers, EV and plasma miRNAs) also maximizes robustness in diagnostics. Consistent with this, our biomarkers were able to differentiate confirmed NSCLC patients from LDCT false-positives, suggesting that addition of these non-invasive biomarkers to LDCT screening has the potential to improve diagnosis accuracy.

Confirmed NSCLC patients are stratified to diverse treatment options, including chemotherapy, immunotherapy, and targeted therapies based on histological and genetic mutations profiles obtained from tissue biopsies. Due to positional constraints, however, tissues biopsies often fail to capture the broader complexity of genetic driver mutations, leading to incomplete targeted therapy responses. The profiling of cancer-derived EV from patients’ plasma may provide deeper insights into the overall cancer mutational landscape of the tumor and thus better guide treatment decisions in the future. Indeed, pathway analyses of let-7b-5p, miR-184, and miR-22-3p target proteins revealed that these miRNAs converge on therapy resistance signals, including AKT. We propose that circulating and EV miRNAs functionally cooperate to modulate oncogenesis or patients’ clinical outcomes (Figure 3q, r). The AKT-suppressing miRNAs miR-184/miR-22-3p function as tumor suppressors and delay oncogenesis (Figure 3q). Cancers downregulate the expression of these miRNAs and/or reduce their systemic abundance via an unknown mechanism, leading to high baseline AKT activity and potentially resulting into accelerated cancer growth and drug resistance (Figure 3r). EV and plasma miRNAs may act directly or via tumor-interacting immune components to modulate tumor cell signaling and behavior. Mimicking the expression profile of miR-184 (EV) and miR-22-3p (plasma) in NSCLC patients’ blood using miRNA inhibitors cooperatively activated AKT and desensitized NSCLC cells (H1975) to Osimertinib. Thus, Let-7b-5p, miR-184, and miR-22-3p represent liquid biopsy biomarkers that will potentially assist with the identification of patients that will likely benefit from Osimertinib/AKT blockade combination treatments. AKT inhibition re-sensitizes TKI-resistant NSCLC cells to erlotinib and gefitinib^49^. A clinical trial evaluating the efficacy of combining Osimertinib with aspirin (an AKT inhibitor) in advanced NSCLC patients is pending (NCT04184921). Further, aberrant activation of AKT/mTOR and WNT/β-catenin signaling are associated with therapy resistance across different cancer types, suggesting broad translatability for these markers and their underlying mechanisms of action. Future studies will include determining to what extent blood levels of Let-7b-5p, miR-184, and miR-22-3p predict disease relapse in larger patient cohorts.

We noted that EV derived from high-risk control patients triggered mTOR activation in A549 cells. We cannot rule out the possibility that this EV-induced mTOR activation is specific to A549 cells, but it is likely that this effect reflects a priming state that precedes NSCLC onset in these high-risk patients.

Finally, the observation that circulating and EV miRNAs that suppress cancer cell survival and growth signals are upregulated in blood from high-risk cancer-free individuals compared to cancer patients also provides insights into why, despite having comparable NSCLC risk profiles, some individuals do not develop cancer, but others do.

### Ethical considerations

All blood samples were collected with participants’ written informed consent and in compliance with the University of Missouri ethical guidelines. The protocol was approved by the University of Missouri Ethics committee (IRB approval #2010166).

## Materials and methods

### Isolation of extracellular vesicles and circulating micro-RNA

Patients’ blood was double-centrifuged to remove platelets and the platelet-poor plasma was used for EV analysis. Purification of the EVs was done through size exclusion chromatography (SEC) using qEV70s single columns (ddIzon, Netherlands). These columns were filled with 150 μl of plasma and used based on recommendations from the manufacturer; fraction 8 through 11 were collected and pooled to obtain the fraction of purified EVs. The particle size and concentration for all EV samples were determined using nanoparticle tracking analysis (NTA).

Total RNA extraction was performed as previously described ^50^ using 200 μl of plasma or 200 μl of pooled EV fractions and the miRNeasy mini kit (Qiagen). Samples were thawed at room temperature followed by centrifugation at 12,000×g for 5 minutes at 4°C to remove any debris. Extraction was conducted by use of the miRNA easy Qiagen kit. For homogenization, 200 μL of plasma/EV suspension were mixed with 1000 μL Qiazol and 1 μL of a mix of 3 synthetic spike-in controls (Qiagen, Germany). After a 10-minute incubation at room temperature, 200 μL chloroform were added to the lysates followed by cooled centrifugation at 12,000×g for 15 minutes at 4°C. Precisely 650 μL of the upper aqueous phase were mixed with 7 μL glycogen (50 mg/mL) to enhance precipitation. Samples were transferred to a miRNeasy mini column, and RNA was precipitated with 750 μL ethanol followed by automated washing with RPE and RWT buffer in a QiaCube liquid handling robot. Finally, total RNA was eluted in 30 μL nuclease free water and stored at −80°C until further use.

### Electron microscopy

Exosomes were processed for negative staining as described in M. Rames, et al. ^51^. Briefly, 5 μL of purified EV sample was applied to a freshly glow discharged (Pelco Easiglow, Ted Pella Redding CA) carbon coated TEM grid (Electron Microscopy Sciences, Hatfield PA) over ice. Samples were washed three times with distilled water, incubated 2 min on 2% paraformaldehyde (Electron Microscopy Sciences), then incubated for 30 sec on 2% uranyl acetate (aqueous, Electron Microscopy Sciences) and back-blotted with filter paper (Whatman P1, Fisher Scientific) and allowed to dry. Images were collected on a JEOL JEM 1400 transmission electron microscope operated at 120V equipped with a Gatan Ultrascan 1000 CCD camera.

### Small RNA sequencing

Small RNA sequencing was performed as described previously ^52^. Equal volumes of total RNA (2 μL) were used for small RNA library preparation using the Clean Tag small RNA library preparation kit (TriLink Biotechnologies, US). that utilizes chemically modified adapters to prevent formation of adapter dimers ^53^. Adapter-ligated libraries were amplified using barcoded Illumina reverse primers in combination with the Illumina forward primer. A pool consisting of 40 plasma samples, and a second pool consisting of 40 EV samples was prepared by mixing samples at equimolar rates based on a DNA-1000 bioanalyzer results (Agilent, CA). The DNA library pool underwent size-selection (BluePippin, SageScience, US) to enrich for microRNAs with an insert size of 18-36 nt, corresponding to a library size of approximately 145 bp.

Sequencing was performed on an Illumina NextSeq 550 with 75 bp single end runs. Overall quality of the next-generation sequencing data was evaluated automatically and manually with FastQC v0.11.8 and MultiQC v1.7. Reads from all passing samples were adapter trimmed and quality filtered using Cutadapt v2.3 and filtered for a minimum length of 17nt. Mapping steps were performed with bowtie v1.2.2 and miRDeep2 v2.0.1.2, whereas reads were mapped first against the genomic reference GRCh38.p12 provided by Ensemble allowing for two mismatches and subsequently miRBase v22.1, filtered for miRNAs of hsa only, allowing for one mismatch. For a general RNA composition overview, non-miRNA mapped reads were mapped against RNAcentral and then assigned to various RNA species of interest.

Statistical analysis of preprocessed NGS data was done with R v3.6 and the packages pheatmap v1.0.12, pcaMethods v1.78 and genefilter v1.68. Differential expression analysis with edgeR v3.28 used the quasi-likelihood negative binomial generalized log-linear model (GLM) functions provided by the package. False discovery rate (FDR) correction was performed to adjust for multiple testing, and a cut-off of FDR < 5% was applied.

### Target network analysis

miRNA target network analyses and genes ontology enrichments analyses (KEGG and Reactome) were conducted using miRNet (www.mirnet.ca).

### Cell lines and cell culture

The human lung cancer cell line A549 were grown in Dulbecco’s Modified Eagle Medium (DMEM, 11965-092), with L-Glutamine, and high glucose supplemented with 10% FBS. Cells were grown in the nutrient medium as suggested by ATCC. Cells were incubated in a humidified incubator with 5% CO2 at 37°C. H1975 cells were grown in ATCC-formulated RPMI-1640 Medium (ATCC, Cat number 30-2001), with 10% FBS (fetal bovine serum) at 37 C, 5% CO2.

### Peripheral blood mononuclear cells (PBMC) isolation

10 mL of blood from healthy individual was collected into EDTA coated anti-coagulant vacutainer tubes (Cat #367899, BD Biosciences). Transferred onto 50 mL sterile 50 mL centrifuge tube and added equal volume with of ice cold DPBS pH 7.4 and gently mix by inversion. Using transfer pipette, carefully transferred diluted blood to sterile centrifuge tube containing 10 mL of Ficoll-Paque plus (Amersham #17144003,) then centrifuge at 2000 rpm for 25 min. after centrifuged from the fractionated phases, carefully collect the PBMCs fraction between ficol-paque and plasma layer. Collected PBMCs washed with ice cold PBS and centrifuge at 1700 rpm for 10 min. pellet re-suspended with 2 ml of Pharm Lyse lysing buffer (Biosciences #555899), mix well incubate at 37°C for 4 min and adjust volume with PBS to 50 mL and proceed with centrifuge at 1700 rpm for 10 min. PBMC pellet re suspend in PBS with 1×10^6^ cells/mL for further experimental purpose.

### EV/PBMC supernatant transfer experiments

Isolated human PBMCs were seeded and grown in 6 well culture dish with hybridoma-(SFM) serum free medium (Cat #12045076, Gibco). PBMCs were treated with patient EVs under serum free conditions for 24 hrs. After incubation cell free PBMC or EV/PBMC conditioned media were collected and used for culturing A549 cells. After 24 hours A549 cells were collected for cell counting or lysed for Western blotting.

### Western blotting

#### Extracellular vesicles (EV)

EV pellets were dissolved in 100 ul of modified lysis buffer (20 mM Tris-HCl pH-7.5, 150 mM NaCl, 1mM Na_2_EDTA, 1mM EGTA, 1%TritonX-100, 2.5 mM sodium pyrophosphate, 1 mM b-glycerophosphate, 1 mM Na3VO4, 1μg/mL leupeptin). Samples were treated with 10% glycerol, 1 M urea, 0.1% SDS, and loading buffer before SDS-PAGE. Gels were transferred onto PVDF membranes and stained with primary antibodies against CD-9 (1:1000, Millipore #CBL162,), CD-63 (1:1000, Molecular probes #A15712), CD-81 (1:1000, BioLegend #349561), and flotillin (1:1000, Santa Cruz #74566). Secondary antibodies were anti-mouse horseradish peroxidase (1:10000, Invitrogen #31430). Protein bands were detected using the Pierce ECL Chemiluminescence kit western (Thermo Fisher #32106) and the ChemiDoc™ Imaging System, Bio-Rad Laboratories Inc.

#### EV-treated A549 cells

A549 cells were cultured with similar loads of EV derived either from high-risk controls or confirmed lung cancer patients. Lysates were prepared in lysis buffer (20 mM Tris-HCl pH-7.5, 150 mM NaCl, 1 mM Na_2_ EDTA, 1 mM EGTA, 1%TritonX-100) and processed for Western blotting. Blots were stained against β-catenin to determine WNT signaling levels and GAPDH as a loading control (1:1000, Cell signaling #5174S). Secondary horseradish peroxidase (HRP) antibodies were obtained from Invitrogen. Pierce ECL Chemiluminescence kit (Thermo Fisher scientific #32106) and the ChemiDoc Imaging System (Bio-Rad) were used to detect protein bands.

A549 cells were washed with PBS and lysed in a lysis buffer (20 mM Tris-HCl pH-7.5, 150 mM, NaCl, 1 mM Na_2_EDTA, 1 mM EGTA, 1%Triton, 2.5 mM sodium pyrophosphate, 1 mM b-glycerophosphate, 1 mM Na3VO4, 1μg/ml leupeptin) supplemented with protease and a phosphatase inhibitor cocktail (Cell Signaling #9803S). Proteins were electrophoresed on SDS-PAGE using 4-20% Mini-PROTEAN® TGX™ precast gel (Biorad #456-1094,), transferred onto methanol pretreated PVDF membrane. PVDF membranes were probed overnight with mouse anti β-catenin (DSHB, #PY489), mouse anti-mTOR (1:1000, Santa Cruz, #517464) and mouse anti-GAPDH (DSHB, #2G7) at 4°C and after washing 3 times, membranes were incubated with anti-rabbit mouse horseradish peroxidase (HRP) (1:5000, #31460, Invitrogen). Primary or secondary antibodies were diluted in 5% BSA-TBST. Anti-mouse horseradish peroxidase (HRP) (1:5000, Invitrogen #31430,), anti-rabbit horseradish peroxidase (HRP) (1:5000, Invitrogen #31460,) used to develop respective blots. Membranes were developed using Pierce ECL western blotting substrate (Thermo Fisher scientific #32106,) and imaged on ChemiDoc™ MP Imaging System, Bio-Rad Laboratories Inc.

#### H1975 cells

H1975 cells were lysed in 1× RIPA buffer (Thermofisher #89900) containing 1× Halt™ Protease Inhibitor Cocktail (Thermofisher #78425). Proteins were separated on a 4-12% Bis-Tris gel (Invitrogen, Cat# NP0321) and transferred to PVDF membranes. Membranes were incubated with primary antibodies against Phospho-AKT (1:5000, Proteintech #66444-1-Ig,) or GAPDH (Sigma #G8795) overnight at 4°C. Secondary antibodies were purchased from Cell Signaling (Cat#7076). Blots were developed with Immobilon Western Chemiluminescence Kit (Millipore, #WBKLS0500).

miRNA Transfections and Osimertinib treatments miRNA stock was prepared by suspending in RNAse-free water. Cells were seeded such that they were 70-80% confluent at the time of transfection. Cells were allowed to adhere for 24 hrs and then transfected with 25 nM of miRNA inhibitors against hsa-miR184 (Sigma #HSTUD0282), hsa-miR22-3p (Sigma #HSTUD0393) or 100 nM of hsa-let7b-5p miRNA mimic (Sigma #HMI0007) using Lipofectamine 3000 (Invitrogen #L3000001). Osimertinib (100 nM) was added an hour after transfection to cells. Cells were allowed to incubate with transfection mix for 24 hrs at 37 °C, 5% CO2 and then washed with 1 × PBS and trypsinized to be used for cell count and Western blotting.

### cDNA synthesis and qRT-PCR

cDNA synthesis was carried out using the miRCURY LNA RT Kit (Qiagen #339340) from human purified EVs or plasma RNA (10 ng) according to manufacturer’s instructions. The synthesized CDNA diluted in nuclease free water and stored until further use at −80°C as per the kit instructions. Quantitative polymerase chain reaction was conducted using miRCURY LNA miRNA SYBR PCR kit (Qiagen #339345) as per manufacturer’s instruction on BioRad CFX96™ System in 96-well plates in 3-6 repeats. A two-step thermal cycling protocol i.e., 95°C for 2 min followed by 40 cycles at 95°C for 10 sec and 56°C for 60 sec, was used. A no-reverse transcriptase (NRT) and no-template control (NTC) were included in each reaction to check for primer specificity and any non-specific amplification. miRNA targets include miRNA-184 (Qiagen #YP00204601), miRNA-21-5p (Qiagen #YP00204230), Let-7b-5p (Qiagen #YP00204750), miRNA 22-3p (Qiagen #YP00204606). The expression levels of each miRNA target were normalized to calibrators U6-snRNA or GAPDH. Fold changed of miRNA was calculated by ΔΔCt and 2^−ΔΔCt^ method. ΔCt was calculated by subtracting the average of Ct values of calibrator from Ct values of target miRNA. ΔΔCt was computed by subtracting ΔCt of the screening control from ΔCt of RADS IV group.

### Patient Data Analysis

MicroRNA expression (miRNAseq) and clinical data from Lung adenocarcinoma (LUAD) were collected from the publicly accessible TCGA database using the Bioconductor tool TCGA Biolinks RTCGA R packages. The “surv_cutpoint” function of the “survminer” R package was used to identify high versus low expressing patients’ samples for survival analysis. Survminer uses selected rank statistics to determine the optimal cut-point of a continuous variable in an unbiased manner. Kaplan–Meier (KM) survival plots and related statistics were generated using the Survival R package. (Supplemental information 2)

## Authors contributions

C.Y.C conceived the study. A.G and C.Y.C wrote the manuscript. J.K was the IRB primary investigator and coordinated the acquisition of patient blood samples with Y.M. A.G and G.P performed miRNA and miRNA target analyses. G.P.V and Y.M processed patient blood samples. V.A isolated patients EV and performed qPCR analyses. G.P.V carried out qPCR analyses, biochemical and morphometric EV studies. B.D planned and performed all cell growth and cell death and Western blotting analyses. N.P performed TCGA miRNA expression patient survival analyses. C.Y.C and G.L examined the data. This work was funded by MU start-up funds to C.Y.C, MOLSAMP support to A.G, and NIH-IMSD support to G.P.

## Acknowledgment

We thank D. Porcini and R. Kannan for providing H1975 cells, E. King for assistance with developing the R code for TCGA studies. We also thank M. Hackl and A. Diendorfer (TAmiRNA, Austria) for assistance with NGS analyses.

## Data availability

All data will be made available upon reasonable request.

