## Supplemental Figure 1 for "Combining Plasma Extracellular Vesicle Let-7b-5p, miR-184 and Circulating miR-22-3p Levels for NSCLC Diagnosis and for Predicting Drug Resistance"

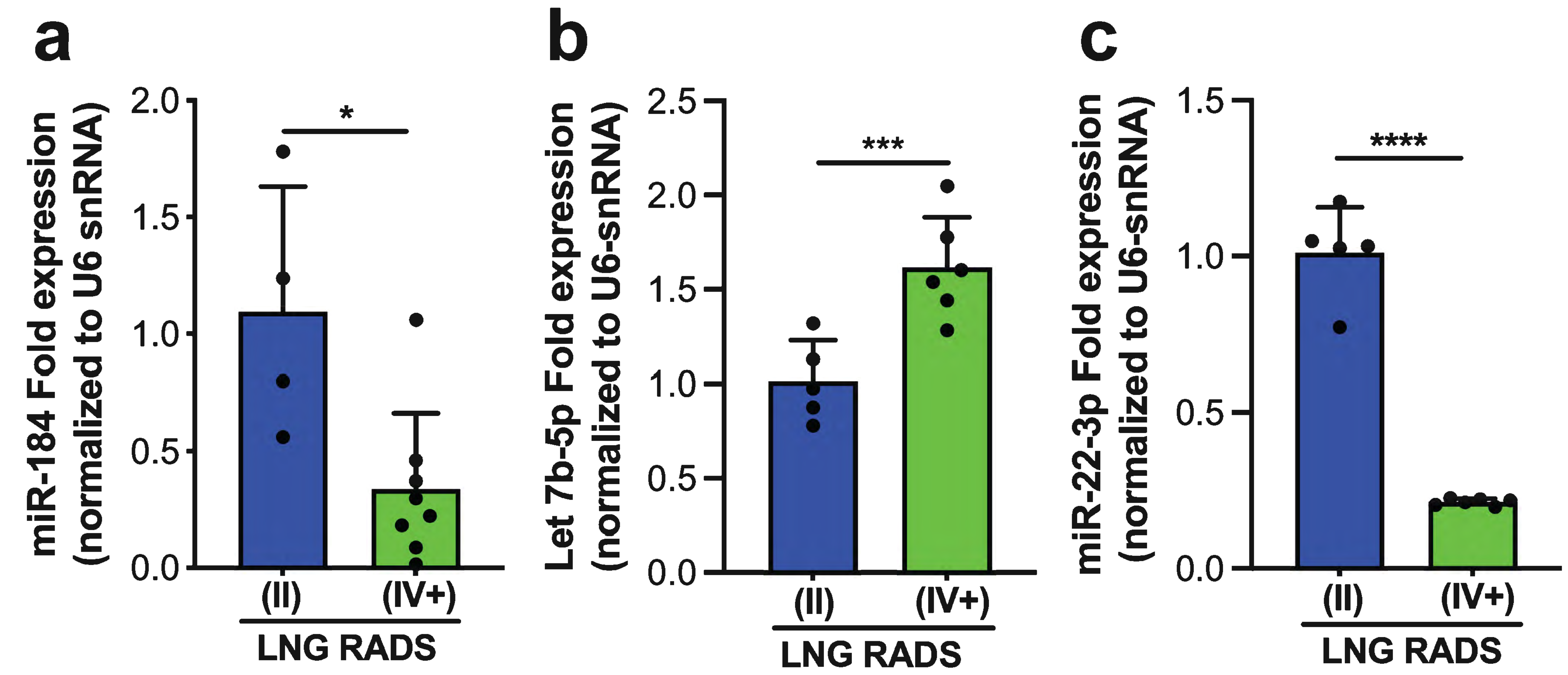

**d**

| Covariates analyses | | | | | | |
|---|---|---|---|---|---|---|
| **Continuous variables** | **miR** | **Estimate** | **Test** | **p.adj** | **p.adj.global** | |
| Age | plasma.miR -22-3p | -0.338 | spearman | 0.0658 | 0.1453 | |
| Age | ev.hsa -let-7b-5p | 0.3181 | spearman | 0.0969 | 0.1453 | |
| Age | ev.hsa -miR-184 | -0.1278 | spearman | 0.4381 | 0.6571 | |
| Smoking History (pack -year) | plasma.miR -22-3p | -0.0157 | spearman | 0.9234 | 0.9234 | |
| Smoking History (pack -year) | ev.hsa -let-7b-5p | -0.0683 | spearman | 0.6794 | 0.8153 | |
| Smoking History (pack -year) | ev.hsa -miR-184 | 0.1459 | spearman | 0.4381 | 0.6571 | |
| **category variable** | **groupA** | **groupB** | **miR** | **p.adj** | **test** | **p.adj.global** |
| Gender | Male | Female | plasma.miR -22-3p | 0.3466 | Wilcoxon test | 0.4159 |
| Gender | Male | Female | ev.hsa -let-7b-5p | 0.2919 | Wilcoxon test | 0.4159 |
| Gender | Male | Female | ev.hsa -miR-184 | 0.4427 | Wilcoxon test | 0.4427 |

**Supplementary Figure 1. Validation of miR-184, let7b-5p, and miR-22-3p disease-specific expression profile**

(a-c) qPCR analysis showing the mean of fold changes in the expression levels of miR-184 (a) or let-7b-5p (b) or miR-22-3p (c) in pooled samples from 14 confirmed cancer patients (IV+) versus 14 randomly selected individuals in the screening control group (II). Expression was normalized to the U6-snRNA gene. Error bars denote SD values. P values are derived from student t-test analyses. *p<0.05, **p<0.01, ***p<0.001, ****<0.0001.

(d) Sequencing covariates analyses for the expression of miRNA-184 or let-7b-5p or miR-22-3p. The related R-session information is presented in supplementary information 2
