## Supplemental Table 1 for "Combining Plasma Extracellular Vesicle Let-7b-5p, miR-184 and Circulating miR-22-3p Levels for NSCLC Diagnosis and for Predicting Drug Resistance"

| LEGEND | |
|---|---|
| A (N=20) | Screening control |
| B (N=20) | Initial RADS4 Diagnosis |
| B1 (N=14) | Confirmed cases |
| B2 (N=6) | LNG-RADS4-False Positive |
| C (N=5) | Disease progression (LDCT/Death) |

| Source | miRNA | logFC | logCPM | F | P-value | Comparisons | Implicated in cancer (PMIDs) |
|---|---|---|---|---|---|---|---|
| EV | hsa-miR-10b-5p | -0.593 | 17.225 | 6.986 | 0.012 | A vs B | 32236623, 31798622, 30853516 |
| EV | hsa-let-7b-5p | 0.611 | 16.706 | 4.538 | 0.039 | A vs B | 33283713, 29017393, 33733556, 32300359, 30995921, 25611389 |
| EV | hsa-miR-26a-5p | 0.545 | 15.337 | 3.966 | 0.053 | A vs B | 34313719, 34149922, 34141617, 33898304, 33841514, 33795765, 33761638, 33744438, 33553298 |
| EV | hsa-miR-184 | -1.79 | 13.17 | 10.43 | 0.00 | B1 vs B2 | 34328198, 34076278, 34055058, 34039407, 34000513, 33982217, 33816469, 33656787, 33354213 |
| EV | hsa-miR-92b-3p | -2.19 | 11.44 | 6.73 | 0.01 | B1 vs B2 | 34283053, 34136482, 34047469, 33984967, 33240410, 32864858, 32514152, 32256526, 31975504 |
| EV | hsa-miR-574-5p | -1.29 | 12.94 | 5.07 | 0.02 | B1 vs B2 | 34136389, 33989902, 33907569, 33869216, 33354213, 33317407, 33231826, 32832554, 32769887 |
| EV | hsa-miR-203a-3p | -3.14 | 14.36 | 4.65 | 0.05 | B1 vs B2 | 34316330, 34147083, 33664577, 33504224, 33080572, 32806571, 32708601, 3267869, 332672054 |
| EV | hsa-miR-122-5p | 3.899287 | 12.2346089 | 26.0619736 | 7.9417E-06 | C vs A | 34313924, 34053467 , 33992764, 33760201, 33752480, 33738672, 33583123, 33531550, 33505957 |
| EV | hsa-miR-200c-3p | 5.125889 | 9.91150971 | 20.6179926 | 1.2535E-05 | C vs A | 34315420, 34259142, 34257614, 34071861, 34067060, 33911091, 33866108, 33804458, 33174523 |
| EV | hsa-miR-31-5p | -5.44366 | 10.5156451 | 11.2580472 | 0.00105829 | C vs A | 34201353, 34082869, 34074322, 33867838, 33682215, 33531839, 33526040, 33510047, 33370717 |
| EV | hsa-miR-100-5p | -4.88756 | 10.1486038 | 9.14525193 | 0.00301642 | C vs A | 34218800, 34102608, 33948855, 33911091, 33504224, 33465676, 33335853, 32911741, 32767323 |
| EV | hsa-miR-148a-3p | 2.146505 | 11.325399 | 9.04796255 | 0.00315259 | C vs A | 34324740, 34288373, 34257725, 34098776, 34082012, 34021072, 34007215, 33959508, 33922968 |
| EV | hsa-miR-378a-3p | 2.840503 | 9.80565516 | 8.21489885 | 0.00484017 | C vs A | 34222341, 34113564, 33823894, 33766217, 33751805, 33634974, 33634097, 33578293, 33531456 |
| EV | hsa-miR-192-5p | 1.65028 | 12.3389158 | 7.85517362 | 0.00583524 | C vs A | 34104082, 34042256, 33806966, 33735626, 33708127, 33256802, 33024199, 32914665, 32626956 |
| EV | hsa-miR-363-3p | -3.7842 | 10.2131451 | 6.66152515 | 0.01094898 | C vs A | 34286520, 33950983, 33742335, 33640021, 33015780, 32779194, 32638620, 32561457, 32380791 |

| | | | | | | | |
|---|---|---|---|---|---|---|---|
| EV | hsa-miR-4454 | -3.97105 | 9.66192374 | 6.48518854 | 0.01206807 | C vs A | 34315950, 33003646, 32816024, 32422901, 32350057, 28571588, 28469417, 26576778 |
| EV | hsa-miR-21-5p | 1.196673 | 13.5532275 | 6.37311549 | 0.01277623 | C vs A | 34336825, 34316687, 34295456, 34288551, 34288231, 34254453, 34249905, 34201504, 34101749 |
| EV | hsa-miR-141-3p | 2.440423 | 9.90951629 | 6.09247612 | 0.01486065 | C vs A | 34136401, 34112774, 34086253, 33977869, 33866108, 33854632, 33845141, 33820555, 33784000 |
| EV | hsa-miR-27a-5p | -3.25124 | 9.42401734 | 4.82880742 | 0.02979586 | C vs A | 34201504, 34160887, 34073426, 34007020, 33931086, 33675923, 33465676, 33161496, 32910366 |
| EV | hsa-miR-4700-5p | -3.06506 | 9.34585606 | 4.72515738 | 0.03157331 | C vs A | NA |
| EV | hsa-miR-7155-5p | -3.40754 | 9.51460762 | 4.70024454 | 0.03201701 | C vs A | 30128020 |
| EV | hsa-miR-127-3p | -2.95369 | 9.32769432 | 4.49875803 | 0.03585731 | C vs A | 34295456, 33744851, 33732019, 33734890, 33468140, 33155194, 32742197, 32703147, 32538067 |
| EV | hsa-miR-29c-3p | -2.83713 | 9.3066489 | 4.11911272 | 0.04448431 | C vs A | 34199886, 34035375, 33887587, 33798549, 33569427, 33536761, 33391354, 33314669, 33261630 |
| EV | hsa-miR-155-5p | 1.38374 | 11.4926268 | 4.088312 | 0.04521116 | C vs A | 34295456, 34277059, 34201504, 34192708, 34190442, 34131104, 34103994, 34024195, 33999777 |
| Plasma | hsa-miR-184 | -2.12128 | 8.81691171 | 7.05539364 | 0.01251046 | A vs C | 34328198, 34076278, 34055058, 34039407, 34000513, 33982217, 33816469, 33656787, 33354213 |
| Plasma | hsa-miR-3976 | -4.91 | 7.33 | 14.11 | 0.001 | A vs B | 27573701, 25388097 |
| Plasma | hsa-miR-423-5p | 1.05 | 13.70 | 9.81 | 0.003 | A vs B | 34283074, 34048164, 33924142, 33833530, 33505907 |
| Plasma | hsa-miR-4259 | -2.22 | 5.51 | 8.83 | 0.004 | A vs B | NA |
| Plasma | hsa-miR-486-5p | 0.83 | 19.85 | 8.25 | 0.007 | A vs B | 34323695, 34199766, 34165171, 34094949, 33955505 |
| Plasma | hsa-miR-320a-3p | 0.76 | 13.51 | 7.55 | 0.007 | A vs B | 30634628, 29803922, 32814878 |
| Plasma | hsa-miR-185-5p | 1.55 | 6.51 | 7.50 | 0.008 | A vs B | 34280472, 34270733, 34205829, 34084747, 33962174, 33885171, 33858489, 33817236, 33734890 |
| Plasma | hsa-miR-598-3p | -1.80 | 5.69 | 7.04 | 0.010 | A vs B | 34069838, 27347136 |
| Plasma | hsa-miR-7-5p | -1.42 | 7.47 | 6.76 | 0.011 | A vs B | 33283713, 29017393, 33733556, 32300359, 30995921, 25611389 |
| Plasma | hsa-miR-223-3p | -1.59 | 6.46 | 6.25 | 0.015 | A vs B | 34320033, 34237309, 34205829, 34149922, 33986876, 33750388 |
| Plasma | hsa-miR-99a-5p | 1.70 | 5.44 | 5.99 | 0.017 | A vs B | 34200463, 34192880, 34038200, 33846533, 33575116 |
| Plasma | hsa-let-7b-5p | 0.66 | 14.03 | 5.42 | 0.023 | A vs B | 33283713, 29017393, 33733556, 32300359, 30995921, 25611389 |
| Plasma | hsa-miR-16-5p | -1.25 | 11.53 | 7.46 | 0.011 | B1 vs B2 | 34132358, 34101749, 33993193, 33975517, 33789312 |
| Plasma | hsa-miR-22-3p | -0.89 | 11.01 | 6.27 | 0.017 | B1 vs B2 | 34320711, 34239562, 34036905, 34019487, 33974352 |
| Plasma | hsa-miR-425-5p | -1.49 | 9.55 | 5.44 | 0.025 | B1 vs B2 | 34280472, 34204158, 33670246, 33633568, 33123581 |
| Plasma | hsa-miR-146a-5p | -0.65 | 10.02 | 3.16 | 0.083 | B1 vs B2 | 34293926, 34232111, 34073426, 34055487, 33948855 |
| Plasma | hsa-miR-92b-3p | -0.65 | 10.09 | 2.96 | 0.094 | B1 vs B2 | 34283053, 34136482, 34047469, 33984967, 33240410, 32864858, 32514152, 32256526, 31975504 |
| Plasma | hsa-miR-423-5p | -1.08 | 13.56 | 17.29 | 0.000 | AB2 vs B1 | 34283074, 34048164, 33924142, 33833530, 33505907 |
| Plasma | hsa-let-7b-5p | -0.66 | 13.87 | 10.28 | 0.002 | AB2 vs B1 | 33283713, 29017393, 33733556, 32300359, 30995921, 25611389 |

| | | | | | | | |
|---|---|---|---|---|---|---|---|
| Plasma | hsa-miR-22-3p | 0.70 | 11.48 | 8.45 | 0.006 | AB2 vs B1 | 34320711, 34239562, 34036905, 34019487, 33974352 |
| Plasma | hsa-miR-320a-3p | -0.62 | 13.44 | 7.48 | 0.009 | AB2 vs B1 | 30634628, 29803922, 32814878 |
| Plasma | hsa-miR-486-5p | -0.63 | 19.75 | 7.30 | 0.010 | AB2 vs B1 | 34323695, 34199766, 34165171, 34094949, 33955505 |
| Plasma | hsa-let-7i-5p | -0.43 | 12.21 | 6.14 | 0.017 | AB2 vs B1 | 34009009, 33806966, 33775245, 33537093, 32799221 |
| Plasma | hsa-miR-122-5p | 1.48 | 12.06 | 5.02 | 0.031 | AB2 vs B1 | 34313924, 34053467 , 33992764, 33760201, 33752480, 33738672, 33583123, 33531550, 33505957 |
| Plasma | hsa-miR-192-5p | 3.46382 | 12.1063752 | 29.5907537 | 1.8992E-05 | A vs C | 34104082, 34042256, 33806966, 33735626, 33708127, 33256802, 33024199, 32914665, 32626956 |
| Plasma | hsa-miR-148a-3p | 3.23502 | 11.7392659 | 29.3569297 | 1.9988E-05 | A vs C | 34324740, 34288373, 34257725, 34098776, 34082012, 34021072, 34007215, 33959508, 33922968 |
| Plasma | hsa-miR-122-5p | 3.331167 | 12.9334566 | 18.0407132 | 0.00033661 | A vs C | 34313924, 34053467 , 33992764, 33760201, 33752480, 33738672, 33583123, 33531550, 33505957 |
| Plasma | hsa-miR-423-5p | 1.295447 | 13.5276312 | 10.2355684 | 0.00323341 | A vs C | 34283074, 34048164, 33924142, 33833530, 33505907 |
| Plasma | hsa-miR-320a-3p | 1.145576 | 13.5353561 | 9.29090428 | 0.00475998 | A vs C | 30634628, 29803922, 32814878 |
| Plasma | hsa-miR-486-5p | 0.819955 | 19.6941712 | 6.78175629 | 0.01416047 | A vs C | 34323695, 34199766, 34165171, 34094949, 33955505 |
| Plasma | hsa-miR-451a | -0.94834 | 12.7512907 | 6.05582553 | 0.01980453 | A vs C | 34345270, 34333815, 34257614, 34199766, 34183927 |
| Plasma | hsa-miR-142-5p | -0.71891 | 10.968385 | 5.82535735 | 0.02207851 | A vs C | 34323174, 34321149, 34178135, 34163033, 34040437 |
| Plasma | hsa-miR-21-5p | 0.633327 | 11.8878778 | 4.97194941 | 0.03335729 | A vs C | 34336825, 34316687, 34295456, 34288551, 34288231, 34254453, 34249905, 34201504, 34101749 |
| Plasma | hsa-miR-143-3p | 1.231633 | 8.70671219 | 4.29172087 | 0.04695037 | A vs C | 34335889, 34322386, 34279157, 34202934, 34085707 |
| Plasma | hsa-let-7c-5p | -1.71626 | 11.7255483 | 4.32640405 | 0.04952009 | A vs C | 34308851, 34026328, 33910598, 33461652, 33376529 |

**Supplementary table 1. List of differentially expressed miRNAs from NGS analyses**

Highlighted rows denote prioritized miRNAs that are significantly deregulated in at least two of the following groups comparisons: Lung-RADS2 versus Lung-RADS4; Lung-RADS4 false positive versus confirmed cancer patients; Lung-RADS2 combined with false-positive Lung-RADS4 patients versus confirmed cancer patients; any of the preceding groups versus patients who rapidly progressed (LDCT imaging). The last column shows PMID of prior reports implicating the corresponding miRNA in cancer biology.
