## Supplemental Table 2 for "Combining Plasma Extracellular Vesicle Let-7b-5p, miR-184 and Circulating miR-22-3p Levels for NSCLC Diagnosis and for Predicting Drug Resistance"

| miRNA | Accession | Target | Target ID | Experiment | Literature |
|---|---|---|---|---|---|
| hsa-let-7b-5p | MIMAT0000063 | ABCF1 | 23 | CLASH | 23622248 |
| hsa-let-7b-5p | MIMAT0000063 | ABL1 | 25 | CLASH | 23622248 |
| hsa-let-7b-5p | MIMAT0000063 | ACACA | 31 | CLASH//Proteomics | 18668040\|23622248 |
| hsa-let-7b-5p | MIMAT0000063 | ASIC1 | 41 | CLASH | 23622248 |
| hsa-let-7b-5p | MIMAT0000063 | ACPP | 55 | Proteomics | 18668040 |
| hsa-let-7b-5p | MIMAT0000063 | ACTA1 | 58 | PAR-CLIP | 21572407 |
| hsa-let-7b-5p | MIMAT0000063 | ACTB | 60 | CLASH | 23622248 |
| hsa-let-7b-5p | MIMAT0000063 | ACTG1 | 71 | CLASH//Luciferase reporter assay//Reporter assay | 15131085\|23622248 |
| hsa-let-7b-5p | MIMAT0000063 | ACTN4 | 81 | CLASH | 23622248 |
| hsa-let-7b-5p | MIMAT0000063 | ACVR1 | 90 | Luciferase reporter assay//Microarray//qRT-PCR | 22995917 |
| hsa-let-7b-5p | MIMAT0000063 | ADCY1 | 107 | CLASH | 23622248 |
| hsa-let-7b-5p | MIMAT0000063 | ADH5 | 128 | HITS-CLIP//PAR-CLIP | 23313552\|26701625\|27292025 |
| hsa-let-7b-5p | MIMAT0000063 | AGL | 178 | Proteomics | 18668040 |
| hsa-let-7b-5p | MIMAT0000063 | JAG1 | 182 | CLASH | 23622248 |
| hsa-let-7b-5p | MIMAT0000063 | AHR | 196 | PAR-CLIP | 23592263 |
| hsa-let-7b-5p | MIMAT0000063 | AK4 | 205 | PAR-CLIP | 23446348\|20371350 |
| hsa-let-7b-5p | MIMAT0000063 | AKT2 | 208 | Luciferase reporter assay// qRT-PCR//Western blot | 25288334 |
| hsa-let-7b-5p | MIMAT0000063 | AMD1 | 262 | HITS-CLIP | 23313552 |
| hsa-let-7b-5p | MIMAT0000063 | AMPH | 273 | CLASH | 23622248 |
| hsa-let-7b-5p | MIMAT0000063 | SLC25A4 | 291 | CLASH | 23622248 |
| hsa-let-7b-5p | MIMAT0000063 | BIRC5 | 332 | CLASH | 23622248 |
| hsa-let-7b-5p | MIMAT0000063 | APRT | 353 | Proteomics | 18668040 |
| hsa-let-7b-5p | MIMAT0000063 | AQP6 | 363 | PAR-CLIP | 27292025 |
| hsa-let-7b-5p | MIMAT0000063 | AR | 367 | CLASH | 23622248 |
| hsa-let-7b-5p | MIMAT0000063 | ARCN1 | 372 | Proteomics | 18668040 |
| hsa-let-7b-5p | MIMAT0000063 | RHOB | 388 | Proteomics//pSILAC | 18668040 |
| hsa-let-7b-5p | MIMAT0000063 | RHOG | 391 | Proteomics//pSILAC | 18668040 |
| hsa-let-7b-5p | MIMAT0000063 | ASNA1 | 439 | CLASH | 23622248 |

| | | | | | |
|---|---|---|---|---|---|
| hsa-let-7b-5p | MIMAT0000063 | ASPA | 443 | CLASH | 23622248 |
| hsa-let-7b-5p | MIMAT0000063 | ATOX1 | 475 | CLASH | 23622248 |
| hsa-let-7b-5p | MIMAT0000063 | ATP1A1 | 476 | CLASH | 23622248 |
| hsa-let-7b-5p | MIMAT0000063 | ATP2A2 | 488 | CLASH//Proteomics | 18668040\|23622248 |
| hsa-let-7b-5p | MIMAT0000063 | ALDH7A1 | 501 | CLASH | 23622248 |
| hsa-let-7b-5p | MIMAT0000063 | ATP6V0A1 | 535 | Proteomics//pSILAC | 18668040 |
| hsa-let-7b-5p | MIMAT0000063 | AUP1 | 550 | CLASH | 23622248 |
| hsa-let-7b-5p | MIMAT0000063 | BACH1 | 571 | PAR-CLIP | 20371350 |
| hsa-let-7b-5p | MIMAT0000063 | BCAT1 | 586 | CLASH | 23622248 |
| hsa-let-7b-5p | MIMAT0000063 | CCND1 | 595 | Annexin V-FITC//immunoblot//Immunoblot//Immunofluorescence//Luciferase reporter assay//Microarray//Northern blot//PAR-CLIP//qRT-PCR//Reporter assay//Western blot | 18379589\|20133835\|23806108\|23592263\|26701625 |
| hsa-let-7b-5p | MIMAT0000063 | BCL7A | 605 | Microarray//qRT-PCR | 18026111 |
| hsa-let-7b-5p | MIMAT0000063 | BFSP1 | 631 | CLASH | 23622248 |
| hsa-let-7b-5p | MIMAT0000063 | BGLAP | 632 | Proteomics | 18668040 |
| hsa-let-7b-5p | MIMAT0000063 | PRDM1 | 639 | Immunohistochemistry//Luciferase reporter assay//qRT-PCR//Western blot | 20651244 |
| hsa-let-7b-5p | MIMAT0000063 | BMP7 | 655 | CLASH | 23622248 |
| hsa-let-7b-5p | MIMAT0000063 | POLR3D | 661 | HITS-CLIP | 23313552 |
| hsa-let-7b-5p | MIMAT0000063 | BNIP3L | 665 | CLASH | 23622248 |
| hsa-let-7b-5p | MIMAT0000063 | VPS51 | 738 | Proteomics | 18668040 |
| hsa-let-7b-5p | MIMAT0000063 | CA12 | 771 | Proteomics | 18668040 |
| hsa-let-7b-5p | MIMAT0000063 | CALU | 813 | PAR-CLIP | 20371350 |
| hsa-let-7b-5p | MIMAT0000063 | CAPG | 822 | Proteomics//pSILAC | 18668040 |
| hsa-let-7b-5p | MIMAT0000063 | CBFB | 865 | Microarray | 17699775 |
| hsa-let-7b-5p | MIMAT0000063 | CCNA2 | 890 | Immunoblot//Immunofluorescence//Luciferase reporter assay//qRT-PCR | 18379589 |
| hsa-let-7b-5p | MIMAT0000063 | CCNB1 | 891 | CLASH | 23622248 |

| | | | | | |
|---|---|---|---|---|---|
| hsa-let-7b-5p | MIMAT0000063 | CCND2 | 894 | Immunohistochemistry//Luciferase reporter assay//qRT-PCR//QRTPCR//Western blot | 17699775\|17942906\|23482325 |
| hsa-let-7b-5p | MIMAT0000063 | CCND3 | 896 | CLASH | 23622248 |
| hsa-let-7b-5p | MIMAT0000063 | CCNF | 899 | CLASH//Microarray | 17699775\|23622248 |
| hsa-let-7b-5p | MIMAT0000063 | CCNG1 | 900 | CLASH | 23622248 |
| hsa-let-7b-5p | MIMAT0000063 | CCNT2 | 905 | PAR-CLIP | 21572407 |
| hsa-let-7b-5p | MIMAT0000063 | ENTPD6 | 955 | CLASH | 23622248 |
| hsa-let-7b-5p | MIMAT0000063 | CD59 | 966 | PAR-CLIP | 23446348\|20371350 |
| hsa-let-7b-5p | MIMAT0000063 | CD81 | 975 | CLASH | 23622248 |
| hsa-let-7b-5p | MIMAT0000063 | CD151 | 977 | CLASH | 23622248 |
| hsa-let-7b-5p | MIMAT0000063 | CDC25A | 993 | Immunohistochemistry//Luciferase reporter assay//Microarray//qRT-PCR//Western blot | 19966857\|17699775 |
| hsa-let-7b-5p | MIMAT0000063 | CDC34 | 997 | Immunoblot//Luciferase reporter assay//Microarray//qRT-PCR//Western blot | 19126550\|17699775\|21252116 |
| hsa-let-7b-5p | MIMAT0000063 | CDK6 | 1021 | Luciferase reporter assay//Microarray//Western blot | 17699775 |
| hsa-let-7b-5p | MIMAT0000063 | CDKN1A | 1026 | Immunoblot//Microarray//PAR-CLIP//qRT-PCR | 21572407\|25578966 |
| hsa-let-7b-5p | MIMAT0000063 | CDKN1B | 1027 | Immunoblot//Microarray//qRT-PCR | 25578966 |
| hsa-let-7b-5p | MIMAT0000063 | CENPB | 1059 | Proteomics | 18668040 |
| hsa-let-7b-5p | MIMAT0000063 | CHD1 | 1105 | Proteomics | 18668040 |
| hsa-let-7b-5p | MIMAT0000063 | CHD3 | 1107 | CLASH//Proteomics | 18668040\|23622248 |
| hsa-let-7b-5p | MIMAT0000063 | CHD4 | 1108 | CLASH//Proteomics | 18668040\|23622248 |
| hsa-let-7b-5p | MIMAT0000063 | CKB | 1152 | CLASH | 23622248 |
| hsa-let-7b-5p | MIMAT0000063 | CKS2 | 1164 | CLASH | 23622248 |
| hsa-let-7b-5p | MIMAT0000063 | AP1S1 | 1174 | PAR-CLIP | 23592263\|26701625 |
| hsa-let-7b-5p | MIMAT0000063 | COL3A1 | 1281 | Luciferase reporter assay//Microarray//qRT-PCR | 22995917 |
| hsa-let-7b-5p | MIMAT0000063 | COL8A1 | 1295 | PAR-CLIP | 23592263 |

| | | | | | |
|---|---|---|---|---|---|
| hsa-let-7b-5p | MIMAT0000063 | COX6B1 | 1340 | HITS-CLIP | 23706177 |
| hsa-let-7b-5p | MIMAT0000063 | COX7B | 1349 | Proteomics | 18668040 |
| hsa-let-7b-5p | MIMAT0000063 | CRKL | 1399 | Proteomics | 18668040 |
| hsa-let-7b-5p | MIMAT0000063 | CRX | 1406 | PAR-CLIP | 26701625 |
| hsa-let-7b-5p | MIMAT0000063 | CRY2 | 1408 | PAR-CLIP | 23592263\|26701625 |
| hsa-let-7b-5p | MIMAT0000063 | CS | 1431 | Proteomics | 18668040 |
| hsa-let-7b-5p | MIMAT0000063 | CSNK1D | 1453 | Proteomics//pSILAC | 18668040 |
| hsa-let-7b-5p | MIMAT0000063 | CSNK2A1 | 1457 | CLASH | 23622248 |
| hsa-let-7b-5p | MIMAT0000063 | CTBP2 | 1488 | CLASH | 23622248 |
| hsa-let-7b-5p | MIMAT0000063 | CTPS1 | 1503 | HITS-CLIP//Proteomics | 18668040\|23313552 |
| hsa-let-7b-5p | MIMAT0000063 | CUX1 | 1523 | CLASH | 23622248 |
| hsa-let-7b-5p | MIMAT0000063 | CYP1A2 | 1544 | CLASH | 23622248 |
| hsa-let-7b-5p | MIMAT0000063 | CYP2J2 | 1573 | Luciferase reporter assay//qRT-PCR//Western blot | 22761738 |
| hsa-let-7b-5p | MIMAT0000063 | DCTD | 1635 | CLASH | 23622248 |
| hsa-let-7b-5p | MIMAT0000063 | DHX9 | 1660 | CLASH | 23622248 |
| hsa-let-7b-5p | MIMAT0000063 | DDX10 | 1662 | Proteomics | 18668040 |
| hsa-let-7b-5p | MIMAT0000063 | DFFA | 1676 | CLASH | 23622248 |
| hsa-let-7b-5p | MIMAT0000063 | TIMM8A | 1678 | Proteomics | 18668040 |
| hsa-let-7b-5p | MIMAT0000063 | DIAPH1 | 1729 | CLASH//Proteomics | 18668040\|23622248 |
| hsa-let-7b-5p | MIMAT0000063 | DLAT | 1737 | Proteomics | 18668040 |
| hsa-let-7b-5p | MIMAT0000063 | DMD | 1756 | Microarray | 17699775 |
| hsa-let-7b-5p | MIMAT0000063 | DNA2 | 1763 | PAR-CLIP | 23446348 |
| hsa-let-7b-5p | MIMAT0000063 | DNAH9 | 1770 | HITS-CLIP | 23706177 |
| hsa-let-7b-5p | MIMAT0000063 | DYNC1H1 | 1778 | CLASH | 23622248 |
| hsa-let-7b-5p | MIMAT0000063 | DRG2 | 1819 | Proteomics | 18668040 |
| hsa-let-7b-5p | MIMAT0000063 | ARID3A | 1820 | Microarray//PAR-CLIP | 17699775\|23592263 |
| hsa-let-7b-5p | MIMAT0000063 | RCAN1 | 1827 | Microarray | 17699775 |
| hsa-let-7b-5p | MIMAT0000063 | DSG2 | 1829 | Proteomics | 18668040 |
| hsa-let-7b-5p | MIMAT0000063 | DSP | 1832 | CLASH//Proteomics//pSILAC | 18668040\|23622248 |
| hsa-let-7b-5p | MIMAT0000063 | DUSP1 | 1843 | PAR-CLIP | 21572407 |
| hsa-let-7b-5p | MIMAT0000063 | DVL3 | 1857 | PAR-CLIP | 23446348 |
| hsa-let-7b-5p | MIMAT0000063 | E2F2 | 1870 | Immunofluorescence//Luciferase reporter assay//qRT-PCR//Western blot | 27520092 |

| | | | | | |
|---|---|---|---|---|---|
| hsa-let-7b-5p | MIMAT0000063 | E2F3 | 1871 | CLASH | 23622248 |
| hsa-let-7b-5p | MIMAT0000063 | E2F5 | 1875 | Microarray | 17699775 |
| hsa-let-7b-5p | MIMAT0000063 | E2F6 | 1876 | Microarray//PAR-CLIP | 17699775\|20371350 |
| hsa-let-7b-5p | MIMAT0000063 | EDN1 | 1906 | PAR-CLIP | 23592263 |
| hsa-let-7b-5p | MIMAT0000063 | EEF1A1 | 1915 | CLASH | 23622248 |
| hsa-let-7b-5p | MIMAT0000063 | EEF2 | 1938 | CLASH | 23622248 |
| hsa-let-7b-5p | MIMAT0000063 | EIF4A1 | 1973 | Proteomics | 18668040 |
| hsa-let-7b-5p | MIMAT0000063 | EIF4A2 | 1974 | Proteomics | 18668040 |
| hsa-let-7b-5p | MIMAT0000063 | EIF4G2 | 1982 | PAR-CLIP//Proteomics | 18668040\|23592263\|21572407\|27292025 |
| hsa-let-7b-5p | MIMAT0000063 | ELK4 | 2005 | CLASH | 23622248 |
| hsa-let-7b-5p | MIMAT0000063 | ENG | 2022 | Proteomics | 18668040 |
| hsa-let-7b-5p | MIMAT0000063 | EP300 | 2033 | CLASH | 23622248 |
| hsa-let-7b-5p | MIMAT0000063 | EPHA4 | 2043 | PAR-CLIP | 21572407 |
| hsa-let-7b-5p | MIMAT0000063 | ERCC1 | 2067 | CLASH | 23622248 |
| hsa-let-7b-5p | MIMAT0000063 | ETFA | 2108 | CLASH | 23622248 |
| hsa-let-7b-5p | MIMAT0000063 | EZH2 | 2146 | qRT-PCR//Western blot | 25611389 |
| hsa-let-7b-5p | MIMAT0000063 | F2 | 2147 | Proteomics | 18668040 |
| hsa-let-7b-5p | MIMAT0000063 | FANCD2 | 2177 | CLASH | 23622248 |
| hsa-let-7b-5p | MIMAT0000063 | ACSL1 | 2180 | Proteomics | 18668040 |
| hsa-let-7b-5p | MIMAT0000063 | FEN1 | 2237 | CLASH | 23622248 |
| hsa-let-7b-5p | MIMAT0000063 | FLII | 2314 | CLASH | 23622248 |
| hsa-let-7b-5p | MIMAT0000063 | FLNA | 2316 | CLASH | 23622248 |
| hsa-let-7b-5p | MIMAT0000063 | FMO4 | 2329 | HITS-CLIP | 23824327 |
| hsa-let-7b-5p | MIMAT0000063 | FPR1 | 2357 | HITS-CLIP | 23313552 |
| hsa-let-7b-5p | MIMAT0000063 | FXN | 2395 | PAR-CLIP | 23592263 |
| hsa-let-7b-5p | MIMAT0000063 | GABPB1 | 2553 | HITS-CLIP//PAR-CLIP | 23592263\|23313552 |
| hsa-let-7b-5p | MIMAT0000063 | GALNT2 | 2590 | Proteomics | 18668040 |
| hsa-let-7b-5p | MIMAT0000063 | GAPDH | 2597 | CLASH | 23622248 |
| hsa-let-7b-5p | MIMAT0000063 | GATA6 | 2627 | CLASH | 23622248 |
| hsa-let-7b-5p | MIMAT0000063 | GATM | 2628 | PAR-CLIP | 26701625 |
| hsa-let-7b-5p | MIMAT0000063 | NR6A1 | 2649 | PAR-CLIP | 23592263 |
| hsa-let-7b-5p | MIMAT0000063 | GLB1 | 2720 | Proteomics | 18668040 |
| hsa-let-7b-5p | MIMAT0000063 | GLO1 | 2739 | PAR-CLIP//Proteomics | 18668040\|23592263 |
| hsa-let-7b-5p | MIMAT0000063 | GNAS | 2778 | CLASH | 23622248 |
| hsa-let-7b-5p | MIMAT0000063 | GNB1 | 2782 | CLASH | 23622248 |
| hsa-let-7b-5p | MIMAT0000063 | GNG5 | 2787 | PAR-CLIP | 20371350\|27292025 |
| hsa-let-7b-5p | MIMAT0000063 | GOLGA4 | 2803 | PAR-CLIP | 20371350 |
| hsa-let-7b-5p | MIMAT0000063 | GPI | 2821 | CLASH | 23622248 |

| | | | | | |
|---|---|---|---|---|---|
| hsa-let-7b-5p | MIMAT0000063 | GPM6B | 2824 | CLASH | 23622248 |
| hsa-let-7b-5p | MIMAT0000063 | GPX7 | 2882 | Microarray | 17699775 |
| hsa-let-7b-5p | MIMAT0000063 | GSK3A | 2931 | CLASH | 23622248 |
| hsa-let-7b-5p | MIMAT0000063 | GSPT1 | 2935 | Proteomics | 18668040 |
| hsa-let-7b-5p | MIMAT0000063 | GSR | 2936 | CLASH//Proteomics | 18668040\|23622248 |
| hsa-let-7b-5p | MIMAT0000063 | GTF2I | 2969 | Microarray | 17699775 |
| hsa-let-7b-5p | MIMAT0000063 | GTF3C1 | 2975 | CLASH | 23622248 |
| hsa-let-7b-5p | MIMAT0000063 | GYG1 | 2992 | CLASH | 23622248 |
| hsa-let-7b-5p | MIMAT0000063 | GYS1 | 2997 | Proteomics//pSILAC | 18668040 |
| hsa-let-7b-5p | MIMAT0000063 | GYS2 | 2998 | Proteomics | 18668040 |
| hsa-let-7b-5p | MIMAT0000063 | HIST1H1C | 3006 | CLASH | 23622248 |
| hsa-let-7b-5p | MIMAT0000063 | HIST1H2BD | 3017 | PAR-CLIP | 24398324\|23446348 |
| hsa-let-7b-5p | MIMAT0000063 | HADHA | 3030 | CLASH | 23622248 |
| hsa-let-7b-5p | MIMAT0000063 | HARS | 3035 | CLASH | 23622248 |
| hsa-let-7b-5p | MIMAT0000063 | HCFC1 | 3054 | CLASH | 23622248 |
| hsa-let-7b-5p | MIMAT0000063 | HTT | 3064 | CLASH | 23622248 |
| hsa-let-7b-5p | MIMAT0000063 | HELLS | 3070 | Proteomics | 18668040 |
| hsa-let-7b-5p | MIMAT0000063 | HIF1A | 3091 | CLASH | 23622248 |
| hsa-let-7b-5p | MIMAT0000063 | HK1 | 3098 | CLASH | 23622248 |
| hsa-let-7b-5p | MIMAT0000063 | HMGB1 | 3146 | CLASH | 23622248 |
| hsa-let-7b-5p | MIMAT0000063 | HMGCS1 | 3157 | CLASH | 23622248 |
| hsa-let-7b-5p | MIMAT0000063 | HMGA1 | 3159 | CLASH//HITS-CLIP//Luciferase reporter assay//Microarray//PAR-CLIP//Proteomics//pSILAC//qRT-PCR//Western blot | 18668040\|23622248\|23798998\|23592263\|23313552 |
| hsa-let-7b-5p | MIMAT0000063 | HNRNPF | 3185 | CLASH | 23622248 |
| hsa-let-7b-5p | MIMAT0000063 | HNRNPL | 3191 | CLASH | 23622248 |
| hsa-let-7b-5p | MIMAT0000063 | HOXD11 | 3237 | CLASH | 23622248 |
| hsa-let-7b-5p | MIMAT0000063 | HRAS | 3265 | Immunoblot//qRT-PCR//Western blot | 21252116 |
| hsa-let-7b-5p | MIMAT0000063 | AGFG2 | 3268 | CLASH | 23622248 |
| hsa-let-7b-5p | MIMAT0000063 | HES1 | 3280 | CLASH | 23622248 |
| hsa-let-7b-5p | MIMAT0000063 | HSF2 | 3298 | CLASH | 23622248 |
| hsa-let-7b-5p | MIMAT0000063 | HSPA1B | 3304 | CLASH | 23622248 |
| hsa-let-7b-5p | MIMAT0000063 | HSPA8 | 3312 | CLASH | 23622248 |
| hsa-let-7b-5p | MIMAT0000063 | HSP90AA1 | 3320 | CLASH | 23622248 |

| | | | | | |
|---|---|---|---|---|---|
| hsa-let-7b-5p | MIMAT0000063 | IGSF3 | 3321 | Proteomics | 18668040 |
| hsa-let-7b-5p | MIMAT0000063 | IDI1 | 3422 | Proteomics | 18668040 |
| hsa-let-7b-5p | MIMAT0000063 | IFNB1 | 3456 | ELISA//Luciferase reporter assay//qRT-PCR | 20130213 |
| hsa-let-7b-5p | MIMAT0000063 | IFRD1 | 3475 | Proteomics//pSILAC | 18668040 |
| hsa-let-7b-5p | MIMAT0000063 | IGF1R | 3480 | Immunohistochemistry//Luciferase reporter assay//PAR-CLIP//QRTPCR//Western blot | 23482325\|23592263\|24398324\|23446348\|21572407 |
| hsa-let-7b-5p | MIMAT0000063 | IGHMBP2 | 3508 | CLASH | 23622248 |
| hsa-let-7b-5p | MIMAT0000063 | RBPJ | 3516 | CLASH | 23622248 |
| hsa-let-7b-5p | MIMAT0000063 | IL6R | 3570 | PAR-CLIP | 27292025 |
| hsa-let-7b-5p | MIMAT0000063 | CXCL8 | 3576 | PAR-CLIP | 26701625 |
| hsa-let-7b-5p | MIMAT0000063 | IMPDH1 | 3614 | Proteomics | 18668040 |
| hsa-let-7b-5p | MIMAT0000063 | IMPDH2 | 3615 | Proteomics | 18668040 |
| hsa-let-7b-5p | MIMAT0000063 | INPPL1 | 3636 | Proteomics | 18668040 |
| hsa-let-7b-5p | MIMAT0000063 | ITGA3 | 3675 | HITS-CLIP | 23706177\|23313552 |
| hsa-let-7b-5p | MIMAT0000063 | ITGB5 | 3693 | CLASH | 23622248 |
| hsa-let-7b-5p | MIMAT0000063 | KCNC4 | 3749 | CLASH | 23622248 |
| hsa-let-7b-5p | MIMAT0000063 | KIF2A | 3796 | Proteomics | 18668040 |
| hsa-let-7b-5p | MIMAT0000063 | KIFC1 | 3833 | CLASH | 23622248 |
| hsa-let-7b-5p | MIMAT0000063 | KPNA5 | 3841 | HITS-CLIP//PAR-CLIP | 24398324\|23446348\|21572407\|20371350 |
| hsa-let-7b-5p | MIMAT0000063 | TNPO1 | 3842 | CLASH | 23622248 |
| hsa-let-7b-5p | MIMAT0000063 | RPSA | 3921 | CLASH | 23622248 |
| hsa-let-7b-5p | MIMAT0000063 | LBR | 3930 | Proteomics | 18668040 |
| hsa-let-7b-5p | MIMAT0000063 | LDLR | 3949 | Proteomics | 18668040 |
| hsa-let-7b-5p | MIMAT0000063 | LPL | 4023 | Proteomics | 18668040 |
| hsa-let-7b-5p | MIMAT0000063 | LTA4H | 4048 | CLASH | 23622248 |
| hsa-let-7b-5p | MIMAT0000063 | LYN | 4067 | PAR-CLIP | 23592263 |
| hsa-let-7b-5p | MIMAT0000063 | MAB21L1 | 4081 | CLASH | 23622248 |
| hsa-let-7b-5p | MIMAT0000063 | MXD1 | 4084 | PAR-CLIP | 21572407 |
| hsa-let-7b-5p | MIMAT0000063 | MAGEA3 | 4102 | PAR-CLIP | 27292025 |
| hsa-let-7b-5p | MIMAT0000063 | MAGEA6 | 4105 | PAR-CLIP | 27292025 |
| hsa-let-7b-5p | MIMAT0000063 | MAGEA12 | 4111 | PAR-CLIP | 27292025 |
| hsa-let-7b-5p | MIMAT0000063 | MAP4 | 4134 | CLASH | 23622248 |
| hsa-let-7b-5p | MIMAT0000063 | MARS | 4141 | Proteomics | 18668040 |

| | | | | | |
|---|---|---|---|---|---|
| hsa-let-7b-5p | MIMAT0000063 | MCM4 | 4173 | CLASH | 23622248 |
| hsa-let-7b-5p | MIMAT0000063 | MCM7 | 4176 | CLASH | 23622248 |
| hsa-let-7b-5p | MIMAT0000063 | MDM4 | 4194 | CLASH//PAR-CLIP | 23622248\|23592263\|24398324 |
| hsa-let-7b-5p | MIMAT0000063 | MEF2C | 4208 | CLASH | 23622248 |
| hsa-let-7b-5p | MIMAT0000063 | MEF2D | 4209 | PAR-CLIP | 26701625 |
| hsa-let-7b-5p | MIMAT0000063 | MEIS3P1 | 4213 | PAR-CLIP | 23592263 |
| hsa-let-7b-5p | MIMAT0000063 | CD99 | 4267 | CLASH | 23622248 |
| hsa-let-7b-5p | MIMAT0000063 | MIPEP | 4285 | Proteomics | 18668040 |
| hsa-let-7b-5p | MIMAT0000063 | MLLT1 | 4298 | Proteomics//pSILAC | 18668040 |
| hsa-let-7b-5p | MIMAT0000063 | MOV10 | 4343 | CLASH | 23622248 |
| hsa-let-7b-5p | MIMAT0000063 | MPG | 4350 | Proteomics | 18668040 |
| hsa-let-7b-5p | MIMAT0000063 | ABCC1 | 4363 | Proteomics | 18668040 |
| hsa-let-7b-5p | MIMAT0000063 | MSI1 | 4440 | CLASH | 23622248 |
| hsa-let-7b-5p | MIMAT0000063 | MSN | 4478 | CLASH | 23622248 |
| hsa-let-7b-5p | MIMAT0000063 | ATP6 | 4508 | CLASH | 23622248 |
| hsa-let-7b-5p | MIMAT0000063 | COX1 | 4512 | CLASH | 23622248 |
| hsa-let-7b-5p | MIMAT0000063 | COX2 | 4513 | CLASH | 23622248 |
| hsa-let-7b-5p | MIMAT0000063 | COX3 | 4514 | CLASH | 23622248 |
| hsa-let-7b-5p | MIMAT0000063 | ND1 | 4535 | CLASH | 23622248 |
| hsa-let-7b-5p | MIMAT0000063 | ND2 | 4536 | CLASH | 23622248 |
| hsa-let-7b-5p | MIMAT0000063 | ND4 | 4538 | CLASH | 23622248 |
| hsa-let-7b-5p | MIMAT0000063 | ND4L | 4539 | CLASH | 23622248 |
| hsa-let-7b-5p | MIMAT0000063 | ND5 | 4540 | CLASH | 23622248 |
| hsa-let-7b-5p | MIMAT0000063 | MTRR | 4552 | Proteomics//pSILAC | 18668040 |
| hsa-let-7b-5p | MIMAT0000063 | MYC | 4609 | CLASH//TRAP | 23622248\|24510096 |
| hsa-let-7b-5p | MIMAT0000063 | MYO1C | 4641 | CLASH | 23622248 |
| hsa-let-7b-5p | MIMAT0000063 | MYO1E | 4643 | Proteomics | 18668040 |
| hsa-let-7b-5p | MIMAT0000063 | NACA | 4666 | CLASH | 23622248 |
| hsa-let-7b-5p | MIMAT0000063 | NAP1L1 | 4673 | HITS-CLIP | 19536157 |
| hsa-let-7b-5p | MIMAT0000063 | NDUFA10 | 4705 | CLASH | 23622248 |
| hsa-let-7b-5p | MIMAT0000063 | NEDD4 | 4734 | Proteomics//pSILAC | 18668040 |
| hsa-let-7b-5p | MIMAT0000063 | NFATC1 | 4772 | CLASH | 23622248 |
| hsa-let-7b-5p | MIMAT0000063 | NFATC3 | 4775 | CLASH | 23622248 |
| hsa-let-7b-5p | MIMAT0000063 | NFKBIA | 4792 | CLASH | 23622248 |
| hsa-let-7b-5p | MIMAT0000063 | NME4 | 4833 | CLASH | 23622248 |
| hsa-let-7b-5p | MIMAT0000063 | CNOT2 | 4848 | CLASH | 23622248 |
| hsa-let-7b-5p | MIMAT0000063 | SLC11A2 | 4891 | HITS-CLIP//PAR-CLIP | 23592263\|23313552 |

| | | | | | |
|---|---|---|---|---|---|
| hsa-let-7b-5p | MIMAT0000063 | NRAS | 4893 | Immunohistochemistry//Luciferase reporter assay//Proteomics//qRT-PCR | 17699775\|19966857\|18668040 |
| hsa-let-7b-5p | MIMAT0000063 | NRDC | 4898 | CLASH | 23622248 |
| hsa-let-7b-5p | MIMAT0000063 | NUCB2 | 4925 | PAR-CLIP | 23592263 |
| hsa-let-7b-5p | MIMAT0000063 | NVL | 4931 | Proteomics | 18668040 |
| hsa-let-7b-5p | MIMAT0000063 | OPRL1 | 4987 | PAR-CLIP | 22012620 |
| hsa-let-7b-5p | MIMAT0000063 | ORC4 | 5000 | Proteomics | 18668040 |
| hsa-let-7b-5p | MIMAT0000063 | OXA1L | 5018 | Proteomics | 18668040 |
| hsa-let-7b-5p | MIMAT0000063 | PA2G4 | 5036 | CLASH | 23622248 |
| hsa-let-7b-5p | MIMAT0000063 | PAFAH1B3 | 5050 | Proteomics | 18668040 |
| hsa-let-7b-5p | MIMAT0000063 | PAFAH2 | 5051 | HITS-CLIP | 19536157 |
| hsa-let-7b-5p | MIMAT0000063 | PAK1 | 5058 | CLASH | 23622248 |
| hsa-let-7b-5p | MIMAT0000063 | PAX3 | 5077 | CLASH | 23622248 |
| hsa-let-7b-5p | MIMAT0000063 | PBX2 | 5089 | PAR-CLIP | 26701625 |
| hsa-let-7b-5p | MIMAT0000063 | PCBP2 | 5094 | CLASH | 23622248 |
| hsa-let-7b-5p | MIMAT0000063 | PCCB | 5096 | Proteomics | 18668040 |
| hsa-let-7b-5p | MIMAT0000063 | PCYT1A | 5130 | Proteomics | 18668040 |
| hsa-let-7b-5p | MIMAT0000063 | PDGFB | 5155 | PAR-CLIP | 27292025 |
| hsa-let-7b-5p | MIMAT0000063 | PDGFRA | 5156 | Luciferase reporter assay//Microarray//qRT-PCR | 17942906\|22995917 |
| hsa-let-7b-5p | MIMAT0000063 | PDK1 | 5163 | Proteomics | 18668040 |
| hsa-let-7b-5p | MIMAT0000063 | PER1 | 5187 | CLASH | 23622248 |
| hsa-let-7b-5p | MIMAT0000063 | PEX6 | 5190 | CLASH | 23622248 |
| hsa-let-7b-5p | MIMAT0000063 | PFKM | 5213 | CLASH | 23622248 |
| hsa-let-7b-5p | MIMAT0000063 | PFN1 | 5216 | CLASH | 23622248 |
| hsa-let-7b-5p | MIMAT0000063 | PGM1 | 5236 | CLASH | 23622248 |
| hsa-let-7b-5p | MIMAT0000063 | PGM3 | 5238 | Proteomics | 18668040 |
| hsa-let-7b-5p | MIMAT0000063 | PHKA1 | 5255 | CLASH | 23622248 |
| hsa-let-7b-5p | MIMAT0000063 | PLAGL2 | 5326 | Microarray//PAR-CLIP | 17699775\|23592263\|26701625 |
| hsa-let-7b-5p | MIMAT0000063 | PLCB3 | 5331 | Proteomics | 18668040 |
| hsa-let-7b-5p | MIMAT0000063 | PLCG2 | 5336 | PAR-CLIP | 24398324\|23446348 |
| hsa-let-7b-5p | MIMAT0000063 | PLK1 | 5347 | CLASH | 23622248 |
| hsa-let-7b-5p | MIMAT0000063 | PMAIP1 | 5366 | PAR-CLIP | 23592263 |
| hsa-let-7b-5p | MIMAT0000063 | Sep-04 | 5414 | Proteomics | 18668040 |
| hsa-let-7b-5p | MIMAT0000063 | POLD2 | 5425 | Proteomics//pSILAC | 18668040 |
| hsa-let-7b-5p | MIMAT0000063 | POLR2A | 5430 | CLASH | 23622248 |

| | | | | | |
|---|---|---|---|---|---|
| hsa-let-7b-5p | MIMAT0000063 | POLR2C | 5432 | Proteomics//pSILAC | 18668040 |
| hsa-let-7b-5p | MIMAT0000063 | POLR2D | 5433 | PAR-CLIP | 23592263\|21572407\|26701625 |
| hsa-let-7b-5p | MIMAT0000063 | POLR2H | 5437 | Proteomics | 18668040 |
| hsa-let-7b-5p | MIMAT0000063 | POLR2L | 5441 | Proteomics | 18668040 |
| hsa-let-7b-5p | MIMAT0000063 | PPID | 5481 | Proteomics | 18668040 |
| hsa-let-7b-5p | MIMAT0000063 | PPM1G | 5496 | Proteomics | 18668040 |
| hsa-let-7b-5p | MIMAT0000063 | PPP1R7 | 5510 | Proteomics//pSILAC | 18668040 |
| hsa-let-7b-5p | MIMAT0000063 | PPP2R2A | 5520 | CLASH//PAR-CLIP | 23622248\|23446348 |
| hsa-let-7b-5p | MIMAT0000063 | PPP2R5E | 5529 | CLASH | 23622248 |
| hsa-let-7b-5p | MIMAT0000063 | PRIM1 | 5557 | Proteomics//pSILAC | 18668040 |
| hsa-let-7b-5p | MIMAT0000063 | PRIM2 | 5558 | HITS-CLIP | 23706177 |
| hsa-let-7b-5p | MIMAT0000063 | PRKAA2 | 5563 | CLASH | 23622248 |
| hsa-let-7b-5p | MIMAT0000063 | PRKAR2A | 5576 | Proteomics | 18668040 |
| hsa-let-7b-5p | MIMAT0000063 | PRKD1 | 5587 | CLASH | 23622248 |
| hsa-let-7b-5p | MIMAT0000063 | MAPK1 | 5594 | CLASH | 23622248 |
| hsa-let-7b-5p | MIMAT0000063 | MAPK6 | 5597 | PAR-CLIP | 21572407 |
| hsa-let-7b-5p | MIMAT0000063 | MAP2K2 | 5605 | Proteomics | 18668040 |
| hsa-let-7b-5p | MIMAT0000063 | MAP2K7 | 5609 | PAR-CLIP | 23592263\|26701625 |
| hsa-let-7b-5p | MIMAT0000063 | PRPS1 | 5631 | Proteomics | 18668040 |
| hsa-let-7b-5p | MIMAT0000063 | PSMD9 | 5715 | Proteomics | 18668040 |
| hsa-let-7b-5p | MIMAT0000063 | PTGFRN | 5738 | CLASH | 23622248 |
| hsa-let-7b-5p | MIMAT0000063 | PTGS2 | 5743 | Proteomics//pSILAC | 18668040 |
| hsa-let-7b-5p | MIMAT0000063 | PVR | 5817 | Proteomics | 18668040 |
| hsa-let-7b-5p | MIMAT0000063 | PCYT2 | 5833 | CLASH | 23622248 |
| hsa-let-7b-5p | MIMAT0000063 | QARS | 5859 | Proteomics | 18668040 |
| hsa-let-7b-5p | MIMAT0000063 | QDPR | 5860 | PAR-CLIP | 23592263 |
| hsa-let-7b-5p | MIMAT0000063 | RALB | 5899 | CLASH | 23622248 |
| hsa-let-7b-5p | MIMAT0000063 | RBBP6 | 5930 | CLASH | 23622248 |
| hsa-let-7b-5p | MIMAT0000063 | RDX | 5962 | PAR-CLIP//Proteomics | 18668040\|23446348\|21572407\|26701625 |
| hsa-let-7b-5p | MIMAT0000063 | DPF2 | 5977 | Proteomics | 18668040 |
| hsa-let-7b-5p | MIMAT0000063 | RFC2 | 5982 | HITS-CLIP | 23706177 |
| hsa-let-7b-5p | MIMAT0000063 | RHD | 6007 | HITS-CLIP//PAR-CLIP | 23592263\|23706177\|23313552 |
| hsa-let-7b-5p | MIMAT0000063 | BRD2 | 6046 | CLASH | 23622248 |
| hsa-let-7b-5p | MIMAT0000063 | RPL12 | 6136 | CLASH | 23622248 |
| hsa-let-7b-5p | MIMAT0000063 | RPL18 | 6141 | CLASH | 23622248 |
| hsa-let-7b-5p | MIMAT0000063 | RPL18A | 6142 | CLASH | 23622248 |
| hsa-let-7b-5p | MIMAT0000063 | MRPL12 | 6182 | PAR-CLIP | 23592263 |
| hsa-let-7b-5p | MIMAT0000063 | RPS4X | 6191 | CLASH | 23622248 |

| | | | | | |
|---|---|---|---|---|---|
| hsa-let-7b-5p | MIMAT0000063 | RPS24 | 6229 | CLASH | 23622248 |
| hsa-let-7b-5p | MIMAT0000063 | RRAD | 6236 | PAR-CLIP | 22012620 |
| hsa-let-7b-5p | MIMAT0000063 | RRBP1 | 6238 | CLASH | 23622248 |
| hsa-let-7b-5p | MIMAT0000063 | RRM1 | 6240 | PAR-CLIP | 23592263 |
| hsa-let-7b-5p | MIMAT0000063 | RRM2 | 6241 | Microarray//PAR-CLIP//Proteomics | 17699775\|18668040\|21572407 |
| hsa-let-7b-5p | MIMAT0000063 | RXRB | 6257 | CLASH | 23622248 |
| hsa-let-7b-5p | MIMAT0000063 | SAFB | 6294 | CLASH | 23622248 |
| hsa-let-7b-5p | MIMAT0000063 | SALL2 | 6297 | CLASH | 23622248 |
| hsa-let-7b-5p | MIMAT0000063 | SC5D | 6309 | CLASH | 23622248 |
| hsa-let-7b-5p | MIMAT0000063 | ATXN2 | 6311 | CLASH//HITS-CLIP | 23622248\|23313552 |
| hsa-let-7b-5p | MIMAT0000063 | SCD | 6319 | CLASH//Proteomics | 18668040\|23622248 |
| hsa-let-7b-5p | MIMAT0000063 | SEMG2 | 6407 | Proteomics | 18668040 |
| hsa-let-7b-5p | MIMAT0000063 | SKI | 6497 | CLASH | 23622248 |
| hsa-let-7b-5p | MIMAT0000063 | SLC1A4 | 6509 | Proteomics//pSILAC | 18668040 |
| hsa-let-7b-5p | MIMAT0000063 | SLC20A1 | 6574 | PAR-CLIP | 23592263\|20371350 |
| hsa-let-7b-5p | MIMAT0000063 | SLC25A1 | 6576 | Proteomics | 18668040 |
| hsa-let-7b-5p | MIMAT0000063 | SMARCA1 | 6594 | Proteomics | 18668040 |
| hsa-let-7b-5p | MIMAT0000063 | SMARCA4 | 6597 | CLASH//Proteomics | 18668040\|23622248 |
| hsa-let-7b-5p | MIMAT0000063 | SMARCB1 | 6598 | CLASH | 23622248 |
| hsa-let-7b-5p | MIMAT0000063 | SMARCC1 | 6599 | Proteomics | 18668040 |
| hsa-let-7b-5p | MIMAT0000063 | SMARCC2 | 6601 | CLASH | 23622248 |
| hsa-let-7b-5p | MIMAT0000063 | SMARCD1 | 6602 | CLASH//Proteomics | 18668040\|23622248 |
| hsa-let-7b-5p | MIMAT0000063 | SUMO2 | 6613 | CLASH | 23622248 |
| hsa-let-7b-5p | MIMAT0000063 | SNRPA | 6626 | CLASH | 23622248 |
| hsa-let-7b-5p | MIMAT0000063 | SNRPE | 6635 | CLASH | 23622248 |
| hsa-let-7b-5p | MIMAT0000063 | SOD2 | 6648 | PAR-CLIP | 22012620 |
| hsa-let-7b-5p | MIMAT0000063 | SON | 6651 | CLASH | 23622248 |
| hsa-let-7b-5p | MIMAT0000063 | SOX9 | 6662 | Microarray | 17699775 |
| hsa-let-7b-5p | MIMAT0000063 | SP1 | 6667 | CLASH | 23622248 |
| hsa-let-7b-5p | MIMAT0000063 | SP100 | 6672 | Proteomics | 18668040 |
| hsa-let-7b-5p | MIMAT0000063 | SPN | 6693 | CLASH | 23622248 |
| hsa-let-7b-5p | MIMAT0000063 | SPR | 6697 | Proteomics | 18668040 |
| hsa-let-7b-5p | MIMAT0000063 | SPTBN2 | 6712 | CLASH | 23622248 |
| hsa-let-7b-5p | MIMAT0000063 | TROVE2 | 6738 | Proteomics | 18668040 |
| hsa-let-7b-5p | MIMAT0000063 | SSR1 | 6745 | Microarray | 17699775 |
| hsa-let-7b-5p | MIMAT0000063 | ST13 | 6767 | CLASH | 23622248 |

| | | | | | |
|---|---|---|---|---|---|
| hsa-let-7b-5p | MIMAT0000063 | STAT2 | 6773 | HITS-CLIP | 23706177 |
| hsa-let-7b-5p | MIMAT0000063 | STIM1 | 6786 | Proteomics | 18668040 |
| hsa-let-7b-5p | MIMAT0000063 | STK4 | 6789 | PAR-CLIP | 23446348 |
| hsa-let-7b-5p | MIMAT0000063 | AURKA | 6790 | Microarray//Proteomics | 17699775\|18668040 |
| hsa-let-7b-5p | MIMAT0000063 | STRN | 6801 | PAR-CLIP | 24398324\|20371350 |
| hsa-let-7b-5p | MIMAT0000063 | STX3 | 6809 | PAR-CLIP | 27292025 |
| hsa-let-7b-5p | MIMAT0000063 | SUOX | 6821 | PAR-CLIP | 21572407 |
| hsa-let-7b-5p | MIMAT0000063 | SURF4 | 6836 | PAR-CLIP | 20371350 |
| hsa-let-7b-5p | MIMAT0000063 | SYT1 | 6857 | PAR-CLIP | 26701625 |
| hsa-let-7b-5p | MIMAT0000063 | TAF9 | 6880 | Proteomics | 18668040 |
| hsa-let-7b-5p | MIMAT0000063 | TCOF1 | 6949 | Proteomics | 18668040 |
| hsa-let-7b-5p | MIMAT0000063 | TGFBR1 | 7046 | ELISA//GFP reporter assay//Western blot | 24978044 |
| hsa-let-7b-5p | MIMAT0000063 | TGFBR3 | 7049 | PAR-CLIP | 21572407 |
| hsa-let-7b-5p | MIMAT0000063 | THBS1 | 7057 | PAR-CLIP//Proteomics//pSILAC | 18668040\|24398324 |
| hsa-let-7b-5p | MIMAT0000063 | TIAM1 | 7074 | CLASH | 23622248 |
| hsa-let-7b-5p | MIMAT0000063 | TJP1 | 7082 | CLASH | 23622248 |
| hsa-let-7b-5p | MIMAT0000063 | TLN1 | 7094 | CLASH | 23622248 |
| hsa-let-7b-5p | MIMAT0000063 | TLR4 | 7099 | Luciferase reporter assay | 23437218 |
| hsa-let-7b-5p | MIMAT0000063 | NR2E1 | 7101 | immunoblot//Luciferase reporter assay//Northern blot//Western blot | 20133835 |
| hsa-let-7b-5p | MIMAT0000063 | TRAPPC10 | 7109 | PAR-CLIP | 26701625 |
| hsa-let-7b-5p | MIMAT0000063 | TPBG | 7162 | Proteomics | 18668040 |
| hsa-let-7b-5p | MIMAT0000063 | TPD52L2 | 7165 | CLASH | 23622248 |
| hsa-let-7b-5p | MIMAT0000063 | TPM4 | 7171 | CLASH | 23622248 |
| hsa-let-7b-5p | MIMAT0000063 | TPP2 | 7174 | Proteomics | 18668040 |
| hsa-let-7b-5p | MIMAT0000063 | TPT1 | 7178 | CLASH | 23622248 |
| hsa-let-7b-5p | MIMAT0000063 | TST | 7263 | Proteomics | 18668040 |
| hsa-let-7b-5p | MIMAT0000063 | TUBB2A | 7280 | PAR-CLIP | 23592263 |
| hsa-let-7b-5p | MIMAT0000063 | TYMS | 7298 | Proteomics//pSILAC | 18668040 |
| hsa-let-7b-5p | MIMAT0000063 | UBA1 | 7317 | CLASH | 23622248 |
| hsa-let-7b-5p | MIMAT0000063 | UBE2A | 7319 | CLASH | 23622248 |
| hsa-let-7b-5p | MIMAT0000063 | UBE2D2 | 7322 | Proteomics | 18668040 |
| hsa-let-7b-5p | MIMAT0000063 | UBE2D3 | 7323 | Proteomics | 18668040 |
| hsa-let-7b-5p | MIMAT0000063 | UBE2I | 7329 | Proteomics | 18668040 |
| hsa-let-7b-5p | MIMAT0000063 | SUMO1 | 7341 | PAR-CLIP | 24398324 |

| | | | | | |
|---|---|---|---|---|---|
| hsa-let-7b-5p | MIMAT0000063 | UGT8 | 7368 | Proteomics//pSILAC | 18668040 |
| hsa-let-7b-5p | MIMAT0000063 | UCK2 | 7371 | Proteomics | 18668040 |
| hsa-let-7b-5p | MIMAT0000063 | UTRN | 7402 | CLASH | 23622248 |
| hsa-let-7b-5p | MIMAT0000063 | VCL | 7414 | PAR-CLIP | 23592263 |
| hsa-let-7b-5p | MIMAT0000063 | YWHAE | 7531 | CLASH | 23622248 |
| hsa-let-7b-5p | MIMAT0000063 | YWHAZ | 7534 | CLASH//PAR-CLIP | 23622248\|23592263 |
| hsa-let-7b-5p | MIMAT0000063 | ZNF3 | 7551 | CLASH | 23622248 |
| hsa-let-7b-5p | MIMAT0000063 | ZNF8 | 7554 | PAR-CLIP | 23592263 |
| hsa-let-7b-5p | MIMAT0000063 | ZNF28 | 7576 | PAR-CLIP | 23592263 |
| hsa-let-7b-5p | MIMAT0000063 | ZNF148 | 7707 | CLASH | 23622248 |
| hsa-let-7b-5p | MIMAT0000063 | ZMYM2 | 7750 | CLASH | 23622248 |
| hsa-let-7b-5p | MIMAT0000063 | ZNF200 | 7752 | PAR-CLIP | 21572407\|20371350 |
| hsa-let-7b-5p | MIMAT0000063 | ZNF207 | 7756 | CLASH | 23622248 |
| hsa-let-7b-5p | MIMAT0000063 | SLC30A1 | 7779 | Proteomics | 18668040 |
| hsa-let-7b-5p | MIMAT0000063 | BTG2 | 7832 | CLASH | 23622248 |
| hsa-let-7b-5p | MIMAT0000063 | PXDN | 7837 | Proteomics//pSILAC | 18668040 |
| hsa-let-7b-5p | MIMAT0000063 | BRPF1 | 7862 | CLASH | 23622248 |
| hsa-let-7b-5p | MIMAT0000063 | PRRC2A | 7916 | CLASH//Proteomics//pSILAC | 18668040\|23622248 |
| hsa-let-7b-5p | MIMAT0000063 | BAG6 | 7917 | CLASH | 23622248 |
| hsa-let-7b-5p | MIMAT0000063 | ARHGEF5 | 7984 | CLASH | 23622248 |
| hsa-let-7b-5p | MIMAT0000063 | UBXN8 | 7993 | Proteomics | 18668040 |
| hsa-let-7b-5p | MIMAT0000063 | NUP214 | 8021 | CLASH//Proteomics | 18668040\|23622248 |
| hsa-let-7b-5p | MIMAT0000063 | MLLT10 | 8028 | PAR-CLIP | 23592263\|27292025 |
| hsa-let-7b-5p | MIMAT0000063 | PDHX | 8050 | Proteomics | 18668040 |
| hsa-let-7b-5p | MIMAT0000063 | PTP4A2 | 8073 | CLASH | 23622248 |
| hsa-let-7b-5p | MIMAT0000063 | KMT2D | 8085 | PAR-CLIP | 24398324\|21572407 |
| hsa-let-7b-5p | MIMAT0000063 | HMGA2 | 8091 | HITS-CLIP//Immunohistochemistry//Luciferase reporter assay//Microarray//PAR-CLIP//qRT-PCR//QRTPCR//Western blot | 17437991\|23482325\|23318420\|20371350\|21572407\|25600877\|26701625 |
| hsa-let-7b-5p | MIMAT0000063 | COIL | 8161 | PAR-CLIP//Proteomics | 18668040\|23446348\|21572407\|20371350 |
| hsa-let-7b-5p | MIMAT0000063 | NCOA3 | 8202 | PAR-CLIP | 24398324\|26701625 |
| hsa-let-7b-5p | MIMAT0000063 | SMC1A | 8243 | CLASH//PAR-CLIP | 23622248\|23446348\|21572407\|20371350\|26701625 |

| | | | | | |
|---|---|---|---|---|---|
| hsa-let-7b-5p | MIMAT0000063 | NAA10 | 8260 | CLASH | 23622248 |
| hsa-let-7b-5p | MIMAT0000063 | SLC10A3 | 8273 | CLASH | 23622248 |
| hsa-let-7b-5p | MIMAT0000063 | FZD9 | 8326 | PAR-CLIP | 23446348\|21572407 |
| hsa-let-7b-5p | MIMAT0000063 | HIST1H3B | 8358 | CLASH | 23622248 |
| hsa-let-7b-5p | MIMAT0000063 | DYRK3 | 8444 | PAR-CLIP | 26701625 |
| hsa-let-7b-5p | MIMAT0000063 | DHX16 | 8449 | Proteomics | 18668040 |
| hsa-let-7b-5p | MIMAT0000063 | CUL3 | 8452 | Proteomics | 18668040 |
| hsa-let-7b-5p | MIMAT0000063 | CUL2 | 8453 | CLASH | 23622248 |
| hsa-let-7b-5p | MIMAT0000063 | CUL1 | 8454 | Proteomics | 18668040 |
| hsa-let-7b-5p | MIMAT0000063 | IRS4 | 8471 | CLASH | 23622248 |
| hsa-let-7b-5p | MIMAT0000063 | DENR | 8562 | CLASH | 23622248 |
| hsa-let-7b-5p | MIMAT0000063 | THOC5 | 8563 | CLASH | 23622248 |
| hsa-let-7b-5p | MIMAT0000063 | KHSRP | 8570 | CLASH | 23622248 |
| hsa-let-7b-5p | MIMAT0000063 | NOP14 | 8602 | Proteomics | 18668040 |
| hsa-let-7b-5p | MIMAT0000063 | SLC25A12 | 8604 | Proteomics | 18668040 |
| hsa-let-7b-5p | MIMAT0000063 | USO1 | 8615 | Proteomics | 18668040 |
| hsa-let-7b-5p | MIMAT0000063 | RTCA | 8634 | CLASH//Proteomics | 18668040\|23622248 |
| hsa-let-7b-5p | MIMAT0000063 | SOCS1 | 8651 | PAR-CLIP | 23592263 |
| hsa-let-7b-5p | MIMAT0000063 | IRS2 | 8660 | CLASH//Flow//Luciferase reporter assay//qRT-PCR//Western blot | 23622248\|24810113 |
| hsa-let-7b-5p | MIMAT0000063 | EIF3C | 8663 | CLASH | 23622248 |
| hsa-let-7b-5p | MIMAT0000063 | EIF3D | 8664 | CLASH | 23622248 |
| hsa-let-7b-5p | MIMAT0000063 | GBF1 | 8729 | CLASH | 23622248 |
| hsa-let-7b-5p | MIMAT0000063 | RNMT | 8731 | Proteomics | 18668040 |
| hsa-let-7b-5p | MIMAT0000063 | TNFSF12 | 8742 | CLASH | 23622248 |
| hsa-let-7b-5p | MIMAT0000063 | TNFSF9 | 8744 | PAR-CLIP | 23592263\|26701625 |
| hsa-let-7b-5p | MIMAT0000063 | SNAP23 | 8773 | Proteomics//pSILAC | 18668040 |
| hsa-let-7b-5p | MIMAT0000063 | RIOK3 | 8780 | CLASH | 23622248 |
| hsa-let-7b-5p | MIMAT0000063 | TNFRSF10B | 8795 | qRT-PCR//Western blot | 24120475 |
| hsa-let-7b-5p | MIMAT0000063 | PEX11B | 8799 | PAR-CLIP//Proteomics | 18668040\|21572407 |
| hsa-let-7b-5p | MIMAT0000063 | TRIM24 | 8805 | CLASH//Proteomics | 18668040\|23622248 |
| hsa-let-7b-5p | MIMAT0000063 | CCNK | 8812 | Proteomics | 18668040 |
| hsa-let-7b-5p | MIMAT0000063 | SLC5A6 | 8884 | PAR-CLIP | 24398324\|21572407 |
| hsa-let-7b-5p | MIMAT0000063 | DDX18 | 8886 | CLASH | 23622248 |
| hsa-let-7b-5p | MIMAT0000063 | EIF2B3 | 8891 | Proteomics | 18668040 |
| hsa-let-7b-5p | MIMAT0000063 | CCNA1 | 8900 | Reporter assay | 18379589 |

| | | | | | |
|---|---|---|---|---|---|
| hsa-let-7b-5p | MIMAT0000063 | MBD2 | 8932 | PAR-CLIP | 24398324\|23446348\|21572407\|20371350 |
| hsa-let-7b-5p | MIMAT0000063 | WASF1 | 8936 | CLASH | 23622248 |
| hsa-let-7b-5p | MIMAT0000063 | WASL | 8976 | CLASH//PAR-CLIP | 23622248\|22012620\|21572407\|20371350 |
| hsa-let-7b-5p | MIMAT0000063 | BAZ1B | 9031 | CLASH//Proteomics | 18668040\|23622248 |
| hsa-let-7b-5p | MIMAT0000063 | UBA3 | 9039 | Proteomics | 18668040 |
| hsa-let-7b-5p | MIMAT0000063 | UBE2M | 9040 | Proteomics | 18668040 |
| hsa-let-7b-5p | MIMAT0000063 | MAP7 | 9053 | Proteomics | 18668040 |
| hsa-let-7b-5p | MIMAT0000063 | PAPSS1 | 9061 | Proteomics | 18668040 |
| hsa-let-7b-5p | MIMAT0000063 | CLDN12 | 9069 | PAR-CLIP | 24398324\|23446348\|20371350\|27292025 |
| hsa-let-7b-5p | MIMAT0000063 | USP14 | 9097 | CLASH | 23622248 |
| hsa-let-7b-5p | MIMAT0000063 | USP10 | 9100 | Proteomics | 18668040 |
| hsa-let-7b-5p | MIMAT0000063 | CCNB2 | 9133 | CLASH | 23622248 |
| hsa-let-7b-5p | MIMAT0000063 | ATG12 | 9140 | PAR-CLIP | 23592263 |
| hsa-let-7b-5p | MIMAT0000063 | SYNGR2 | 9144 | CLASH//Proteomics | 18668040\|23622248 |
| hsa-let-7b-5p | MIMAT0000063 | HGS | 9146 | CLASH | 23622248 |
| hsa-let-7b-5p | MIMAT0000063 | DDX21 | 9188 | Proteomics | 18668040 |
| hsa-let-7b-5p | MIMAT0000063 | DEDD | 9191 | CLASH | 23622248 |
| hsa-let-7b-5p | MIMAT0000063 | AURKB | 9212 | Proteomics//pSILAC | 18668040 |
| hsa-let-7b-5p | MIMAT0000063 | TIAF1 | 9220 | PAR-CLIP | 26701625 |
| hsa-let-7b-5p | MIMAT0000063 | NOLC1 | 9221 | Proteomics | 18668040 |
| hsa-let-7b-5p | MIMAT0000063 | PTTG1 | 9232 | CLASH | 23622248 |
| hsa-let-7b-5p | MIMAT0000063 | TBRG4 | 9238 | Proteomics | 18668040 |
| hsa-let-7b-5p | MIMAT0000063 | MED14 | 9282 | CLASH | 23622248 |
| hsa-let-7b-5p | MIMAT0000063 | ADGRG1 | 9289 | Proteomics//pSILAC | 18668040 |
| hsa-let-7b-5p | MIMAT0000063 | ATP6V1F | 9296 | PAR-CLIP//Proteomics//pSILAC | 18668040\|23592263\|24398324\|26701625\|27292025 |
| hsa-let-7b-5p | MIMAT0000063 | COPS2 | 9318 | Proteomics | 18668040 |
| hsa-let-7b-5p | MIMAT0000063 | TRIP12 | 9320 | Proteomics | 18668040 |
| hsa-let-7b-5p | MIMAT0000063 | GTF3C4 | 9329 | Proteomics | 18668040 |
| hsa-let-7b-5p | MIMAT0000063 | VAMP3 | 9341 | Proteomics | 18668040 |
| hsa-let-7b-5p | MIMAT0000063 | SLC9A3R1 | 9368 | Proteomics | 18668040 |
| hsa-let-7b-5p | MIMAT0000063 | CIAO1 | 9391 | Proteomics//pSILAC | 18668040 |
| hsa-let-7b-5p | MIMAT0000063 | FADS2 | 9415 | CLASH//Proteomics//pSILAC | 18668040\|23622248 |
| hsa-let-7b-5p | MIMAT0000063 | HAND1 | 9421 | CLASH//PAR-CLIP | 23622248\|21572407 |

| | | | | | |
|---|---|---|---|---|---|
| hsa-let-7b-5p | MIMAT0000063 | ZNF264 | 9422 | PAR-CLIP | 21572407 |
| hsa-let-7b-5p | MIMAT0000063 | QKI | 9444 | CLASH | 23622248 |
| hsa-let-7b-5p | MIMAT0000063 | GGPS1 | 9453 | CLASH | 23622248 |
| hsa-let-7b-5p | MIMAT0000063 | PCYT1B | 9468 | Proteomics | 18668040 |
| hsa-let-7b-5p | MIMAT0000063 | ONECUT2 | 9480 | PAR-CLIP | 23446348 |
| hsa-let-7b-5p | MIMAT0000063 | FXR2 | 9513 | CLASH | 23622248 |
| hsa-let-7b-5p | MIMAT0000063 | EEF1E1 | 9521 | Proteomics | 18668040 |
| hsa-let-7b-5p | MIMAT0000063 | SCAMP1 | 9522 | Proteomics | 18668040 |
| hsa-let-7b-5p | MIMAT0000063 | BAG5 | 9529 | Proteomics | 18668040 |
| hsa-let-7b-5p | MIMAT0000063 | ATP6V1G1 | 9550 | PAR-CLIP | 22100165\|22291592\|23446348 |
| hsa-let-7b-5p | MIMAT0000063 | SOX13 | 9580 | CLASH | 23622248 |
| hsa-let-7b-5p | MIMAT0000063 | ENTPD4 | 9583 | CLASH | 23622248 |
| hsa-let-7b-5p | MIMAT0000063 | NCOR1 | 9611 | CLASH | 23622248 |
| hsa-let-7b-5p | MIMAT0000063 | NUP155 | 9631 | CLASH//PAR-CLIP//Proteomics | 18668040\|23622248\|23446348\|21572407\|20371350 |
| hsa-let-7b-5p | MIMAT0000063 | CLCA2 | 9635 | CLASH | 23622248 |
| hsa-let-7b-5p | MIMAT0000063 | SH3PXD2A | 9644 | CLASH | 23622248 |
| hsa-let-7b-5p | MIMAT0000063 | CTR9 | 9646 | CLASH | 23622248 |
| hsa-let-7b-5p | MIMAT0000063 | HS2ST1 | 9653 | Proteomics | 18668040 |
| hsa-let-7b-5p | MIMAT0000063 | CEP135 | 9662 | PAR-CLIP | 24398324 |
| hsa-let-7b-5p | MIMAT0000063 | KDM4A | 9682 | CLASH | 23622248 |
| hsa-let-7b-5p | MIMAT0000063 | CLINT1 | 9685 | CLASH | 23622248 |
| hsa-let-7b-5p | MIMAT0000063 | BZW1 | 9689 | PAR-CLIP | 23592263\|20371350\|26701625 |
| hsa-let-7b-5p | MIMAT0000063 | KIAA0391 | 9692 | PAR-CLIP | 21572407 |
| hsa-let-7b-5p | MIMAT0000063 | PUM1 | 9698 | CLASH//Proteomics | 18668040\|23622248 |
| hsa-let-7b-5p | MIMAT0000063 | ESPL1 | 9700 | PAR-CLIP | 23446348\|21572407 |
| hsa-let-7b-5p | MIMAT0000063 | HERPUD1 | 9709 | HITS-CLIP//PAR-CLIP | 23592263\|23706177 |
| hsa-let-7b-5p | MIMAT0000063 | EIF4A3 | 9775 | HITS-CLIP | 23706177 |
| hsa-let-7b-5p | MIMAT0000063 | SNX17 | 9784 | PAR-CLIP | 22291592 |
| hsa-let-7b-5p | MIMAT0000063 | SCRN1 | 9805 | Proteomics | 18668040 |
| hsa-let-7b-5p | MIMAT0000063 | IP6K1 | 9807 | CLASH | 23622248 |
| hsa-let-7b-5p | MIMAT0000063 | RNF40 | 9810 | Proteomics | 18668040 |
| hsa-let-7b-5p | MIMAT0000063 | KIAA0141 | 9812 | CLASH | 23622248 |
| hsa-let-7b-5p | MIMAT0000063 | EFCAB14 | 9813 | CLASH | 23622248 |
| hsa-let-7b-5p | MIMAT0000063 | TSC22D2 | 9819 | PAR-CLIP | 23592263\|24398324 |
| hsa-let-7b-5p | MIMAT0000063 | AREL1 | 9870 | PAR-CLIP | 23592263 |
| hsa-let-7b-5p | MIMAT0000063 | ZC3H11A | 9877 | Proteomics | 18668040 |

| | | | | | |
|---|---|---|---|---|---|
| hsa-let-7b-5p | MIMAT0000063 | POM121 | 9883 | CLASH//Proteomics//pSILAC | 18668040\|23622248 |
| hsa-let-7b-5p | MIMAT0000063 | SMG7 | 9887 | CLASH | 23622248 |
| hsa-let-7b-5p | MIMAT0000063 | UBAP2L | 9898 | CLASH//Proteomics | 18668040\|23622248 |
| hsa-let-7b-5p | MIMAT0000063 | RBM19 | 9904 | Proteomics//pSILAC | 18668040 |
| hsa-let-7b-5p | MIMAT0000063 | DENND4B | 9909 | CLASH | 23622248 |
| hsa-let-7b-5p | MIMAT0000063 | NCAPD2 | 9918 | CLASH | 23622248 |
| hsa-let-7b-5p | MIMAT0000063 | SEC16A | 9919 | Proteomics | 18668040 |
| hsa-let-7b-5p | MIMAT0000063 | ZBTB5 | 9925 | PAR-CLIP | 23592263\|24398324\|21572407\|20371350 |
| hsa-let-7b-5p | MIMAT0000063 | LPGAT1 | 9926 | Proteomics | 18668040 |
| hsa-let-7b-5p | MIMAT0000063 | USP15 | 9958 | Proteomics | 18668040 |
| hsa-let-7b-5p | MIMAT0000063 | MED13 | 9969 | CLASH | 23622248 |
| hsa-let-7b-5p | MIMAT0000063 | NUP153 | 9972 | CLASH | 23622248 |
| hsa-let-7b-5p | MIMAT0000063 | HNRNPDL | 9987 | qRT-PCR | 17942906 |
| hsa-let-7b-5p | MIMAT0000063 | CHAF1A | 10036 | Proteomics | 18668040 |
| hsa-let-7b-5p | MIMAT0000063 | SCAMP3 | 10067 | qRT-PCR | 17942906 |
| hsa-let-7b-5p | MIMAT0000063 | HUWE1 | 10075 | CLASH | 23622248 |
| hsa-let-7b-5p | MIMAT0000063 | TSPAN3 | 10099 | CLASH | 23622248 |
| hsa-let-7b-5p | MIMAT0000063 | NUBP2 | 10101 | Proteomics | 18668040 |
| hsa-let-7b-5p | MIMAT0000063 | RBM12 | 10137 | CLASH | 23622248 |
| hsa-let-7b-5p | MIMAT0000063 | TRIM28 | 10155 | CLASH | 23622248 |
| hsa-let-7b-5p | MIMAT0000063 | FARP1 | 10160 | Microarray | 17699775 |
| hsa-let-7b-5p | MIMAT0000063 | LPCAT3 | 10162 | Proteomics | 18668040 |
| hsa-let-7b-5p | MIMAT0000063 | SLC25A13 | 10165 | CLASH//Proteomics//pSILAC | 18668040\|23622248 |
| hsa-let-7b-5p | MIMAT0000063 | ALG3 | 10195 | Proteomics//pSILAC | 18668040 |
| hsa-let-7b-5p | MIMAT0000063 | PSME3 | 10197 | CLASH//Proteomics | 18668040\|23622248 |
| hsa-let-7b-5p | MIMAT0000063 | NME6 | 10201 | Proteomics | 18668040 |
| hsa-let-7b-5p | MIMAT0000063 | GDF11 | 10220 | CLASH | 23622248 |
| hsa-let-7b-5p | MIMAT0000063 | ZNF443 | 10224 | HITS-CLIP | 23313552 |
| hsa-let-7b-5p | MIMAT0000063 | DCAF7 | 10238 | CLASH | 23622248 |
| hsa-let-7b-5p | MIMAT0000063 | CALCOCO2 | 10241 | Proteomics//pSILAC | 18668040 |
| hsa-let-7b-5p | MIMAT0000063 | GPHN | 10243 | CLASH | 23622248 |
| hsa-let-7b-5p | MIMAT0000063 | AKAP8 | 10270 | PAR-CLIP//Proteomics//pSILAC | 18668040\|21572407\|20371350 |

| | | | | | |
|---|---|---|---|---|---|
| hsa-let-7b-5p | MIMAT0000063 | SIGMAR1 | 10280 | Microarray//Proteomics//pSILAC | 17699775\|18668040 |
| hsa-let-7b-5p | MIMAT0000063 | MARCH6 | 10299 | CLASH | 23622248 |
| hsa-let-7b-5p | MIMAT0000063 | AKR1A1 | 10327 | Proteomics | 18668040 |
| hsa-let-7b-5p | MIMAT0000063 | PCGF3 | 10336 | CLASH//PAR-CLIP | 23622248\|23592263\|23446348\|20371350 |
| hsa-let-7b-5p | MIMAT0000063 | WARS2 | 10352 | Proteomics | 18668040 |
| hsa-let-7b-5p | MIMAT0000063 | TUBA1B | 10376 | CLASH | 23622248 |
| hsa-let-7b-5p | MIMAT0000063 | TUBB4A | 10382 | PAR-CLIP | 26701625 |
| hsa-let-7b-5p | MIMAT0000063 | SCML2 | 10389 | CLASH | 23622248 |
| hsa-let-7b-5p | MIMAT0000063 | DLC1 | 10395 | Microarray | 17699775 |
| hsa-let-7b-5p | MIMAT0000063 | NDRG1 | 10397 | CLASH | 23622248 |
| hsa-let-7b-5p | MIMAT0000063 | NSA2 | 10412 | Proteomics | 18668040 |
| hsa-let-7b-5p | MIMAT0000063 | YAP1 | 10413 | Microarray | 17699775 |
| hsa-let-7b-5p | MIMAT0000063 | CD2BP2 | 10421 | CLASH | 23622248 |
| hsa-let-7b-5p | MIMAT0000063 | CDIPT | 10423 | Proteomics//pSILAC | 18668040 |
| hsa-let-7b-5p | MIMAT0000063 | TUBGCP3 | 10426 | CLASH | 23622248 |
| hsa-let-7b-5p | MIMAT0000063 | ZER1 | 10444 | CLASH | 23622248 |
| hsa-let-7b-5p | MIMAT0000063 | SEC23B | 10483 | Proteomics | 18668040 |
| hsa-let-7b-5p | MIMAT0000063 | LRRC41 | 10489 | CLASH | 23622248 |
| hsa-let-7b-5p | MIMAT0000063 | UNC13B | 10497 | CLASH | 23622248 |
| hsa-let-7b-5p | MIMAT0000063 | DDX17 | 10521 | CLASH | 23622248 |
| hsa-let-7b-5p | MIMAT0000063 | IPO8 | 10526 | Proteomics | 18668040 |
| hsa-let-7b-5p | MIMAT0000063 | IPO7 | 10527 | CLASH | 23622248 |
| hsa-let-7b-5p | MIMAT0000063 | RPP38 | 10557 | Proteomics//pSILAC | 18668040 |
| hsa-let-7b-5p | MIMAT0000063 | PRPF8 | 10594 | CLASH | 23622248 |
| hsa-let-7b-5p | MIMAT0000063 | PDLIM5 | 10611 | PAR-CLIP | 26701625 |
| hsa-let-7b-5p | MIMAT0000063 | TGOLN2 | 10618 | PAR-CLIP | 20371350 |
| hsa-let-7b-5p | MIMAT0000063 | ARID3B | 10620 | CLASH//PAR-CLIP | 23622248\|23446348\|21572407\|20371350 |
| hsa-let-7b-5p | MIMAT0000063 | POLR3G | 10622 | CLASH | 23622248 |
| hsa-let-7b-5p | MIMAT0000063 | LEFTY1 | 10637 | PAR-CLIP | 22012620 |
| hsa-let-7b-5p | MIMAT0000063 | IGF2BP1 | 10642 | Immunoblot//Immunofluorescence//Luciferase reporter assay//PAR-CLIP//Proteomics//pSILAC//qRT-PCR//Western blot | 18668040\|21252116\|21572407\|23824794 |
| hsa-let-7b-5p | MIMAT0000063 | IGF2BP3 | 10643 | PAR-CLIP//Proteomics | 18668040\|21572407\|20371350 |

| | | | | | |
|---|---|---|---|---|---|
| hsa-let-7b-5p | MIMAT0000063 | IGF2BP2 | 10644 | Immunohistochemistry//Luciferase reporter assay//Proteomics//pSILAC//QRTPCR//Western blot | 18668040\|23482325 |
| hsa-let-7b-5p | MIMAT0000063 | CELF1 | 10658 | PAR-CLIP | 24398324 |
| hsa-let-7b-5p | MIMAT0000063 | CTCF | 10664 | CLASH | 23622248 |
| hsa-let-7b-5p | MIMAT0000063 | SLC12A7 | 10723 | PAR-CLIP | 22291592 |
| hsa-let-7b-5p | MIMAT0000063 | RAI1 | 10743 | CLASH | 23622248 |
| hsa-let-7b-5p | MIMAT0000063 | KIF1C | 10749 | CLASH | 23622248 |
| hsa-let-7b-5p | MIMAT0000063 | AHCYL1 | 10768 | CLASH | 23622248 |
| hsa-let-7b-5p | MIMAT0000063 | ARPP19 | 10776 | CLASH | 23622248 |
| hsa-let-7b-5p | MIMAT0000063 | WDR4 | 10785 | Proteomics | 18668040 |
| hsa-let-7b-5p | MIMAT0000063 | ZNF460 | 10794 | PAR-CLIP | 21572407\|20371350 |
| hsa-let-7b-5p | MIMAT0000063 | TUBGCP2 | 10844 | Proteomics | 18668040 |
| hsa-let-7b-5p | MIMAT0000063 | CLPX | 10845 | Proteomics | 18668040 |
| hsa-let-7b-5p | MIMAT0000063 | SRCAP | 10847 | CLASH | 23622248 |
| hsa-let-7b-5p | MIMAT0000063 | PGRMC1 | 10857 | PAR-CLIP//Proteomics//pSILAC | 18668040\|21572407 |
| hsa-let-7b-5p | MIMAT0000063 | WDR3 | 10885 | CLASH | 23622248 |
| hsa-let-7b-5p | MIMAT0000063 | RAB10 | 10890 | CLASH | 23622248 |
| hsa-let-7b-5p | MIMAT0000063 | PPARGC1A | 10891 | CLASH | 23622248 |
| hsa-let-7b-5p | MIMAT0000063 | TXNL4A | 10907 | CLASH | 23622248 |
| hsa-let-7b-5p | MIMAT0000063 | PAPOLA | 10914 | CLASH | 23622248 |
| hsa-let-7b-5p | MIMAT0000063 | POP1 | 10940 | Proteomics | 18668040 |
| hsa-let-7b-5p | MIMAT0000063 | KDELR1 | 10945 | CLASH | 23622248 |
| hsa-let-7b-5p | MIMAT0000063 | LMAN2 | 10960 | CLASH | 23622248 |
| hsa-let-7b-5p | MIMAT0000063 | STIP1 | 10963 | CLASH | 23622248 |
| hsa-let-7b-5p | MIMAT0000063 | ASCC3 | 10973 | Proteomics | 18668040 |
| hsa-let-7b-5p | MIMAT0000063 | GCN1 | 10985 | CLASH | 23622248 |
| hsa-let-7b-5p | MIMAT0000063 | SLC27A2 | 11001 | Proteomics | 18668040 |
| hsa-let-7b-5p | MIMAT0000063 | ADRM1 | 11047 | CLASH | 23622248 |
| hsa-let-7b-5p | MIMAT0000063 | TMEM115 | 11070 | CLASH | 23622248 |
| hsa-let-7b-5p | MIMAT0000063 | HNRNPUL1 | 11100 | CLASH | 23622248 |
| hsa-let-7b-5p | MIMAT0000063 | ATE1 | 11101 | CLASH | 23622248 |
| hsa-let-7b-5p | MIMAT0000063 | PRDM4 | 11108 | CLASH | 23622248 |
| hsa-let-7b-5p | MIMAT0000063 | LSM6 | 11157 | Proteomics | 18668040 |

| | | | | | |
|---|---|---|---|---|---|
| hsa-let-7b-5p | MIMAT0000063 | RABL2B | 11158 | PAR-CLIP | 23592263\|26701625 |
| hsa-let-7b-5p | MIMAT0000063 | RABL2A | 11159 | PAR-CLIP | 23592263\|26701625 |
| hsa-let-7b-5p | MIMAT0000063 | FSTL1 | 11167 | CLASH | 23622248 |
| hsa-let-7b-5p | MIMAT0000063 | BAZ2A | 11176 | CLASH | 23622248 |
| hsa-let-7b-5p | MIMAT0000063 | BAZ1A | 11177 | Proteomics | 18668040 |
| hsa-let-7b-5p | MIMAT0000063 | ABCB8 | 11194 | CLASH | 23622248 |
| hsa-let-7b-5p | MIMAT0000063 | SUPT16H | 11198 | CLASH | 23622248 |
| hsa-let-7b-5p | MIMAT0000063 | DDX20 | 11218 | CLASH | 23622248 |
| hsa-let-7b-5p | MIMAT0000063 | PRAF2 | 11230 | CLASH | 23622248 |
| hsa-let-7b-5p | MIMAT0000063 | PDCD10 | 11235 | Proteomics | 18668040 |
| hsa-let-7b-5p | MIMAT0000063 | PMF1 | 11243 | Proteomics | 18668040 |
| hsa-let-7b-5p | MIMAT0000063 | DUSP12 | 11266 | Proteomics//pSILAC | 18668040 |
| hsa-let-7b-5p | MIMAT0000063 | ATXN2L | 11273 | CLASH | 23622248 |
| hsa-let-7b-5p | MIMAT0000063 | STK38 | 11329 | CLASH | 23622248 |
| hsa-let-7b-5p | MIMAT0000063 | IKZF3 | 22806 | HITS-CLIP//PAR-CLIP | 23446348\|23706177 |
| hsa-let-7b-5p | MIMAT0000063 | COPG1 | 22820 | Proteomics | 18668040 |
| hsa-let-7b-5p | MIMAT0000063 | DNAJC8 | 22826 | CLASH | 23622248 |
| hsa-let-7b-5p | MIMAT0000063 | SCAF8 | 22828 | CLASH | 23622248 |
| hsa-let-7b-5p | MIMAT0000063 | ZNF652 | 22834 | CLASH | 23622248 |
| hsa-let-7b-5p | MIMAT0000063 | RNF44 | 22838 | PAR-CLIP | 24398324\|20371350\|27292025 |
| hsa-let-7b-5p | MIMAT0000063 | ZNF507 | 22847 | CLASH | 23622248 |
| hsa-let-7b-5p | MIMAT0000063 | CPEB3 | 22849 | Luciferase reporter assay//Microarray//qRT-PCR | 22995917 |
| hsa-let-7b-5p | MIMAT0000063 | FNDC3A | 22862 | PAR-CLIP//Proteomics//pSILAC | 18668040\|23592263 |
| hsa-let-7b-5p | MIMAT0000063 | DZIP1 | 22873 | Microarray | 17699775 |
| hsa-let-7b-5p | MIMAT0000063 | BAHD1 | 22893 | CLASH | 23622248 |
| hsa-let-7b-5p | MIMAT0000063 | RPIA | 22934 | Luciferase reporter assay//Reporter assay | 15131085 |
| hsa-let-7b-5p | MIMAT0000063 | PDCD11 | 22984 | CLASH//Proteomics | 18668040\|23622248 |
| hsa-let-7b-5p | MIMAT0000063 | CNOT1 | 23019 | CLASH | 23622248 |
| hsa-let-7b-5p | MIMAT0000063 | RBM34 | 23029 | Proteomics | 18668040 |
| hsa-let-7b-5p | MIMAT0000063 | XPO7 | 23039 | Proteomics | 18668040 |
| hsa-let-7b-5p | MIMAT0000063 | SMG1 | 23049 | CLASH | 23622248 |
| hsa-let-7b-5p | MIMAT0000063 | ZNF609 | 23060 | PAR-CLIP | 26701625 |
| hsa-let-7b-5p | MIMAT0000063 | RRP1B | 23076 | Proteomics//pSILAC | 18668040 |
| hsa-let-7b-5p | MIMAT0000063 | VWA8 | 23078 | Proteomics | 18668040 |

| | | | | | |
|---|---|---|---|---|---|
| hsa-let-7b-5p | MIMAT0000063 | PPRC1 | 23082 | CLASH | 23622248 |
| hsa-let-7b-5p | MIMAT0000063 | ERC1 | 23085 | Proteomics//pSILAC | 18668040 |
| hsa-let-7b-5p | MIMAT0000063 | PEG10 | 23089 | PAR-CLIP | 20371350 |
| hsa-let-7b-5p | MIMAT0000063 | ARHGAP 26 | 23092 | CLASH | 23622248 |
| hsa-let-7b-5p | MIMAT0000063 | MCF2L2 | 23101 | HITS-CLIP//PAR-CLIP | 21572407\|20371350\|23706177 |
| hsa-let-7b-5p | MIMAT0000063 | TNRC6B | 23112 | CLASH | 23622248 |
| hsa-let-7b-5p | MIMAT0000063 | TAB2 | 23118 | CLASH//Microarray | 17699775\|23622248 |
| hsa-let-7b-5p | MIMAT0000063 | PLXND1 | 23129 | PAR-CLIP | 23592263 |
| hsa-let-7b-5p | MIMAT0000063 | EPB41L3 | 23136 | CLASH | 23622248 |
| hsa-let-7b-5p | MIMAT0000063 | GGA3 | 23163 | PAR-CLIP | 24398324 |
| hsa-let-7b-5p | MIMAT0000063 | TTLL12 | 23170 | Proteomics | 18668040 |
| hsa-let-7b-5p | MIMAT0000063 | ATG4B | 23192 | Proteomics | 18668040 |
| hsa-let-7b-5p | MIMAT0000063 | PMPCA | 23203 | PAR-CLIP | 21572407\|20371350 |
| hsa-let-7b-5p | MIMAT0000063 | ARL6IP1 | 23204 | CLASH | 23622248 |
| hsa-let-7b-5p | MIMAT0000063 | SYNE2 | 23224 | CLASH | 23622248 |
| hsa-let-7b-5p | MIMAT0000063 | PDS5A | 23244 | CLASH | 23622248 |
| hsa-let-7b-5p | MIMAT0000063 | ADGRL2 | 23266 | CLASH | 23622248 |
| hsa-let-7b-5p | MIMAT0000063 | DNMBP | 23268 | CLASH | 23622248 |
| hsa-let-7b-5p | MIMAT0000063 | CLUH | 23277 | CLASH//Proteomics | 18668040\|23622248 |
| hsa-let-7b-5p | MIMAT0000063 | ICOSLG | 23308 | PAR-CLIP | 23592263 |
| hsa-let-7b-5p | MIMAT0000063 | KIAA093 0 | 23313 | PAR-CLIP | 23592263 |
| hsa-let-7b-5p | MIMAT0000063 | ZCCHC11 | 23318 | CLASH | 23622248 |
| hsa-let-7b-5p | MIMAT0000063 | USP22 | 23326 | CLASH | 23622248 |
| hsa-let-7b-5p | MIMAT0000063 | VPS39 | 23339 | Proteomics//pSILAC | 18668040 |
| hsa-let-7b-5p | MIMAT0000063 | SYNE1 | 23345 | Proteomics | 18668040 |
| hsa-let-7b-5p | MIMAT0000063 | ZNF629 | 23361 | CLASH | 23622248 |
| hsa-let-7b-5p | MIMAT0000063 | PSD3 | 23362 | CLASH | 23622248 |
| hsa-let-7b-5p | MIMAT0000063 | LARP1 | 23367 | CLASH | 23622248 |
| hsa-let-7b-5p | MIMAT0000063 | RRP8 | 23378 | Proteomics//pSILAC | 18668040 |
| hsa-let-7b-5p | MIMAT0000063 | AHCYL2 | 23382 | PAR-CLIP | 23592263 |
| hsa-let-7b-5p | MIMAT0000063 | ADNP | 23394 | CLASH | 23622248 |
| hsa-let-7b-5p | MIMAT0000063 | DICER1 | 23405 | Immunoblot | 18812516 |
| hsa-let-7b-5p | MIMAT0000063 | SF3B1 | 23451 | CLASH | 23622248 |
| hsa-let-7b-5p | MIMAT0000063 | ABCB10 | 23456 | Proteomics | 18668040 |
| hsa-let-7b-5p | MIMAT0000063 | CBX6 | 23466 | CLASH | 23622248 |
| hsa-let-7b-5p | MIMAT0000063 | CBX5 | 23468 | CLASH//PAR-CLIP | 23622248\|21572407 |
| hsa-let-7b-5p | MIMAT0000063 | SEC11A | 23478 | Proteomics | 18668040 |

| | | | | | |
|---|---|---|---|---|---|
| hsa-let-7b-5p | MIMAT0000063 | RBFOX2 | 23543 | CLASH//PAR-CLIP//Proteomics | 18668040\|23622248\|24398324\|23446348\|22012620\|21572407\|20371350 |
| hsa-let-7b-5p | MIMAT0000063 | CARHSP1 | 23589 | Proteomics//pSILAC | 18668040 |
| hsa-let-7b-5p | MIMAT0000063 | ACOT9 | 23597 | PAR-CLIP | 23592263\|23446348 |
| hsa-let-7b-5p | MIMAT0000063 | MKRN2 | 23609 | Proteomics | 18668040 |
| hsa-let-7b-5p | MIMAT0000063 | KPNA6 | 23633 | CLASH | 23622248 |
| hsa-let-7b-5p | MIMAT0000063 | PLD3 | 23646 | PAR-CLIP | 26701625 |
| hsa-let-7b-5p | MIMAT0000063 | ARFIP2 | 23647 | Proteomics | 18668040 |
| hsa-let-7b-5p | MIMAT0000063 | PLXNB2 | 23654 | CLASH | 23622248 |
| hsa-let-7b-5p | MIMAT0000063 | TMEM2 | 23670 | Proteomics//pSILAC | 18668040 |
| hsa-let-7b-5p | MIMAT0000063 | RAB38 | 23682 | Proteomics | 18668040 |
| hsa-let-7b-5p | MIMAT0000063 | IFIT5 | 24138 | Proteomics//pSILAC | 18668040 |
| hsa-let-7b-5p | MIMAT0000063 | RAB3GAP2 | 25782 | Proteomics | 18668040 |
| hsa-let-7b-5p | MIMAT0000063 | CIZ1 | 25792 | CLASH | 23622248 |
| hsa-let-7b-5p | MIMAT0000063 | ARIH1 | 25820 | PAR-CLIP | 21572407 |
| hsa-let-7b-5p | MIMAT0000063 | NIPBL | 25836 | Proteomics | 18668040 |
| hsa-let-7b-5p | MIMAT0000063 | YIPF3 | 25844 | CLASH | 23622248 |
| hsa-let-7b-5p | MIMAT0000063 | POLR1A | 25885 | Proteomics | 18668040 |
| hsa-let-7b-5p | MIMAT0000063 | INTS7 | 25896 | HITS-CLIP//Proteomics | 18668040\|23706177 |
| hsa-let-7b-5p | MIMAT0000063 | AHCTF1 | 25909 | Proteomics | 18668040 |
| hsa-let-7b-5p | MIMAT0000063 | GEMIN5 | 25929 | Proteomics | 18668040 |
| hsa-let-7b-5p | MIMAT0000063 | PTPN23 | 25930 | CLASH | 23622248 |
| hsa-let-7b-5p | MIMAT0000063 | VIRMA | 25962 | CLASH | 23622248 |
| hsa-let-7b-5p | MIMAT0000063 | MMACHC | 25974 | CLASH | 23622248 |
| hsa-let-7b-5p | MIMAT0000063 | RPAP1 | 26015 | CLASH | 23622248 |
| hsa-let-7b-5p | MIMAT0000063 | LRIG1 | 26018 | Luciferase reporter assay//Microarray//qRT-PCR | 22995917 |
| hsa-let-7b-5p | MIMAT0000063 | PTCD1 | 26024 | Proteomics | 18668040 |
| hsa-let-7b-5p | MIMAT0000063 | IPCEF1 | 26034 | CLASH | 23622248 |
| hsa-let-7b-5p | MIMAT0000063 | PPP1R16B | 26051 | CLASH | 23622248 |
| hsa-let-7b-5p | MIMAT0000063 | DNM3 | 26052 | Proteomics | 18668040 |
| hsa-let-7b-5p | MIMAT0000063 | ANKRD17 | 26057 | CLASH | 23622248 |
| hsa-let-7b-5p | MIMAT0000063 | APPL1 | 26060 | CLASH | 23622248 |
| hsa-let-7b-5p | MIMAT0000063 | CHTOP | 26097 | HITS-CLIP | 23313552 |
| hsa-let-7b-5p | MIMAT0000063 | SZRD1 | 26099 | CLASH | 23622248 |
| hsa-let-7b-5p | MIMAT0000063 | GAPVD1 | 26130 | Proteomics | 18668040 |

| | | | | | |
|---|---|---|---|---|---|
| hsa-let-7b-5p | MIMAT0000063 | SERBP1 | 26135 | CLASH | 23622248 |
| hsa-let-7b-5p | MIMAT0000063 | TES | 26136 | Proteomics | 18668040 |
| hsa-let-7b-5p | MIMAT0000063 | INTS1 | 26173 | CLASH | 23622248 |
| hsa-let-7b-5p | MIMAT0000063 | FBXW2 | 26190 | CLASH//PAR-CLIP | 23622248\|23446348\|20371350 |
| hsa-let-7b-5p | MIMAT0000063 | MYCBP | 26292 | Proteomics | 18668040 |
| hsa-let-7b-5p | MIMAT0000063 | CNNM3 | 26505 | CLASH | 23622248 |
| hsa-let-7b-5p | MIMAT0000063 | TIMM9 | 26520 | Proteomics | 18668040 |
| hsa-let-7b-5p | MIMAT0000063 | AGO1 | 26523 | CLASH//Immunohistochemistry//Immunoprecipitaion//Luciferase reporter assay//Northern blot//qRT-PCR//Western blot | 23622248\|23426184 |
| hsa-let-7b-5p | MIMAT0000063 | AATF | 26574 | CLASH | 23622248 |
| hsa-let-7b-5p | MIMAT0000063 | CKAP2 | 26586 | CLASH | 23622248 |
| hsa-let-7b-5p | MIMAT0000063 | RANBP6 | 26953 | Proteomics | 18668040 |
| hsa-let-7b-5p | MIMAT0000063 | AP3M1 | 26985 | CLASH | 23622248 |
| hsa-let-7b-5p | MIMAT0000063 | PABPC1 | 26986 | CLASH | 23622248 |
| hsa-let-7b-5p | MIMAT0000063 | TRUB2 | 26995 | Proteomics | 18668040 |
| hsa-let-7b-5p | MIMAT0000063 | NPTN | 27020 | CLASH | 23622248 |
| hsa-let-7b-5p | MIMAT0000063 | VPS41 | 27072 | CLASH | 23622248 |
| hsa-let-7b-5p | MIMAT0000063 | EIF2AK1 | 27102 | CLASH | 23622248 |
| hsa-let-7b-5p | MIMAT0000063 | AFF4 | 27125 | CLASH | 23622248 |
| hsa-let-7b-5p | MIMAT0000063 | PALD1 | 27143 | CLASH | 23622248 |
| hsa-let-7b-5p | MIMAT0000063 | BRPF3 | 27154 | CLASH | 23622248 |
| hsa-let-7b-5p | MIMAT0000063 | AGO2 | 27161 | CLASH | 23622248 |
| hsa-let-7b-5p | MIMAT0000063 | SALL3 | 27164 | PAR-CLIP | 21572407 |
| hsa-let-7b-5p | MIMAT0000063 | DISC1 | 27185 | PAR-CLIP | 23592263\|26701625\|27292025 |
| hsa-let-7b-5p | MIMAT0000063 | CHMP2A | 27243 | Proteomics//pSILAC | 18668040 |
| hsa-let-7b-5p | MIMAT0000063 | RNF115 | 27246 | CLASH | 23622248 |
| hsa-let-7b-5p | MIMAT0000063 | NFU1 | 27247 | CLASH | 23622248 |
| hsa-let-7b-5p | MIMAT0000063 | TOX3 | 27324 | CLASH | 23622248 |
| hsa-let-7b-5p | MIMAT0000063 | PRPF19 | 27339 | CLASH | 23622248 |
| hsa-let-7b-5p | MIMAT0000063 | RRP7A | 27341 | Proteomics | 18668040 |
| hsa-let-7b-5p | MIMAT0000063 | POLL | 27343 | PAR-CLIP | 26701625 |
| hsa-let-7b-5p | MIMAT0000063 | MCAT | 27349 | Proteomics | 18668040 |
| hsa-let-7b-5p | MIMAT0000063 | MAT2B | 27430 | Proteomics | 18668040 |
| hsa-let-7b-5p | MIMAT0000063 | OSTM1 | 28962 | CLASH | 23622248 |
| hsa-let-7b-5p | MIMAT0000063 | BZW2 | 28969 | Proteomics | 18668040 |

| | | | | | |
|---|---|---|---|---|---|
| hsa-let-7b-5p | MIMAT0000063 | C19orf53 | 28974 | PAR-CLIP | 23592263\|26701625 |
| hsa-let-7b-5p | MIMAT0000063 | DBNL | 28988 | CLASH | 23622248 |
| hsa-let-7b-5p | MIMAT0000063 | THYN1 | 29087 | PAR-CLIP | 20371350 |
| hsa-let-7b-5p | MIMAT0000063 | COMMD9 | 29099 | Proteomics//pSILAC | 18668040 |
| hsa-let-7b-5p | MIMAT0000063 | UHRF1 | 29128 | Proteomics//pSILAC | 18668040 |
| hsa-let-7b-5p | MIMAT0000063 | ABT1 | 29777 | HITS-CLIP//PAR-CLIP | 23592263\|24398324\|21572407\|23313552\|27292025 |
| hsa-let-7b-5p | MIMAT0000063 | PARVB | 29780 | Proteomics | 18668040 |
| hsa-let-7b-5p | MIMAT0000063 | CPSF1 | 29894 | Proteomics | 18668040 |
| hsa-let-7b-5p | MIMAT0000063 | SNX12 | 29934 | CLASH | 23622248 |
| hsa-let-7b-5p | MIMAT0000063 | SLC25A24 | 29957 | Proteomics//pSILAC | 18668040 |
| hsa-let-7b-5p | MIMAT0000063 | NOP53 | 29997 | Proteomics | 18668040 |
| hsa-let-7b-5p | MIMAT0000063 | ERO1A | 30001 | PAR-CLIP//Proteomics | 18668040\|26701625 |
| hsa-let-7b-5p | MIMAT0000063 | SOCS7 | 30837 | PAR-CLIP | 26701625 |
| hsa-let-7b-5p | MIMAT0000063 | EHD4 | 30844 | Proteomics | 18668040 |
| hsa-let-7b-5p | MIMAT0000063 | TMED5 | 50999 | HITS-CLIP//PAR-CLIP//Proteomics | 18668040\|23592263\|24906430 |
| hsa-let-7b-5p | MIMAT0000063 | TRNT1 | 51095 | Proteomics | 18668040 |
| hsa-let-7b-5p | MIMAT0000063 | SH3GLB1 | 51100 | Proteomics | 18668040 |
| hsa-let-7b-5p | MIMAT0000063 | LACTB2 | 51110 | CLASH | 23622248 |
| hsa-let-7b-5p | MIMAT0000063 | NAA20 | 51126 | CLASH//PAR-CLIP | 23622248\|23592263 |
| hsa-let-7b-5p | MIMAT0000063 | RNFT1 | 51136 | PAR-CLIP | 24398324 |
| hsa-let-7b-5p | MIMAT0000063 | VPS28 | 51160 | CLASH | 23622248 |
| hsa-let-7b-5p | MIMAT0000063 | PLEKHO1 | 51177 | PAR-CLIP | 22291592\|20371350 |
| hsa-let-7b-5p | MIMAT0000063 | IPO11 | 51194 | CLASH | 23622248 |
| hsa-let-7b-5p | MIMAT0000063 | CPA4 | 51200 | HITS-CLIP | 23706177 |
| hsa-let-7b-5p | MIMAT0000063 | NUSAP1 | 51203 | CLASH | 23622248 |
| hsa-let-7b-5p | MIMAT0000063 | HGH1 | 51236 | pSILAC | 18668040 |
| hsa-let-7b-5p | MIMAT0000063 | MRPL37 | 51253 | CLASH | 23622248 |
| hsa-let-7b-5p | MIMAT0000063 | C1RL | 51279 | PAR-CLIP | 23592263 |
| hsa-let-7b-5p | MIMAT0000063 | ERGIC2 | 51290 | Proteomics | 18668040 |
| hsa-let-7b-5p | MIMAT0000063 | THEM6 | 51337 | PAR-CLIP | 23592263\|24398324 |
| hsa-let-7b-5p | MIMAT0000063 | RWDD1 | 51389 | HITS-CLIP | 23824327 |
| hsa-let-7b-5p | MIMAT0000063 | DDX41 | 51428 | Proteomics | 18668040 |
| hsa-let-7b-5p | MIMAT0000063 | SFMBT1 | 51460 | CLASH | 23622248 |
| hsa-let-7b-5p | MIMAT0000063 | NCKIPSD | 51517 | HITS-CLIP//PAR-CLIP | 23446348\|23706177 |
| hsa-let-7b-5p | MIMAT0000063 | TRMO | 51531 | PAR-CLIP | 23446348 |
| hsa-let-7b-5p | MIMAT0000063 | MTFP1 | 51537 | Proteomics | 18668040 |

| | | | | | |
|---|---|---|---|---|---|
| hsa-let-7b-5p | MIMAT0000063 | ZNF581 | 51545 | CLASH | 23622248 |
| hsa-let-7b-5p | MIMAT0000063 | FAM49B | 51571 | Proteomics | 18668040 |
| hsa-let-7b-5p | MIMAT0000063 | GDE1 | 51573 | Proteomics//pSILAC | 18668040 |
| hsa-let-7b-5p | MIMAT0000063 | ERGIC3 | 51614 | Proteomics | 18668040 |
| hsa-let-7b-5p | MIMAT0000063 | TAF9B | 51616 | Proteomics//pSILAC | 18668040 |
| hsa-let-7b-5p | MIMAT0000063 | MRPS33 | 51650 | Proteomics//pSILAC | 18668040 |
| hsa-let-7b-5p | MIMAT0000063 | CHMP3 | 51652 | Proteomics | 18668040 |
| hsa-let-7b-5p | MIMAT0000063 | WBP11 | 51729 | CLASH | 23622248 |
| hsa-let-7b-5p | MIMAT0000063 | FGFRL1 | 53834 | CLASH | 23622248 |
| hsa-let-7b-5p | MIMAT0000063 | CHRAC1 | 54108 | Proteomics | 18668040 |
| hsa-let-7b-5p | MIMAT0000063 | TERF2IP | 54386 | CLASH | 23622248 |
| hsa-let-7b-5p | MIMAT0000063 | SLC38A2 | 54407 | Proteomics | 18668040 |
| hsa-let-7b-5p | MIMAT0000063 | XRN1 | 54464 | Proteomics | 18668040 |
| hsa-let-7b-5p | MIMAT0000063 | MIOS | 54468 | Proteomics | 18668040 |
| hsa-let-7b-5p | MIMAT0000063 | MIEF1 | 54471 | PAR-CLIP | 23592263\|24398324\|21572407\|20371350 |
| hsa-let-7b-5p | MIMAT0000063 | NLE1 | 54475 | Proteomics | 18668040 |
| hsa-let-7b-5p | MIMAT0000063 | CHPF2 | 54480 | Proteomics//pSILAC | 18668040 |
| hsa-let-7b-5p | MIMAT0000063 | FAM105A | 54491 | HITS-CLIP//PAR-CLIP//Proteomics//pSILAC | 18668040\|23592263\|23446348\|23706177 |
| hsa-let-7b-5p | MIMAT0000063 | MIER2 | 54531 | CLASH | 23622248 |
| hsa-let-7b-5p | MIMAT0000063 | DDX49 | 54555 | Proteomics | 18668040 |
| hsa-let-7b-5p | MIMAT0000063 | CCNJ | 54619 | Microarray | 17699775 |
| hsa-let-7b-5p | MIMAT0000063 | ARL15 | 54622 | Proteomics//pSILAC | 18668040 |
| hsa-let-7b-5p | MIMAT0000063 | TBC1D13 | 54662 | Proteomics | 18668040 |
| hsa-let-7b-5p | MIMAT0000063 | WDR74 | 54663 | Proteomics | 18668040 |
| hsa-let-7b-5p | MIMAT0000063 | MANSC1 | 54682 | CLASH | 23622248 |
| hsa-let-7b-5p | MIMAT0000063 | RBFOX1 | 54715 | Proteomics | 18668040 |
| hsa-let-7b-5p | MIMAT0000063 | PPP1R12C | 54776 | CLASH | 23622248 |
| hsa-let-7b-5p | MIMAT0000063 | NSMCE4A | 54780 | Proteomics | 18668040 |
| hsa-let-7b-5p | MIMAT0000063 | HAUS6 | 54801 | CLASH | 23622248 |
| hsa-let-7b-5p | MIMAT0000063 | WDR55 | 54853 | CLASH | 23622248 |
| hsa-let-7b-5p | MIMAT0000063 | TOR4A | 54863 | CLASH | 23622248 |
| hsa-let-7b-5p | MIMAT0000063 | GPATCH4 | 54865 | Proteomics | 18668040 |
| hsa-let-7b-5p | MIMAT0000063 | PIGG | 54872 | CLASH | 23622248 |
| hsa-let-7b-5p | MIMAT0000063 | BCOR | 54880 | CLASH | 23622248 |
| hsa-let-7b-5p | MIMAT0000063 | NSUN2 | 54888 | CLASH | 23622248 |

| | | | | | |
|---|---|---|---|---|---|
| hsa-let-7b-5p | MIMAT0000063 | NCAPG2 | 54892 | Proteomics//pSILAC | 18668040 |
| hsa-let-7b-5p | MIMAT0000063 | CDKAL1 | 54901 | HITS-CLIP//Proteomics//pSILAC | 18668040\|23706177 |
| hsa-let-7b-5p | MIMAT0000063 | SEMA4C | 54910 | PAR-CLIP | 24398324\|23446348\|20371350 |
| hsa-let-7b-5p | MIMAT0000063 | DNAAF5 | 54919 | Proteomics | 18668040 |
| hsa-let-7b-5p | MIMAT0000063 | DUSP23 | 54935 | Proteomics//pSILAC | 18668040 |
| hsa-let-7b-5p | MIMAT0000063 | DNAJC28 | 54943 | HITS-CLIP | 23313552 |
| hsa-let-7b-5p | MIMAT0000063 | C1orf27 | 54953 | Proteomics//pSILAC | 18668040 |
| hsa-let-7b-5p | MIMAT0000063 | PARP16 | 54956 | PAR-CLIP | 20371350 |
| hsa-let-7b-5p | MIMAT0000063 | INTS11 | 54973 | Proteomics | 18668040 |
| hsa-let-7b-5p | MIMAT0000063 | SLC35F6 | 54978 | Proteomics//pSILAC | 18668040 |
| hsa-let-7b-5p | MIMAT0000063 | PIH1D1 | 55011 | Proteomics | 18668040 |
| hsa-let-7b-5p | MIMAT0000063 | USP47 | 55031 | PAR-CLIP | 27292025 |
| hsa-let-7b-5p | MIMAT0000063 | PDPR | 55066 | Proteomics | 18668040 |
| hsa-let-7b-5p | MIMAT0000063 | OXR1 | 55074 | CLASH | 23622248 |
| hsa-let-7b-5p | MIMAT0000063 | CCDC186 | 55088 | CLASH | 23622248 |
| hsa-let-7b-5p | MIMAT0000063 | GPATCH1 | 55094 | CLASH | 23622248 |
| hsa-let-7b-5p | MIMAT0000063 | ARHGAP17 | 55114 | Proteomics | 18668040 |
| hsa-let-7b-5p | MIMAT0000063 | FIGN | 55137 | HITS-CLIP//PAR-CLIP | 23446348\|21572407\|20371350\|23706177 |
| hsa-let-7b-5p | MIMAT0000063 | ANKZF1 | 55139 | Proteomics | 18668040 |
| hsa-let-7b-5p | MIMAT0000063 | CDCA8 | 55143 | Proteomics | 18668040 |
| hsa-let-7b-5p | MIMAT0000063 | SDAD1 | 55153 | Proteomics | 18668040 |
| hsa-let-7b-5p | MIMAT0000063 | TMEM33 | 55161 | Proteomics | 18668040 |
| hsa-let-7b-5p | MIMAT0000063 | KLHL11 | 55175 | CLASH | 23622248 |
| hsa-let-7b-5p | MIMAT0000063 | MRM3 | 55178 | Proteomics | 18668040 |
| hsa-let-7b-5p | MIMAT0000063 | ARL8B | 55207 | HITS-CLIP | 23706177 |
| hsa-let-7b-5p | MIMAT0000063 | SETD5 | 55209 | CLASH | 23622248 |
| hsa-let-7b-5p | MIMAT0000063 | BBS7 | 55212 | CLASH | 23622248 |
| hsa-let-7b-5p | MIMAT0000063 | C11orf57 | 55216 | PAR-CLIP | 23592263\|21572407 |
| hsa-let-7b-5p | MIMAT0000063 | LRRC20 | 55222 | PAR-CLIP | 23592263 |
| hsa-let-7b-5p | MIMAT0000063 | MOB1A | 55233 | Proteomics | 18668040 |
| hsa-let-7b-5p | MIMAT0000063 | SLC38A7 | 55238 | HITS-CLIP | 23706177 |
| hsa-let-7b-5p | MIMAT0000063 | STEAP3 | 55240 | CLASH | 23622248 |
| hsa-let-7b-5p | MIMAT0000063 | QRSL1 | 55278 | Proteomics | 18668040 |
| hsa-let-7b-5p | MIMAT0000063 | TBC1D19 | 55296 | PAR-CLIP | 22291592 |
| hsa-let-7b-5p | MIMAT0000063 | SYNJ2BP | 55333 | PAR-CLIP | 23592263\|26701625 |

| | | | | | |
|---|---|---|---|---|---|
| hsa-let-7b-5p | MIMAT0000063 | WDR33 | 55339 | Proteomics | 18668040 |
| hsa-let-7b-5p | MIMAT0000063 | LSG1 | 55341 | Proteomics | 18668040 |
| hsa-let-7b-5p | MIMAT0000063 | TMEM63B | 55362 | CLASH | 23622248 |
| hsa-let-7b-5p | MIMAT0000063 | LGR4 | 55366 | Flow//Luciferase reporter assay//qRT-PCR//Western blot | 27179410 |
| hsa-let-7b-5p | MIMAT0000063 | YOD1 | 55432 | PAR-CLIP | 23446348\|21572407\|20371350 |
| hsa-let-7b-5p | MIMAT0000063 | NDUFAF7 | 55471 | Proteomics | 18668040 |
| hsa-let-7b-5p | MIMAT0000063 | DHTKD1 | 55526 | CLASH | 23622248 |
| hsa-let-7b-5p | MIMAT0000063 | FOXRED1 | 55572 | Proteomics | 18668040 |
| hsa-let-7b-5p | MIMAT0000063 | CDV3 | 55573 | PAR-CLIP | 21572407 |
| hsa-let-7b-5p | MIMAT0000063 | SUPT20H | 55578 | CLASH | 23622248 |
| hsa-let-7b-5p | MIMAT0000063 | KIF27 | 55582 | PAR-CLIP | 22100165 |
| hsa-let-7b-5p | MIMAT0000063 | UBE2Q1 | 55585 | CLASH | 23622248 |
| hsa-let-7b-5p | MIMAT0000063 | OTUB1 | 55611 | CLASH | 23622248 |
| hsa-let-7b-5p | MIMAT0000063 | TRMT1 | 55621 | Proteomics//pSILAC | 18668040 |
| hsa-let-7b-5p | MIMAT0000063 | LRRC40 | 55631 | Proteomics | 18668040 |
| hsa-let-7b-5p | MIMAT0000063 | CHD7 | 55636 | CLASH | 23622248 |
| hsa-let-7b-5p | MIMAT0000063 | BCAS4 | 55653 | CLASH | 23622248 |
| hsa-let-7b-5p | MIMAT0000063 | YEATS2 | 55689 | CLASH | 23622248 |
| hsa-let-7b-5p | MIMAT0000063 | RBM22 | 55696 | CLASH | 23622248 |
| hsa-let-7b-5p | MIMAT0000063 | IARS2 | 55699 | CLASH | 23622248 |
| hsa-let-7b-5p | MIMAT0000063 | MAP7D1 | 55700 | Proteomics//pSILAC | 18668040 |
| hsa-let-7b-5p | MIMAT0000063 | POLR3B | 55703 | Proteomics | 18668040 |
| hsa-let-7b-5p | MIMAT0000063 | IPO9 | 55705 | HITS-CLIP | 23313552 |
| hsa-let-7b-5p | MIMAT0000063 | TENM3 | 55714 | CLASH | 23622248 |
| hsa-let-7b-5p | MIMAT0000063 | SLF2 | 55719 | CLASH | 23622248 |
| hsa-let-7b-5p | MIMAT0000063 | FAM222B | 55731 | PAR-CLIP | 21572407 |
| hsa-let-7b-5p | MIMAT0000063 | DNAJC11 | 55735 | Proteomics | 18668040 |
| hsa-let-7b-5p | MIMAT0000063 | CNDP2 | 55748 | Proteomics | 18668040 |
| hsa-let-7b-5p | MIMAT0000063 | RIOK2 | 55781 | Proteomics | 18668040 |
| hsa-let-7b-5p | MIMAT0000063 | TXLNG | 55787 | PAR-CLIP | 21572407 |
| hsa-let-7b-5p | MIMAT0000063 | DDX28 | 55794 | Proteomics | 18668040 |
| hsa-let-7b-5p | MIMAT0000063 | UTP6 | 55813 | Proteomics | 18668040 |
| hsa-let-7b-5p | MIMAT0000063 | DBNDD2 | 55861 | CLASH | 23622248 |
| hsa-let-7b-5p | MIMAT0000063 | ECHDC1 | 55862 | PAR-CLIP | 21572407 |
| hsa-let-7b-5p | MIMAT0000063 | NXT2 | 55916 | Microarray | 17699775 |
| hsa-let-7b-5p | MIMAT0000063 | YLPM1 | 56252 | CLASH | 23622248 |

| | | | | | |
|---|---|---|---|---|---|
| hsa-let-7b-5p | MIMAT0000063 | LRRC8A | 56262 | Proteomics | 18668040 |
| hsa-let-7b-5p | MIMAT0000063 | KYAT3 | 56267 | Proteomics//pSILAC | 18668040 |
| hsa-let-7b-5p | MIMAT0000063 | DIABLO | 56616 | PAR-CLIP | 23446348\|20371350 |
| hsa-let-7b-5p | MIMAT0000063 | SAR1A | 56681 | HITS-CLIP | 23706177 |
| hsa-let-7b-5p | MIMAT0000063 | ZC3HAV1 | 56829 | CLASH | 23622248 |
| hsa-let-7b-5p | MIMAT0000063 | RAD18 | 56852 | PAR-CLIP | 21572407 |
| hsa-let-7b-5p | MIMAT0000063 | DPYSL5 | 56896 | CLASH | 23622248 |
| hsa-let-7b-5p | MIMAT0000063 | C1GALT1 | 56913 | CLASH | 23622248 |
| hsa-let-7b-5p | MIMAT0000063 | SMARCAD1 | 56916 | PAR-CLIP | 24398324\|20371350 |
| hsa-let-7b-5p | MIMAT0000063 | DHX33 | 56919 | Proteomics | 18668040 |
| hsa-let-7b-5p | MIMAT0000063 | C5orf15 | 56951 | CLASH | 23622248 |
| hsa-let-7b-5p | MIMAT0000063 | ATXN7L3 | 56970 | PAR-CLIP | 23592263 |
| hsa-let-7b-5p | MIMAT0000063 | YAE1D1 | 57002 | PAR-CLIP | 23592263 |
| hsa-let-7b-5p | MIMAT0000063 | PITHD1 | 57095 | Proteomics | 18668040 |
| hsa-let-7b-5p | MIMAT0000063 | C12orf4 | 57102 | PAR-CLIP | 23446348 |
| hsa-let-7b-5p | MIMAT0000063 | MRPL47 | 57129 | Proteomics | 18668040 |
| hsa-let-7b-5p | MIMAT0000063 | RALGAPB | 57148 | CLASH | 23622248 |
| hsa-let-7b-5p | MIMAT0000063 | SPRYD7 | 57213 | Proteomics | 18668040 |
| hsa-let-7b-5p | MIMAT0000063 | NHSL1 | 57224 | CLASH | 23622248 |
| hsa-let-7b-5p | MIMAT0000063 | SCYL1 | 57410 | Proteomics//pSILAC | 18668040 |
| hsa-let-7b-5p | MIMAT0000063 | BIRC6 | 57448 | qRT-PCR | 17942906 |
| hsa-let-7b-5p | MIMAT0000063 | KIAA1143 | 57456 | PAR-CLIP | 26701625 |
| hsa-let-7b-5p | MIMAT0000063 | SCAF4 | 57466 | Proteomics | 18668040 |
| hsa-let-7b-5p | MIMAT0000063 | LRRC47 | 57470 | Proteomics | 18668040 |
| hsa-let-7b-5p | MIMAT0000063 | KIDINS220 | 57498 | Proteomics | 18668040 |
| hsa-let-7b-5p | MIMAT0000063 | MTUS1 | 57509 | CLASH//PAR-CLIP | 23622248\|23592263\|24398324\|23446348\|26701625 |
| hsa-let-7b-5p | MIMAT0000063 | XPO5 | 57510 | Proteomics | 18668040 |
| hsa-let-7b-5p | MIMAT0000063 | MIB1 | 57534 | CLASH | 23622248 |
| hsa-let-7b-5p | MIMAT0000063 | KIAA1328 | 57536 | HITS-CLIP | 23313552 |
| hsa-let-7b-5p | MIMAT0000063 | PDP2 | 57546 | CLASH//PAR-CLIP | 23622248\|23592263 |
| hsa-let-7b-5p | MIMAT0000063 | CEP126 | 57562 | CLASH | 23622248 |
| hsa-let-7b-5p | MIMAT0000063 | ZNF687 | 57592 | CLASH | 23622248 |
| hsa-let-7b-5p | MIMAT0000063 | KIAA1549 | 57670 | CLASH | 23622248 |
| hsa-let-7b-5p | MIMAT0000063 | KIAA1586 | 57691 | CLASH | 23622248 |
| hsa-let-7b-5p | MIMAT0000063 | ZNF317 | 57693 | CLASH | 23622248 |

| | | | | | |
|---|---|---|---|---|---|
| hsa-let-7b-5p | MIMAT0000063 | IGDCC4 | 57722 | PAR-CLIP | 23446348\|21572407\|20371350\|26701625 |
| hsa-let-7b-5p | MIMAT0000063 | NCOA5 | 57727 | CLASH | 23622248 |
| hsa-let-7b-5p | MIMAT0000063 | RAB40C | 57799 | PAR-CLIP | 23592263 |
| hsa-let-7b-5p | MIMAT0000063 | RAP2C | 57826 | Proteomics | 18668040 |
| hsa-let-7b-5p | MIMAT0000063 | TRAPPC1 | 58485 | CLASH | 23622248 |
| hsa-let-7b-5p | MIMAT0000063 | ABHD17C | 58489 | PAR-CLIP | 20371350 |
| hsa-let-7b-5p | MIMAT0000063 | TLNRD1 | 59274 | CLASH | 23622248 |
| hsa-let-7b-5p | MIMAT0000063 | CACNG8 | 59283 | CLASH | 23622248 |
| hsa-let-7b-5p | MIMAT0000063 | SLC25A19 | 60386 | Proteomics | 18668040 |
| hsa-let-7b-5p | MIMAT0000063 | SPCS3 | 60559 | Proteomics//pSILAC | 18668040 |
| hsa-let-7b-5p | MIMAT0000063 | BCORL1 | 63035 | CLASH | 23622248 |
| hsa-let-7b-5p | MIMAT0000063 | GALNT11 | 63917 | CLASH | 23622248 |
| hsa-let-7b-5p | MIMAT0000063 | PRSS22 | 64063 | HITS-CLIP | 23313552 |
| hsa-let-7b-5p | MIMAT0000063 | XYLT2 | 64132 | CLASH | 23622248 |
| hsa-let-7b-5p | MIMAT0000063 | ERAP2 | 64167 | CLASH | 23622248 |
| hsa-let-7b-5p | MIMAT0000063 | DNAJC1 | 64215 | Proteomics | 18668040 |
| hsa-let-7b-5p | MIMAT0000063 | NSD1 | 64324 | PAR-CLIP | 24398324 |
| hsa-let-7b-5p | MIMAT0000063 | NXN | 64359 | Proteomics//pSILAC | 18668040 |
| hsa-let-7b-5p | MIMAT0000063 | ZNF106 | 64397 | CLASH | 23622248 |
| hsa-let-7b-5p | MIMAT0000063 | GZF1 | 64412 | Microarray | 17699775 |
| hsa-let-7b-5p | MIMAT0000063 | NOM1 | 64434 | PAR-CLIP | 23592263\|23446348 |
| hsa-let-7b-5p | MIMAT0000063 | CPEB1 | 64506 | Luciferase reporter assay//Microarray//qRT-PCR | 22995917 |
| hsa-let-7b-5p | MIMAT0000063 | ANAPC1 | 64682 | Microarray//Proteomics//pSILAC//qRT-PCR//Western blot | 18668040\|26540468 |
| hsa-let-7b-5p | MIMAT0000063 | NUCKS1 | 64710 | CLASH | 23622248 |
| hsa-let-7b-5p | MIMAT0000063 | TBC1D15 | 64786 | Proteomics | 18668040 |
| hsa-let-7b-5p | MIMAT0000063 | ELOVL1 | 64834 | Proteomics | 18668040 |
| hsa-let-7b-5p | MIMAT0000063 | TUT1 | 64852 | CLASH | 23622248 |
| hsa-let-7b-5p | MIMAT0000063 | AIDA | 64853 | CLASH | 23622248 |
| hsa-let-7b-5p | MIMAT0000063 | CCDC71 | 64925 | CLASH | 23622248 |
| hsa-let-7b-5p | MIMAT0000063 | NT5DC2 | 64943 | CLASH | 23622248 |
| hsa-let-7b-5p | MIMAT0000063 | MRPS24 | 64951 | Proteomics//pSILAC | 18668040 |
| hsa-let-7b-5p | MIMAT0000063 | MRPS11 | 64963 | CLASH | 23622248 |
| hsa-let-7b-5p | MIMAT0000063 | NOL6 | 65083 | Proteomics | 18668040 |

| | | | | | |
|---|---|---|---|---|---|
| hsa-let-7b-5p | MIMAT0000063 | MARCKS L1 | 65108 | PAR-CLIP | 26701625 |
| hsa-let-7b-5p | MIMAT0000063 | WNK1 | 65125 | CLASH | 23622248 |
| hsa-let-7b-5p | MIMAT0000063 | PYCR3 | 65263 | Proteomics | 18668040 |
| hsa-let-7b-5p | MIMAT0000063 | PLEKHA3 | 65977 | PAR-CLIP | 21572407 |
| hsa-let-7b-5p | MIMAT0000063 | PHACTR 4 | 65979 | HITS-CLIP | 23706177 |
| hsa-let-7b-5p | MIMAT0000063 | GGCT | 79017 | Proteomics//pSILAC | 18668040 |
| hsa-let-7b-5p | MIMAT0000063 | PDCL3 | 79031 | CLASH | 23622248 |
| hsa-let-7b-5p | MIMAT0000063 | NABP2 | 79035 | CLASH | 23622248 |
| hsa-let-7b-5p | MIMAT0000063 | KXD1 | 79036 | CLASH | 23622248 |
| hsa-let-7b-5p | MIMAT0000063 | ASPSCR1 | 79058 | CLASH | 23622248 |
| hsa-let-7b-5p | MIMAT0000063 | ATG9A | 79065 | PAR-CLIP//Proteomics | 18668040\|23592263\|26701625\|27292025 |
| hsa-let-7b-5p | MIMAT0000063 | FTO | 79068 | CLASH | 23622248 |
| hsa-let-7b-5p | MIMAT0000063 | DCTPP1 | 79077 | Proteomics//pSILAC | 18668040 |
| hsa-let-7b-5p | MIMAT0000063 | ZNF426 | 79088 | CLASH | 23622248 |
| hsa-let-7b-5p | MIMAT0000063 | EFHD2 | 79180 | PAR-CLIP | 23592263 |
| hsa-let-7b-5p | MIMAT0000063 | WDR25 | 79446 | CLASH | 23622248 |
| hsa-let-7b-5p | MIMAT0000063 | ADIPOR2 | 79602 | CLASH//PAR-CLIP | 23622248\|21572407 |
| hsa-let-7b-5p | MIMAT0000063 | RHBDF2 | 79651 | PAR-CLIP | 26701625 |
| hsa-let-7b-5p | MIMAT0000063 | MSANTD 2 | 79684 | CLASH | 23622248 |
| hsa-let-7b-5p | MIMAT0000063 | IPO4 | 79711 | CLASH//Proteomics//pSILAC | 18668040\|23622248 |
| hsa-let-7b-5p | MIMAT0000063 | LIN28A | 79727 | Luciferase reporter assay//Reporter assay | 15131085 |
| hsa-let-7b-5p | MIMAT0000063 | NARS2 | 79731 | Proteomics | 18668040 |
| hsa-let-7b-5p | MIMAT0000063 | GEMIN7 | 79760 | Proteomics//pSILAC | 18668040 |
| hsa-let-7b-5p | MIMAT0000063 | ISOC2 | 79763 | CLASH | 23622248 |
| hsa-let-7b-5p | MIMAT0000063 | ZFHX4 | 79776 | CLASH | 23622248 |
| hsa-let-7b-5p | MIMAT0000063 | C12orf4 9 | 79794 | CLASH | 23622248 |
| hsa-let-7b-5p | MIMAT0000063 | NAA40 | 79829 | Proteomics//pSILAC | 18668040 |
| hsa-let-7b-5p | MIMAT0000063 | FAM57A | 79850 | CLASH | 23622248 |
| hsa-let-7b-5p | MIMAT0000063 | CCDC13 4 | 79879 | Proteomics | 18668040 |
| hsa-let-7b-5p | MIMAT0000063 | MRM1 | 79922 | Proteomics//pSILAC | 18668040 |
| hsa-let-7b-5p | MIMAT0000063 | WLS | 79971 | Proteomics | 18668040 |
| hsa-let-7b-5p | MIMAT0000063 | CPED1 | 79974 | Microarray | 17699775 |
| hsa-let-7b-5p | MIMAT0000063 | DOCK5 | 80005 | Proteomics//pSILAC | 18668040 |
| hsa-let-7b-5p | MIMAT0000063 | NAA25 | 80018 | Proteomics//pSILAC | 18668040 |
| hsa-let-7b-5p | MIMAT0000063 | FOXRED 2 | 80020 | CLASH | 23622248 |

| | | | | | |
|---|---|---|---|---|---|
| hsa-let-7b-5p | MIMAT0000063 | ZNF556 | 80032 | HITS-CLIP | 23824327 |
| hsa-let-7b-5p | MIMAT0000063 | ZNF606 | 80095 | CLASH | 23622248 |
| hsa-let-7b-5p | MIMAT0000063 | PTGES2 | 80142 | CLASH | 23622248 |
| hsa-let-7b-5p | MIMAT0000063 | SIKE1 | 80143 | CLASH | 23622248 |
| hsa-let-7b-5p | MIMAT0000063 | NAA15 | 80155 | Proteomics | 18668040 |
| hsa-let-7b-5p | MIMAT0000063 | OPA3 | 80207 | PAR-CLIP | 23592263\|26701625 |
| hsa-let-7b-5p | MIMAT0000063 | NAA50 | 80218 | Proteomics | 18668040 |
| hsa-let-7b-5p | MIMAT0000063 | PAAF1 | 80227 | Proteomics | 18668040 |
| hsa-let-7b-5p | MIMAT0000063 | WDR26 | 80232 | CLASH | 23622248 |
| hsa-let-7b-5p | MIMAT0000063 | EDEM3 | 80267 | HITS-CLIP//Proteomics//pSILAC | 18668040\|23313552 |
| hsa-let-7b-5p | MIMAT0000063 | WDCP | 80304 | CLASH | 23622248 |
| hsa-let-7b-5p | MIMAT0000063 | TRABD | 80305 | CLASH//Proteomics//pSILAC | 18668040\|23622248 |
| hsa-let-7b-5p | MIMAT0000063 | MED28 | 80306 | CLASH | 23622248 |
| hsa-let-7b-5p | MIMAT0000063 | FLAD1 | 80308 | CLASH//Proteomics | 18668040\|23622248 |
| hsa-let-7b-5p | MIMAT0000063 | CPEB4 | 80315 | Luciferase reporter assay//Microarray//qRT-PCR | 22995917 |
| hsa-let-7b-5p | MIMAT0000063 | PUS1 | 80324 | Proteomics | 18668040 |
| hsa-let-7b-5p | MIMAT0000063 | REEP4 | 80346 | Proteomics | 18668040 |
| hsa-let-7b-5p | MIMAT0000063 | SLC19A3 | 80704 | HITS-CLIP | 23706177 |
| hsa-let-7b-5p | MIMAT0000063 | AARSD1 | 80755 | Proteomics//pSILAC | 18668040 |
| hsa-let-7b-5p | MIMAT0000063 | CPTP | 80772 | CLASH | 23622248 |
| hsa-let-7b-5p | MIMAT0000063 | LIMD2 | 80774 | PAR-CLIP | 23592263 |
| hsa-let-7b-5p | MIMAT0000063 | INTS5 | 80789 | Proteomics | 18668040 |
| hsa-let-7b-5p | MIMAT0000063 | SLC25A32 | 81034 | Proteomics//pSILAC | 18668040 |
| hsa-let-7b-5p | MIMAT0000063 | COLEC12 | 81035 | PAR-CLIP | 21572407 |
| hsa-let-7b-5p | MIMAT0000063 | SLC38A1 | 81539 | Proteomics | 18668040 |
| hsa-let-7b-5p | MIMAT0000063 | GDPD5 | 81544 | CLASH | 23622248 |
| hsa-let-7b-5p | MIMAT0000063 | RCC1L | 81554 | Proteomics | 18668040 |
| hsa-let-7b-5p | MIMAT0000063 | INTS14 | 81556 | Proteomics | 18668040 |
| hsa-let-7b-5p | MIMAT0000063 | C1orf21 | 81563 | PAR-CLIP | 20371350 |
| hsa-let-7b-5p | MIMAT0000063 | ANP32E | 81611 | Proteomics | 18668040 |
| hsa-let-7b-5p | MIMAT0000063 | C6orf62 | 81688 | CLASH | 23622248 |
| hsa-let-7b-5p | MIMAT0000063 | ZNF611 | 81856 | HITS-CLIP//PAR-CLIP | 21572407\|20371350\|23706177 |
| hsa-let-7b-5p | MIMAT0000063 | MED25 | 81857 | CLASH | 23622248 |
| hsa-let-7b-5p | MIMAT0000063 | EMC6 | 83460 | Proteomics | 18668040 |

| | | | | | |
|---|---|---|---|---|---|
| hsa-let-7b-5p | MIMAT0000063 | DNAL1 | 83544 | PAR-CLIP | 21572407\|27292025 |
| hsa-let-7b-5p | MIMAT0000063 | UCK1 | 83549 | CLASH | 23622248 |
| hsa-let-7b-5p | MIMAT0000063 | ATAD3B | 83858 | Proteomics//pSILAC | 18668040 |
| hsa-let-7b-5p | MIMAT0000063 | CDCA7 | 83879 | Microarray | 17699775 |
| hsa-let-7b-5p | MIMAT0000063 | HASPIN | 83903 | HITS-CLIP//PAR-CLIP | 21572407\|20371350\|23706177 |
| hsa-let-7b-5p | MIMAT0000063 | KREMEN1 | 83999 | PAR-CLIP | 23592263\|26701625 |
| hsa-let-7b-5p | MIMAT0000063 | EMILIN2 | 84034 | HITS-CLIP//PAR-CLIP | 20371350\|23706177 |
| hsa-let-7b-5p | MIMAT0000063 | SLC10A7 | 84068 | PAR-CLIP | 24398324\|21572407\|20371350 |
| hsa-let-7b-5p | MIMAT0000063 | WDR75 | 84128 | Proteomics | 18668040 |
| hsa-let-7b-5p | MIMAT0000063 | TOMM40L | 84134 | PAR-CLIP | 26701625\|27292025 |
| hsa-let-7b-5p | MIMAT0000063 | UTP15 | 84135 | Proteomics | 18668040 |
| hsa-let-7b-5p | MIMAT0000063 | ZNF644 | 84146 | PAR-CLIP | 23446348 |
| hsa-let-7b-5p | MIMAT0000063 | ANTXR1 | 84168 | CLASH | 23622248 |
| hsa-let-7b-5p | MIMAT0000063 | POLR1B | 84172 | Proteomics | 18668040 |
| hsa-let-7b-5p | MIMAT0000063 | FAM96A | 84191 | Proteomics//pSILAC | 18668040 |
| hsa-let-7b-5p | MIMAT0000063 | ZCCHC9 | 84240 | CLASH | 23622248 |
| hsa-let-7b-5p | MIMAT0000063 | NOA1 | 84273 | PAR-CLIP | 23592263\|23446348 |
| hsa-let-7b-5p | MIMAT0000063 | FAM213A | 84293 | Proteomics | 18668040 |
| hsa-let-7b-5p | MIMAT0000063 | LLPH | 84298 | Proteomics | 18668040 |
| hsa-let-7b-5p | MIMAT0000063 | MIEN1 | 84299 | CLASH | 23622248 |
| hsa-let-7b-5p | MIMAT0000063 | CCDC115 | 84317 | Proteomics//pSILAC | 18668040 |
| hsa-let-7b-5p | MIMAT0000063 | LZIC | 84328 | Proteomics | 18668040 |
| hsa-let-7b-5p | MIMAT0000063 | GFM2 | 84340 | Proteomics | 18668040 |
| hsa-let-7b-5p | MIMAT0000063 | CYSTM1 | 84418 | CLASH | 23622248 |
| hsa-let-7b-5p | MIMAT0000063 | RAB11FIP4 | 84440 | PAR-CLIP | 24398324 |
| hsa-let-7b-5p | MIMAT0000063 | ZBTB37 | 84614 | PAR-CLIP | 20371350 |
| hsa-let-7b-5p | MIMAT0000063 | USP38 | 84640 | PAR-CLIP | 21572407 |
| hsa-let-7b-5p | MIMAT0000063 | FAM126A | 84668 | CLASH | 23622248 |
| hsa-let-7b-5p | MIMAT0000063 | GTPBP3 | 84705 | Proteomics//pSILAC | 18668040 |
| hsa-let-7b-5p | MIMAT0000063 | PRRC2B | 84726 | CLASH | 23622248 |
| hsa-let-7b-5p | MIMAT0000063 | FUT10 | 84750 | PAR-CLIP | 23592263\|26701625 |
| hsa-let-7b-5p | MIMAT0000063 | TUBA1C | 84790 | CLASH | 23622248 |
| hsa-let-7b-5p | MIMAT0000063 | IL17RC | 84818 | CLASH | 23622248 |
| hsa-let-7b-5p | MIMAT0000063 | PLXDC2 | 84898 | Proteomics | 18668040 |
| hsa-let-7b-5p | MIMAT0000063 | FAM136A | 84908 | CLASH | 23622248 |
| hsa-let-7b-5p | MIMAT0000063 | ZNF587 | 84914 | PAR-CLIP | 23592263 |
| hsa-let-7b-5p | MIMAT0000063 | PPP1R15B | 84919 | PAR-CLIP | 23592263\|27292025 |

| | | | | | |
|---|---|---|---|---|---|
| hsa-let-7b-5p | MIMAT0000063 | FAM104A | 84923 | PAR-CLIP | 20371350 |
| hsa-let-7b-5p | MIMAT0000063 | ZNF566 | 84924 | PAR-CLIP | 23446348\|21572407\|20371350 |
| hsa-let-7b-5p | MIMAT0000063 | MPND | 84954 | CLASH | 23622248 |
| hsa-let-7b-5p | MIMAT0000063 | FBXL20 | 84961 | PAR-CLIP | 21572407 |
| hsa-let-7b-5p | MIMAT0000063 | REPS1 | 85021 | CLASH | 23622248 |
| hsa-let-7b-5p | MIMAT0000063 | HIST1H2BK | 85236 | PAR-CLIP | 23592263\|27292025 |
| hsa-let-7b-5p | MIMAT0000063 | ZCCHC3 | 85364 | PAR-CLIP | 23446348\|26701625 |
| hsa-let-7b-5p | MIMAT0000063 | EAF1 | 85403 | CLASH | 23622248 |
| hsa-let-7b-5p | MIMAT0000063 | MIDN | 90007 | CLASH//PAR-CLIP | 23622248\|26701625 |
| hsa-let-7b-5p | MIMAT0000063 | ZNF799 | 90576 | HITS-CLIP | 23313552 |
| hsa-let-7b-5p | MIMAT0000063 | PIP4P1 | 90809 | CLASH | 23622248 |
| hsa-let-7b-5p | MIMAT0000063 | ADCK2 | 90956 | CLASH | 23622248 |
| hsa-let-7b-5p | MIMAT0000063 | DHX57 | 90957 | CLASH//Proteomics//pSILAC | 18668040\|23622248 |
| hsa-let-7b-5p | MIMAT0000063 | FMNL3 | 91010 | PAR-CLIP | 22100165 |
| hsa-let-7b-5p | MIMAT0000063 | L3MBTL4 | 91133 | CLASH | 23622248 |
| hsa-let-7b-5p | MIMAT0000063 | YTHDC1 | 91746 | CLASH | 23622248 |
| hsa-let-7b-5p | MIMAT0000063 | NEK9 | 91754 | CLASH | 23622248 |
| hsa-let-7b-5p | MIMAT0000063 | C11orf52 | 91894 | CLASH | 23622248 |
| hsa-let-7b-5p | MIMAT0000063 | ZC3HAV1L | 92092 | PAR-CLIP | 23592263 |
| hsa-let-7b-5p | MIMAT0000063 | OXNAD1 | 92106 | Proteomics | 18668040 |
| hsa-let-7b-5p | MIMAT0000063 | MTSS1L | 92154 | CLASH | 23622248 |
| hsa-let-7b-5p | MIMAT0000063 | NAF1 | 92345 | Proteomics | 18668040 |
| hsa-let-7b-5p | MIMAT0000063 | MOB1B | 92597 | Proteomics | 18668040 |
| hsa-let-7b-5p | MIMAT0000063 | TIMM50 | 92609 | CLASH | 23622248 |
| hsa-let-7b-5p | MIMAT0000063 | MGME1 | 92667 | Proteomics//pSILAC | 18668040 |
| hsa-let-7b-5p | MIMAT0000063 | SLC38A5 | 92745 | Proteomics | 18668040 |
| hsa-let-7b-5p | MIMAT0000063 | MARS2 | 92935 | PAR-CLIP//pSILAC | 18668040\|26701625 |
| hsa-let-7b-5p | MIMAT0000063 | IGSF8 | 93185 | CLASH | 23622248 |
| hsa-let-7b-5p | MIMAT0000063 | SDR42E1 | 93517 | PAR-CLIP | 24398324 |
| hsa-let-7b-5p | MIMAT0000063 | ZFAND4 | 93550 | HITS-CLIP//PAR-CLIP | 23706177\|23313552\|27292025 |
| hsa-let-7b-5p | MIMAT0000063 | MFSD3 | 113655 | CLASH | 23622248 |
| hsa-let-7b-5p | MIMAT0000063 | TOE1 | 114034 | Proteomics | 18668040 |
| hsa-let-7b-5p | MIMAT0000063 | BTBD9 | 114781 | CLASH | 23622248 |
| hsa-let-7b-5p | MIMAT0000063 | MBD6 | 114785 | CLASH | 23622248 |
| hsa-let-7b-5p | MIMAT0000063 | OSBPL10 | 114884 | Proteomics | 18668040 |
| hsa-let-7b-5p | MIMAT0000063 | KCTD12 | 115207 | Proteomics | 18668040 |

| | | | | | |
|---|---|---|---|---|---|
| hsa-let-7b-5p | MIMAT0000063 | CTHRC1 | 115908 | Luciferase reporter assay//qRT-PCR//Western blot | 25510669 |
| hsa-let-7b-5p | MIMAT0000063 | TSEN15 | 116461 | CLASH | 23622248 |
| hsa-let-7b-5p | MIMAT0000063 | DCD | 117159 | Proteomics | 18668040 |
| hsa-let-7b-5p | MIMAT0000063 | RFFL | 117584 | CLASH | 23622248 |
| hsa-let-7b-5p | MIMAT0000063 | PDZD8 | 118987 | PAR-CLIP | 21572407 |
| hsa-let-7b-5p | MIMAT0000063 | LRIG3 | 121227 | PAR-CLIP | 21572407\|20371350 |
| hsa-let-7b-5p | MIMAT0000063 | NAA30 | 122830 | PAR-CLIP | 21572407 |
| hsa-let-7b-5p | MIMAT0000063 | MSI2 | 124540 | PAR-CLIP | 21572407 |
| hsa-let-7b-5p | MIMAT0000063 | C19orf47 | 126526 | PAR-CLIP | 21572407\|20371350 |
| hsa-let-7b-5p | MIMAT0000063 | GABPB2 | 126626 | CLASH | 23622248 |
| hsa-let-7b-5p | MIMAT0000063 | TBC1D20 | 128637 | CLASH | 23622248 |
| hsa-let-7b-5p | MIMAT0000063 | PIGU | 128869 | Proteomics | 18668040 |
| hsa-let-7b-5p | MIMAT0000063 | NUP35 | 129401 | Proteomics | 18668040 |
| hsa-let-7b-5p | MIMAT0000063 | TRIM71 | 131405 | PAR-CLIP | 20371350 |
| hsa-let-7b-5p | MIMAT0000063 | FAM131A | 131408 | CLASH | 23622248 |
| hsa-let-7b-5p | MIMAT0000063 | FAM43A | 131583 | PAR-CLIP | 23592263\|20371350 |
| hsa-let-7b-5p | MIMAT0000063 | GRPEL2 | 134266 | CLASH//PAR-CLIP//Proteomics//pSILAC | 18668040\|23622248\|23592263\|24398324\|23446348\|21572407\|20371350\|26701625 |
| hsa-let-7b-5p | MIMAT0000063 | C5orf24 | 134553 | CLASH | 23622248 |
| hsa-let-7b-5p | MIMAT0000063 | PM20D2 | 135293 | PAR-CLIP | 21572407 |
| hsa-let-7b-5p | MIMAT0000063 | MTPN | 136319 | Reporter assay | 15806104 |
| hsa-let-7b-5p | MIMAT0000063 | UBXN2B | 137886 | PAR-CLIP | 21572407 |
| hsa-let-7b-5p | MIMAT0000063 | GPAT4 | 137964 | PAR-CLIP | 26701625 |
| hsa-let-7b-5p | MIMAT0000063 | BRI3BP | 140707 | HITS-CLIP | 23706177 |
| hsa-let-7b-5p | MIMAT0000063 | SMCR8 | 140775 | PAR-CLIP | 24398324\|23446348\|21572407\|26701625\|27292025 |
| hsa-let-7b-5p | MIMAT0000063 | E2F7 | 144455 | CLASH | 23622248 |
| hsa-let-7b-5p | MIMAT0000063 | ZNF578 | 147660 | HITS-CLIP | 23706177 |
| hsa-let-7b-5p | MIMAT0000063 | ZNF417 | 147687 | HITS-CLIP | 23313552 |
| hsa-let-7b-5p | MIMAT0000063 | ZNF738 | 148203 | HITS-CLIP | 23706177 |
| hsa-let-7b-5p | MIMAT0000063 | SLC30A7 | 148867 | Proteomics | 18668040 |
| hsa-let-7b-5p | MIMAT0000063 | C1orf210 | 149466 | HITS-CLIP//PAR-CLIP | 23446348\|21572407\|23706177 |
| hsa-let-7b-5p | MIMAT0000063 | DTX3L | 151636 | HITS-CLIP | 23706177 |
| hsa-let-7b-5p | MIMAT0000063 | CMC1 | 152100 | Proteomics | 18668040 |
| hsa-let-7b-5p | MIMAT0000063 | DAB2IP | 153090 | CLASH | 23622248 |

| | | | | | |
|---|---|---|---|---|---|
| hsa-let-7b-5p | MIMAT0000063 | CEP120 | 153241 | PAR-CLIP | 23592263 |
| hsa-let-7b-5p | MIMAT0000063 | TMEM167A | 153339 | Proteomics | 18668040 |
| hsa-let-7b-5p | MIMAT0000063 | TMEM65 | 157378 | Proteomics | 18668040 |
| hsa-let-7b-5p | MIMAT0000063 | RDH10 | 157506 | Luciferase reporter assay//Proteomics//pSILAC | 18668040 |
| hsa-let-7b-5p | MIMAT0000063 | ANKRD46 | 157567 | PAR-CLIP | 22100165\|26701625 |
| hsa-let-7b-5p | MIMAT0000063 | FAM84B | 157638 | CLASH | 23622248 |
| hsa-let-7b-5p | MIMAT0000063 | USP54 | 159195 | CLASH | 23622248 |
| hsa-let-7b-5p | MIMAT0000063 | TMTC3 | 160418 | PAR-CLIP//Proteomics | 18668040\|23592263 |
| hsa-let-7b-5p | MIMAT0000063 | IFNLR1 | 163702 | PAR-CLIP | 23592263 |
| hsa-let-7b-5p | MIMAT0000063 | AGO3 | 192669 | Microarray | 17699775 |
| hsa-let-7b-5p | MIMAT0000063 | TMEM201 | 199953 | CLASH//Proteomics | 18668040\|23622248 |
| hsa-let-7b-5p | MIMAT0000063 | TXLNA | 200081 | CLASH//PAR-CLIP | 23622248\|21572407 |
| hsa-let-7b-5p | MIMAT0000063 | IBA57 | 200205 | CLASH | 23622248 |
| hsa-let-7b-5p | MIMAT0000063 | KLHDC8B | 200942 | PAR-CLIP | 23592263 |
| hsa-let-7b-5p | MIMAT0000063 | CENPV | 201161 | Proteomics | 18668040 |
| hsa-let-7b-5p | MIMAT0000063 | UNC13D | 201294 | Proteomics | 18668040 |
| hsa-let-7b-5p | MIMAT0000063 | ZNF584 | 201514 | HITS-CLIP | 23706177 |
| hsa-let-7b-5p | MIMAT0000063 | PDE12 | 201626 | Microarray//PAR-CLIP//Proteomics | 17699775\|18668040\|24398324\|23446348\|21572407\|20371350 |
| hsa-let-7b-5p | MIMAT0000063 | TUBB | 203068 | CLASH | 23622248 |
| hsa-let-7b-5p | MIMAT0000063 | HIPK1 | 204851 | CLASH | 23622248 |
| hsa-let-7b-5p | MIMAT0000063 | CCNY | 219771 | Proteomics | 18668040 |
| hsa-let-7b-5p | MIMAT0000063 | SLC16A9 | 220963 | PAR-CLIP | 21572407 |
| hsa-let-7b-5p | MIMAT0000063 | FOXK1 | 221937 | Proteomics | 18668040 |
| hsa-let-7b-5p | MIMAT0000063 | TMED4 | 222068 | PAR-CLIP | 27292025 |
| hsa-let-7b-5p | MIMAT0000063 | SLC35F1 | 222553 | CLASH | 23622248 |
| hsa-let-7b-5p | MIMAT0000063 | AFG1L | 246269 | CLASH | 23622248 |
| hsa-let-7b-5p | MIMAT0000063 | MMS22L | 253714 | Proteomics//pSILAC | 18668040 |
| hsa-let-7b-5p | MIMAT0000063 | RNF144B | 255488 | PAR-CLIP | 23592263 |
| hsa-let-7b-5p | MIMAT0000063 | ELMOD2 | 255520 | CLASH | 23622248 |
| hsa-let-7b-5p | MIMAT0000063 | MFSD8 | 256471 | PAR-CLIP | 24398324 |
| hsa-let-7b-5p | MIMAT0000063 | PGM2L1 | 283209 | PAR-CLIP | 23592263 |
| hsa-let-7b-5p | MIMAT0000063 | KCTD21 | 283219 | PAR-CLIP | 23592263 |
| hsa-let-7b-5p | MIMAT0000063 | TTC9C | 283237 | Proteomics//pSILAC | 18668040 |
| hsa-let-7b-5p | MIMAT0000063 | ANKRD52 | 283373 | CLASH | 23622248 |
| hsa-let-7b-5p | MIMAT0000063 | SPRYD4 | 283377 | Proteomics//pSILAC | 18668040 |

| | | | | | |
|---|---|---|---|---|---|
| hsa-let-7b-5p | MIMAT0000063 | GATC | 283459 | Proteomics | 18668040 |
| hsa-let-7b-5p | MIMAT0000063 | PRTG | 283659 | CLASH | 23622248 |
| hsa-let-7b-5p | MIMAT0000063 | ZADH2 | 284273 | CLASH//Proteomics | 18668040\|23622248 |
| hsa-let-7b-5p | MIMAT0000063 | ZNF841 | 284371 | CLASH | 23622248 |
| hsa-let-7b-5p | MIMAT0000063 | NBPF15 | 284565 | CLASH | 23622248 |
| hsa-let-7b-5p | MIMAT0000063 | C5orf51 | 285636 | PAR-CLIP | 23592263\|24398324\|23446348\|21572407\|20371350\|26701625\|27292025 |
| hsa-let-7b-5p | MIMAT0000063 | RNASE10 | 338879 | CLASH | 23622248 |
| hsa-let-7b-5p | MIMAT0000063 | ZBTB8OS | 339487 | HITS-CLIP//PAR-CLIP | 23446348\|23706177 |
| hsa-let-7b-5p | MIMAT0000063 | NAT8L | 339983 | PAR-CLIP | 21572407 |
| hsa-let-7b-5p | MIMAT0000063 | ACER2 | 340485 | PAR-CLIP | 24398324 |
| hsa-let-7b-5p | MIMAT0000063 | ZNF774 | 342132 | HITS-CLIP//PAR-CLIP | 22100165\|27418678 |
| hsa-let-7b-5p | MIMAT0000063 | ATXN1L | 342371 | CLASH | 23622248 |
| hsa-let-7b-5p | MIMAT0000063 | MTX3 | 345778 | HITS-CLIP | 23706177 |
| hsa-let-7b-5p | MIMAT0000063 | SPATA12 | 353324 | CLASH | 23622248 |
| hsa-let-7b-5p | MIMAT0000063 | IRF2BP2 | 359948 | CLASH | 23622248 |
| hsa-let-7b-5p | MIMAT0000063 | NDUFA4P1 | 360165 | PAR-CLIP | 22100165 |
| hsa-let-7b-5p | MIMAT0000063 | NHLRC2 | 374354 | HITS-CLIP | 23313552 |
| hsa-let-7b-5p | MIMAT0000063 | DRAXIN | 374946 | CLASH | 23622248 |
| hsa-let-7b-5p | MIMAT0000063 | NHLRC3 | 387921 | PAR-CLIP | 24398324\|21572407\|20371350 |
| hsa-let-7b-5p | MIMAT0000063 | BEND4 | 389206 | PAR-CLIP | 21572407\|20371350 |
| hsa-let-7b-5p | MIMAT0000063 | PSMG4 | 389362 | CLASH | 23622248 |
| hsa-let-7b-5p | MIMAT0000063 | LIN28B | 389421 | Luciferase reporter assay//Microarray | 17699775\|16971064 |
| hsa-let-7b-5p | MIMAT0000063 | RBM12B | 389677 | PAR-CLIP//Proteomics | 18668040\|21572407\|20371350 |
| hsa-let-7b-5p | MIMAT0000063 | NUDT19 | 390916 | Proteomics//pSILAC | 18668040 |
| hsa-let-7b-5p | MIMAT0000063 | ZNF805 | 390980 | CLASH | 23622248 |
| hsa-let-7b-5p | MIMAT0000063 | RAB19 | 401409 | HITS-CLIP//PAR-CLIP | 23706177\|23313552\|27292025 |
| hsa-let-7b-5p | MIMAT0000063 | POTEG | 404785 | CLASH//PAR-CLIP | 23622248\|26701625\|27292025 |
| hsa-let-7b-5p | MIMAT0000063 | NOMO3 | 408050 | CLASH | 23622248 |
| hsa-let-7b-5p | MIMAT0000063 | FNDC9 | 408263 | PAR-CLIP | 26701625 |
| hsa-let-7b-5p | MIMAT0000063 | ATXN7L3B | 552889 | PAR-CLIP | 23592263\|20371350 |
| hsa-let-7b-5p | MIMAT0000063 | PRR5-ARHGAP8 | 553158 | PAR-CLIP | 27292025 |
| hsa-let-7b-5p | MIMAT0000063 | POTEM | 641455 | PAR-CLIP | 26701625\|27292025 |
| hsa-let-7b-5p | MIMAT0000063 | FAM83G | 644815 | PAR-CLIP | 23592263\|24398324 |
| hsa-let-7b-5p | MIMAT0000063 | ANXA8 | 653145 | Proteomics | 18668040 |

| | | | | | |
|---|---|---|---|---|---|
| hsa-let-7b-5p | MIMAT0000063 | ANXA8L1 | 728113 | Proteomics | 18668040 |
| hsa-let-7b-5p | MIMAT0000063 | CASTOR2 | 729438 | PAR-CLIP | 26701625 |
| hsa-let-7b-5p | MIMAT0000063 | POM121C | 100101267 | Proteomics | 18668040 |
| hsa-let-7b-5p | MIMAT0000063 | TIMM23 | 100287932 | Proteomics | 18668040 |
| hsa-let-7b-5p | MIMAT0000063 | OCLN | 100506658 | CLASH | 23622248 |
| hsa-mir-22-3p | MIMAT0000077 | ACLY | 47 | Luciferase reporter assay//qRT-PCR//Western blot | 27317765 |
| hsa-mir-22-3p | MIMAT0000077 | AKT1 | 207 | Luciferase reporter assay//Western blot | 26364720 |
| hsa-mir-22-3p | MIMAT0000077 | BDNF | 627 | Luciferase reporter assay//Microarray | 21168126 |
| hsa-mir-22-3p | MIMAT0000077 | BMP6 | 654 | Luciferase reporter assay | 24163368 |
| hsa-mir-22-3p | MIMAT0000077 | BMP7 | 655 | Luciferase reporter assay//qRT-PCR//Western blot | 19011694\|24163368 |
| hsa-mir-22-3p | MIMAT0000077 | BMPR1B | 658 | Luciferase reporter assay | 24163368 |
| hsa-mir-22-3p | MIMAT0000077 | BSG | 682 | Immunohistochemistry//Luciferase reporter assay//qRT-PCR//Western blot | 24906624\|28184176 |
| hsa-mir-22-3p | MIMAT0000077 | BTF3 | 689 | Sequencing | 20371350 |
| hsa-mir-22-3p | MIMAT0000077 | BTG1 | 694 | Luciferase reporter assay | 25449431 |
| hsa-mir-22-3p | MIMAT0000077 | BUB1B | 701 | CLASH | 23622248 |
| hsa-mir-22-3p | MIMAT0000077 | CCNA2 | 890 | Luciferase reporter assay | 25596928 |
| hsa-mir-22-3p | MIMAT0000077 | CCNT2 | 905 | PAR-CLIP//Sequencing | 20371350\|24398324 |
| hsa-mir-22-3p | MIMAT0000077 | CD151 | 977 | Luciferase reporter assay//qRT-PCR//Western blot | 24495805 |
| hsa-mir-22-3p | MIMAT0000077 | CDK6 | 1021 | Sequencing | 20371350 |
| hsa-mir-22-3p | MIMAT0000077 | CDKN1A | 1026 | Luciferase reporter assay//PAR-CLIP//qRT-PCR//Western blot | 23582783\|21572407 |
| hsa-mir-22-3p | MIMAT0000077 | CSF1R | 1436 | Luciferase reporter assay//qRT-PCR | 24198819 |
| hsa-mir-22-3p | MIMAT0000077 | CSNK2A1 | 1457 | Sequencing | 20371350 |
| hsa-mir-22-3p | MIMAT0000077 | DAD1 | 1603 | PAR-CLIP | 27292025 |
| hsa-mir-22-3p | MIMAT0000077 | DDX6 | 1656 | PAR-CLIP | 22012620 |

| | | | | | |
|---|---|---|---|---|---|
| hsa-mir-22-3p | MIMAT0000077 | E2F2 | 1870 | PAR-CLIP//Sequencing | 20371350\|26701625 |
| hsa-mir-22-3p | MIMAT0000077 | ERBB2 | 2064 | Luciferase reporter assay | 26544868 |
| hsa-mir-22-3p | MIMAT0000077 | ERBB3 | 2065 | Luciferase reporter assay//qRT-PCR//Western blot | 22484852 |
| hsa-mir-22-3p | MIMAT0000077 | ESR1 | 2099 | Immunoblot//Luciferase reporter assay//qRT-PCR//Western blot | 18347104\|19414598 |
| hsa-mir-22-3p | MIMAT0000077 | MECOM | 2122 | 5RACE//ChIP//Co-immunoprecipitation//Luciferase reporter assay//Northern blot//Western blot | 27617961 |
| hsa-mir-22-3p | MIMAT0000077 | FKBP5 | 2289 | CLASH | 23622248 |
| hsa-mir-22-3p | MIMAT0000077 | GRB2 | 2885 | PAR-CLIP | 26701625 |
| hsa-mir-22-3p | MIMAT0000077 | NR3C1 | 2908 | Sequencing | 20371350 |
| hsa-mir-22-3p | MIMAT0000077 | H3F3B | 3021 | PAR-CLIP//Sequencing | 20371350 |
| hsa-mir-22-3p | MIMAT0000077 | HIF1A | 3091 | Immunofluorescence//qRT-PCR//Western blot | 24496460 |
| hsa-mir-22-3p | MIMAT0000077 | HMGB1 | 3146 | Luciferase reporter assay//qRT-PCR//Western blot | 23303785 |
| hsa-mir-22-3p | MIMAT0000077 | HSPA1B | 3304 | CLASH | 23622248 |
| hsa-mir-22-3p | MIMAT0000077 | HTR2C | 3358 | Luciferase reporter assay//Microarray | 21168126 |
| hsa-mir-22-3p | MIMAT0000077 | CYR61 | 3491 | Luciferase reporter assay//qRT-PCR//Western blot | 24449575 |
| hsa-mir-22-3p | MIMAT0000077 | CXCR2 | 3579 | Flow//Gluc reporter assay//GUS reporter assay//HITS-CLIP//Immunoblot//Immunocytochemistry//Immunofluorescence//Immunohistochemistry//Immunoprecipitaion//Luciferase reporter assay//qRT-PCR//RNA immunoprecipitation assay (RIP)//Western blot | 26364720 |

| | | | | | |
|---|---|---|---|---|---|
| hsa-mir-22-3p | MIMAT0000077 | INSIG1 | 3638 | HITS-CLIP | 19536157 |
| hsa-mir-22-3p | MIMAT0000077 | IRF5 | 3663 | Luciferase reporter assay//qRT-PCR//Western blot | 23303785 |
| hsa-mir-22-3p | MIMAT0000077 | RPSA | 3921 | CLASH | 23622248 |
| hsa-mir-22-3p | MIMAT0000077 | LBP | 3929 | CLASH | 23622248 |
| hsa-mir-22-3p | MIMAT0000077 | LGALS1 | 3956 | Immunofluorescence//qRT-PCR//Western blot | 24496460 |
| hsa-mir-22-3p | MIMAT0000077 | LGALS9 | 3965 | Flow//Luciferase reporter assay//qRT-PCR//Western blot | 26239725 |
| hsa-mir-22-3p | MIMAT0000077 | MAOA | 4128 | Luciferase reporter assay//Microarray | 21168126 |
| hsa-mir-22-3p | MIMAT0000077 | MAX | 4149 | PAR-CLIP | 23592263 |
| hsa-mir-22-3p | MIMAT0000077 | MMP14 | 4323 | Luciferase reporter assay//Microarray//qRT-PCR//Western blot | 26610210 |
| hsa-mir-22-3p | MIMAT0000077 | MTHFR | 4524 | Immunoblot//Luciferase reporter assay//qRT-PCR//Western blot | 28045918 |
| hsa-mir-22-3p | MIMAT0000077 | MYO6 | 4646 | CLASH | 23622248 |
| hsa-mir-22-3p | MIMAT0000077 | NTRK2 | 4915 | Luciferase reporter assay//qRT-PCR//Western blot | 27662840 |
| hsa-mir-22-3p | MIMAT0000077 | PDHA1 | 5160 | Sequencing | 20371350 |
| hsa-mir-22-3p | MIMAT0000077 | PIK3C2A | 5286 | Sequencing | 20371350 |
| hsa-mir-22-3p | MIMAT0000077 | PLK1 | 5347 | Microarray//qRT-PCR//Western blot | 25970317 |
| hsa-mir-22-3p | MIMAT0000077 | PPARA | 5465 | Luciferase reporter assay//qRT-PCR//Western blot | 18347104\|19011694 |
| hsa-mir-22-3p | MIMAT0000077 | PRKACA | 5566 | Sequencing | 20371350 |
| hsa-mir-22-3p | MIMAT0000077 | PTEN | 5728 | Luciferase reporter assay | 20388916\|26544868 |
| hsa-mir-22-3p | MIMAT0000077 | PTMS | 5763 | Luciferase reporter assay//Microarray//Northern blot//qRT-PCR//Western blot | 22493679 |
| hsa-mir-22-3p | MIMAT0000077 | PEX5 | 5830 | CLASH | 23622248 |

| | | | | | |
|---|---|---|---|---|---|
| hsa-mir-22-3p | MIMAT0000077 | RAB5B | 5869 | Chromatin immunoprecipitation//FACS//Immunohistochemistry//Luciferase reporter assay//Microarray//Next Generation Sequencing (NGS)//Northern blot//PAR-CLIP//qRT-PCR//Sequencing//Western blot | 20371350\|21572407\|27569217 |
| hsa-mir-22-3p | MIMAT0000077 | RAP2B | 5912 | HITS-CLIP | 23313552 |
| hsa-mir-22-3p | MIMAT0000077 | RBL1 | 5933 | PAR-CLIP | 20371350 |
| hsa-mir-22-3p | MIMAT0000077 | RGS2 | 5997 | Luciferase reporter assay//Microarray | 21168126\|23349832 |
| hsa-mir-22-3p | MIMAT0000077 | RPL24 | 6152 | CLASH | 23622248 |
| hsa-mir-22-3p | MIMAT0000077 | RPL35A | 6165 | CLASH | 23622248 |
| hsa-mir-22-3p | MIMAT0000077 | RPS2 | 6187 | CLASH | 23622248 |
| hsa-mir-22-3p | MIMAT0000077 | RPS4X | 6191 | CLASH | 23622248 |
| hsa-mir-22-3p | MIMAT0000077 | SCD | 6319 | HITS-CLIP | 23313552 |
| hsa-mir-22-3p | MIMAT0000077 | SRSF7 | 6432 | Sequencing | 20371350 |
| hsa-mir-22-3p | MIMAT0000077 | SLC2A1 | 6513 | HITS-CLIP//In situ hybridization//Luciferase reporter assay//qRT-PCR//Western blot | 23313552\|25304371 |
| hsa-mir-22-3p | MIMAT0000077 | SNAI1 | 6615 | Luciferase reporter assay//Microarray//qRT-PCR//Western blot | 26610210 |
| hsa-mir-22-3p | MIMAT0000077 | SP1 | 6667 | Luciferase reporter assay//qRT-PCR//Reporter assay//Western blot | 23529765\|21502362\|27904693 |
| hsa-mir-22-3p | MIMAT0000077 | SRPK1 | 6732 | PAR-CLIP | 20371350 |
| hsa-mir-22-3p | MIMAT0000077 | STX4 | 6810 | CLASH | 23622248 |

| | | | | | |
|---|---|---|---|---|---|
| hsa-mir-22-3p | MIMAT0000077 | TACC1 | 6867 | Chromatin immunoprecipitation//FACS//Immunohistochemistry//Luciferase reporter assay//Microarray//Next Generation Sequencing (NGS)//Northern blot//qRT-PCR//Western blot | 27569217 |
| hsa-mir-22-3p | MIMAT0000077 | TBX3 | 6926 | Sequencing | 20371350 |
| hsa-mir-22-3p | MIMAT0000077 | TCF7 | 6932 | //Luciferase reporter assay//qRT-PCR//Western blot | 26193896 |
| hsa-mir-22-3p | MIMAT0000077 | TFRC | 7037 | flow//Luciferase reporter assay//Northern blot | 19135902 |
| hsa-mir-22-3p | MIMAT0000077 | TIAM1 | 7074 | Luciferase reporter assay//Western blot | 23440286 |
| hsa-mir-22-3p | MIMAT0000077 | TPD52L2 | 7165 | PAR-CLIP | 23592263 |
| hsa-mir-22-3p | MIMAT0000077 | VSNL1 | 7447 | HITS-CLIP//PAR-CLIP | 21572407\|20371350\|23824327\|27418678 |
| hsa-mir-22-3p | MIMAT0000077 | WNT1 | 7471 | Immunohistochemistry//Luciferase reporter assay//Microarray//qRT-PCR//Western blot | 23851184 |
| hsa-mir-22-3p | MIMAT0000077 | YWHAZ | 7534 | PAR-CLIP | 24398324\|23446348\|20371350 |
| hsa-mir-22-3p | MIMAT0000077 | ZNF217 | 7764 | Sequencing | 20371350 |
| hsa-mir-22-3p | MIMAT0000077 | ALMS1 | 7840 | CLASH | 23622248 |
| hsa-mir-22-3p | MIMAT0000077 | NUP214 | 8021 | CLASH | 23622248 |
| hsa-mir-22-3p | MIMAT0000077 | SLC7A5 | 8140 | PAR-CLIP | 26701625 |
| hsa-mir-22-3p | MIMAT0000077 | NCOA1 | 8648 | Luciferase reporter assay//qRT-PCR//Western blot | 21798241\|26244872 |
| hsa-mir-22-3p | MIMAT0000077 | TNFRSF10D | 8793 | PAR-CLIP | 22012620 |
| hsa-mir-22-3p | MIMAT0000077 | FUBP1 | 8880 | CLASH | 23622248 |
| hsa-mir-22-3p | MIMAT0000077 | MTA1 | 9112 | Immunofluorescence//Immunohistochemistry//Luciferase reporter assay//qRT-PCR | 28231399 |
| hsa-mir-22-3p | MIMAT0000077 | VAPB | 9217 | CLASH | 23622248 |

| | | | | | |
|---|---|---|---|---|---|
| hsa-mir-22-3p | MIMAT0000077 | TCEAL1 | 9338 | Immunoblot//Luciferase reporter assay//qRT-PCR | 21565979 |
| hsa-mir-22-3p | MIMAT0000077 | RBM39 | 9584 | CLASH | 23622248 |
| hsa-mir-22-3p | MIMAT0000077 | ZNF646 | 9726 | PAR-CLIP | 22100165\|22291592 |
| hsa-mir-22-3p | MIMAT0000077 | IFT140 | 9742 | PAR-CLIP | 20371350 |
| hsa-mir-22-3p | MIMAT0000077 | HDAC4 | 9759 | Luciferase reporter assay//qRT-PCR//Western blot | 20842113\|23349832 |
| hsa-mir-22-3p | MIMAT0000077 | HDAC6 | 10013 | Luciferase reporter assay//qRT-PCR//Western blot | 22375943\|28195408 |
| hsa-mir-22-3p | MIMAT0000077 | ARPC5 | 10092 | EMSA//Immunohistochemistry//Luciferase reporter assay//Microarray//qRT-PCR//Western blot | 22447776 |
| hsa-mir-22-3p | MIMAT0000077 | NET1 | 10276 | Luciferase reporter assay//qRT-PCR//Western blot | 25041463 |
| hsa-mir-22-3p | MIMAT0000077 | BTN3A3 | 10384 | PAR-CLIP | 20371350 |
| hsa-mir-22-3p | MIMAT0000077 | ZNF460 | 10794 | PAR-CLIP | 23592263 |
| hsa-mir-22-3p | MIMAT0000077 | EFR3B | 22979 | HITS-CLIP | 19536157 |
| hsa-mir-22-3p | MIMAT0000077 | RCOR1 | 23186 | Luciferase reporter assay | 23349832 |
| hsa-mir-22-3p | MIMAT0000077 | TBC1D12 | 23232 | Sequencing | 20371350 |
| hsa-mir-22-3p | MIMAT0000077 | WWC1 | 23286 | PAR-CLIP | 22012620 |
| hsa-mir-22-3p | MIMAT0000077 | FRAT2 | 23401 | PAR-CLIP//Sequencing | 20371350\|21572407 |
| hsa-mir-22-3p | MIMAT0000077 | SIRT1 | 23411 | Luciferase reporter assay//qRT-PCR//Western blot | 26912776\|26662303 |
| hsa-mir-22-3p | MIMAT0000077 | TTC33 | 23548 | HITS-CLIP | 19536157 |
| hsa-mir-22-3p | MIMAT0000077 | ELP5 | 23587 | CLASH | 23622248 |
| hsa-mir-22-3p | MIMAT0000077 | LEMD3 | 23592 | Sequencing | 20371350 |
| hsa-mir-22-3p | MIMAT0000077 | ARHGEF26 | 26084 | PAR-CLIP | 27292025 |
| hsa-mir-22-3p | MIMAT0000077 | SERBP1 | 26135 | PAR-CLIP | 23446348 |
| hsa-mir-22-3p | MIMAT0000077 | TRAF3IP1 | 26146 | HITS-CLIP | 19536157 |
| hsa-mir-22-3p | MIMAT0000077 | MYCBP | 26292 | Immunoblot//Luciferase reporter assay//qRT-PCR | 20562918 |
| hsa-mir-22-3p | MIMAT0000077 | FOXP1 | 27086 | Sequencing | 20371350 |
| hsa-mir-22-3p | MIMAT0000077 | PIGP | 51227 | HITS-CLIP | 23824327 |

| | | | | | |
|---|---|---|---|---|---|
| hsa-mir-22-3p | MIMAT0000077 | UBR5 | 51366 | qRT-PCR//Western blot | 27124677 |
| hsa-mir-22-3p | MIMAT0000077 | GINS2 | 51659 | HITS-CLIP | 23824327 |
| hsa-mir-22-3p | MIMAT0000077 | CYCS | 54205 | HITS-CLIP | 23824327 |
| hsa-mir-22-3p | MIMAT0000077 | DDIT4 | 54541 | PAR-CLIP | 23592263\|21572407 |
| hsa-mir-22-3p | MIMAT0000077 | TET2 | 54790 | Flow//Immunofluorescence//Immunohistochemistry//In situ hybridization//Luciferase reporter assay//qRT-PCR//Western blot | 23830207\|23827711 |
| hsa-mir-22-3p | MIMAT0000077 | DCAF16 | 54876 | HITS-CLIP | 23313552 |
| hsa-mir-22-3p | MIMAT0000077 | LRRC20 | 55222 | HITS-CLIP | 19536157 |
| hsa-mir-22-3p | MIMAT0000077 | LRRC1 | 55227 | Sequencing | 20371350 |
| hsa-mir-22-3p | MIMAT0000077 | MIS18BP1 | 55320 | Sequencing | 20371350 |
| hsa-mir-22-3p | MIMAT0000077 | LIN7C | 55327 | Sequencing | 20371350 |
| hsa-mir-22-3p | MIMAT0000077 | CAMK2N1 | 55450 | HITS-CLIP | 23313552 |
| hsa-mir-22-3p | MIMAT0000077 | RCC2 | 55920 | HITS-CLIP | 23313552 |
| hsa-mir-22-3p | MIMAT0000077 | ZMAT5 | 55954 | PAR-CLIP | 23592263 |
| hsa-mir-22-3p | MIMAT0000077 | RBSN | 64145 | Sequencing | 20371350 |
| hsa-mir-22-3p | MIMAT0000077 | RMND5A | 64795 | PAR-CLIP | 24398324 |
| hsa-mir-22-3p | MIMAT0000077 | PHACTR4 | 65979 | PAR-CLIP | 26701625 |
| hsa-mir-22-3p | MIMAT0000077 | EDC3 | 80153 | Sequencing | 20371350 |
| hsa-mir-22-3p | MIMAT0000077 | CTC1 | 80169 | PAR-CLIP | 23446348\|20371350 |
| hsa-mir-22-3p | MIMAT0000077 | CHD9 | 80205 | HITS-CLIP | 23313552 |
| hsa-mir-22-3p | MIMAT0000077 | SPG11 | 80208 | CLASH | 23622248 |
| hsa-mir-22-3p | MIMAT0000077 | CLPTM1L | 81037 | CLASH | 23622248 |
| hsa-mir-22-3p | MIMAT0000077 | TSC22D4 | 81628 | Sequencing | 20371350 |
| hsa-mir-22-3p | MIMAT0000077 | LONP2 | 83752 | HITS-CLIP | 23824327\|27418678 |
| hsa-mir-22-3p | MIMAT0000077 | KCTD10 | 83892 | PAR-CLIP | 23592263\|24398324\|26701625 |
| hsa-mir-22-3p | MIMAT0000077 | ARID5B | 84159 | Sequencing | 20371350 |
| hsa-mir-22-3p | MIMAT0000077 | GLIS2 | 84662 | CLASH | 23622248 |
| hsa-mir-22-3p | MIMAT0000077 | MTDH | 92140 | Luciferase reporter assay//qRT-PCR//Western blot | 25323629 |
| hsa-mir-22-3p | MIMAT0000077 | SFXN1 | 94081 | CLASH | 23622248 |
| hsa-mir-22-3p | MIMAT0000077 | VASN | 114990 | HITS-CLIP | 19536157 |
| hsa-mir-22-3p | MIMAT0000077 | KCTD12 | 115207 | CLASH | 23622248 |
| hsa-mir-22-3p | MIMAT0000077 | C15orf40 | 123207 | PAR-CLIP | 23446348\|27292025 |

| | | | | | |
|---|---|---|---|---|---|
| hsa-mir-22-3p | MIMAT0000077 | C1orf87 | 127795 | HITS-CLIP | 23824327 |
| hsa-mir-22-3p | MIMAT0000077 | ACVR1C | 130399 | Luciferase reporter assay//Reporter assay//Western blot | 21224400 |
| hsa-mir-22-3p | MIMAT0000077 | C5orf24 | 134553 | PAR-CLIP//Sequencing | 20371350\|23592263\|24398324\|23446348\|21572407 |
| hsa-mir-22-3p | MIMAT0000077 | SOGA1 | 140710 | Sequencing | 20371350 |
| hsa-mir-22-3p | MIMAT0000077 | DNHD1 | 144132 | CLASH | 23622248 |
| hsa-mir-22-3p | MIMAT0000077 | TMEM120B | 144404 | HITS-CLIP | 23706177 |
| hsa-mir-22-3p | MIMAT0000077 | PDIK1L | 149420 | PAR-CLIP | 21572407 |
| hsa-mir-22-3p | MIMAT0000077 | PPM1K | 152926 | Luciferase reporter assay//qRT-PCR//Western blot | 26592513 |
| hsa-mir-22-3p | MIMAT0000077 | PRELID2 | 153768 | PAR-CLIP | 22291592 |
| hsa-mir-22-3p | MIMAT0000077 | ZNF431 | 170959 | PAR-CLIP | 27292025 |
| hsa-mir-22-3p | MIMAT0000077 | TMEM201 | 199953 | PAR-CLIP | 26701625 |
| hsa-mir-22-3p | MIMAT0000077 | IBA57 | 200205 | PAR-CLIP | 26701625 |
| hsa-mir-22-3p | MIMAT0000077 | HNRNPA3 | 220988 | HITS-CLIP | 19536157 |
| hsa-mir-22-3p | MIMAT0000077 | FOXK1 | 221937 | PAR-CLIP | 26701625 |
| hsa-mir-22-3p | MIMAT0000077 | TMED4 | 222068 | PAR-CLIP | 24398324\|21572407\|20371350 |
| hsa-mir-22-3p | MIMAT0000077 | BRWD3 | 254065 | PAR-CLIP | 23592263 |
| hsa-mir-22-3p | MIMAT0000077 | MALAT1 | 378938 | Flow//Gluc reporter assay//GUS reporter assay//HITS-CLIP//Immunoblot//Immunocytochemistry//Immunofluorescence//Immunohistochemistry//Immunoprecipitaion//Luciferase reporter assay//qRT-PCR//RNA immunoprecipitation assay (RIP)//Western blot | 26364720 |
| hsa-mir-22-3p | MIMAT0000077 | ZNF662 | 389114 | PAR-CLIP | 20371350\|27292025 |
| hsa-mir-22-3p | MIMAT0000077 | RAB44 | 401258 | PAR-CLIP | 22012620 |
| hsa-mir-22-3p | MIMAT0000077 | TMEM178B | 100507421 | HITS-CLIP | 23824327\|27418678 |
| hsa-mir-184 | MIMAT0000454 | AKT1 | 207 | Luciferase reporter assay | 27666871 |

| | | | | | |
|---|---|---|---|---|---|
| hsa-mir-184 | MIMAT0000454 | AKT2 | 208 | Luciferase reporter assay//PAR-CLIP//qRT-PCR//Western blot | 20409325\|20371350\|27418134 |
| hsa-mir-184 | MIMAT0000454 | ARHGDIA | 396 | PAR-CLIP | 26701625 |
| hsa-mir-184 | MIMAT0000454 | BCL2 | 596 | Luciferase reporter assay | 24157866 |
| hsa-mir-184 | MIMAT0000454 | BCL2L1 | 598 | PAR-CLIP | 26701625 |
| hsa-mir-184 | MIMAT0000454 | TPP1 | 1200 | PAR-CLIP | 27292025 |
| hsa-mir-184 | MIMAT0000454 | CSF1 | 1435 | PAR-CLIP | 20371350 |
| hsa-mir-184 | MIMAT0000454 | GAS1 | 2619 | qRT-PCR//Western blot | 26805687 |
| hsa-mir-184 | MIMAT0000454 | INPPL1 | 3636 | Luciferase reporter assay//Northern blot//Western blot | 19033458\|24183204 |
| hsa-mir-184 | MIMAT0000454 | LIFR | 3977 | HITS-CLIP | 23313552 |
| hsa-mir-184 | MIMAT0000454 | MEIS3P1 | 4213 | PAR-CLIP | 20371350 |
| hsa-mir-184 | MIMAT0000454 | MYC | 4609 | Luciferase reporter assay | 24157866 |
| hsa-mir-184 | MIMAT0000454 | NFATC2 | 4773 | Luciferase reporter assay//qRT-PCR//Western blot | 19286996 |
| hsa-mir-184 | MIMAT0000454 | NFIC | 4782 | PAR-CLIP | 23592263\|20371350\|26701625\|27292025 |
| hsa-mir-184 | MIMAT0000454 | OPRD1 | 4985 | PAR-CLIP | 20371350 |
| hsa-mir-184 | MIMAT0000454 | PDGFB | 5155 | Immunoblot//Luciferase reporter assay//qRT-PCR | 27825105 |
| hsa-mir-184 | MIMAT0000454 | PKM | 5315 | Immunocytochemistry//Luciferase reporter assay//qRT-PCR//Western blot | 27431728 |
| hsa-mir-184 | MIMAT0000454 | PLAGL2 | 5326 | PAR-CLIP | 26701625 |
| hsa-mir-184 | MIMAT0000454 | PTPA | 5524 | PAR-CLIP | 26701625 |
| hsa-mir-184 | MIMAT0000454 | PRKCB | 5579 | Luciferase reporter assay | 27666871 |
| hsa-mir-184 | MIMAT0000454 | FSCN1 | 6624 | PAR-CLIP | 26701625 |
| hsa-mir-184 | MIMAT0000454 | SURF6 | 6838 | PAR-CLIP | 26701625 |
| hsa-mir-184 | MIMAT0000454 | TNFAIP2 | 7127 | Immunohistochemistry//Luciferase reporter assay//qRT-PCR//Western blot | 25888093 |
| hsa-mir-184 | MIMAT0000454 | EZR | 7430 | Luciferase reporter assay//qRT-PCR//Western blot | 25251993 |

| | | | | | |
|---|---|---|---|---|---|
| hsa-mir-184 | MIMAT0000454 | CNBP | 7555 | PAR-CLIP | 23592263\|20371350 |
| hsa-mir-184 | MIMAT0000454 | SLC7A5 | 8140 | PAR-CLIP | 23592263\|24398324\|22012620\|26701625 |
| hsa-mir-184 | MIMAT0000454 | PLPP3 | 8613 | Immunoblot//Luciferase reporter assay//qRT-PCR | 27825105 |
| hsa-mir-184 | MIMAT0000454 | TM9SF4 | 9777 | PAR-CLIP | 26701625 |
| hsa-mir-184 | MIMAT0000454 | CARM1 | 10498 | PAR-CLIP | 26701625\|27292025 |
| hsa-mir-184 | MIMAT0000454 | RAI1 | 10743 | PAR-CLIP | 23592263\|20371350 |
| hsa-mir-184 | MIMAT0000454 | PPP1R13L | 10848 | Luciferase reporter assay | 28012196 |
| hsa-mir-184 | MIMAT0000454 | PPP6R1 | 22870 | PAR-CLIP | 26701625 |
| hsa-mir-184 | MIMAT0000454 | ZFPM2 | 23414 | Immunoblot//Luciferase reporter assay//qRT-PCR | 27825105 |
| hsa-mir-184 | MIMAT0000454 | SND1 | 27044 | Immunofluorescence//Immunohistochemistry//Luciferase reporter assay//qRT-PCR//Western blot | 25216670 |
| hsa-mir-184 | MIMAT0000454 | TJP3 | 27134 | PAR-CLIP | 23592263\|20371350 |
| hsa-mir-184 | MIMAT0000454 | AGO2 | 27161 | Luciferase reporter assay//PAR-CLIP//Western blot | 24361012\|23696368\|26701625 |
| hsa-mir-184 | MIMAT0000454 | DESI1 | 27351 | PAR-CLIP | 22291592 |
| hsa-mir-184 | MIMAT0000454 | TNPO2 | 30000 | PAR-CLIP | 23592263\|24398324\|22012620\|21572407\|20371350\|26701625\|27292025 |
| hsa-mir-184 | MIMAT0000454 | TACO1 | 51204 | HITS-CLIP | 23313552 |
| hsa-mir-184 | MIMAT0000454 | GNL3L | 54552 | HITS-CLIP | 19536157 |
| hsa-mir-184 | MIMAT0000454 | SELENOS | 55829 | HITS-CLIP//PAR-CLIP | 22012620\|20371350\|23313552 |
| hsa-mir-184 | MIMAT0000454 | BIN3 | 55909 | Luciferase reporter assay | 27666871 |
| hsa-mir-184 | MIMAT0000454 | LRRC8A | 56262 | PAR-CLIP | 20371350 |
| hsa-mir-184 | MIMAT0000454 | CBX8 | 57332 | PAR-CLIP | 26701625 |
| hsa-mir-184 | MIMAT0000454 | FN3K | 64122 | PAR-CLIP | 23592263 |
| hsa-mir-184 | MIMAT0000454 | KLC2 | 64837 | PAR-CLIP | 26701625 |
| hsa-mir-184 | MIMAT0000454 | SOX7 | 83595 | Flow//Luciferase reporter assay//qRT-PCR//Western blot | 24558429 |
| hsa-mir-184 | MIMAT0000454 | GPRIN1 | 114787 | PAR-CLIP | 23592263 |
| hsa-mir-184 | MIMAT0000454 | TNFRSF13C | 115650 | HITS-CLIP | 23313552 |
| hsa-mir-184 | MIMAT0000454 | USH1G | 124590 | PAR-CLIP | 24398324\|20371350 |

| | | | | | |
|---|---|---|---|---|---|
| hsa-mir-184 | MIMAT0000454 | IFFO2 | 126917 | PAR-CLIP | 26701625 |
| hsa-mir-184 | MIMAT0000454 | ZSCAN25 | 221785 | PAR-CLIP | 26701625 |
| hsa-mir-184 | MIMAT0000454 | CEP170B | 283638 | PAR-CLIP | 26701625 |
| hsa-mir-184 | MIMAT0000454 | PEAK3 | 374872 | PAR-CLIP | 27292025 |
| hsa-mir-184 | MIMAT0000454 | POM121 C | 100101 267 | PAR-CLIP | 23592263 |

**Supplementary table 2. List of miRNA protein targets from MIRNET analyses**
Experimental methods used to identify protein targets and the PMID of the related reports are indicated in the last two columns.
