## Supplemental Table 4 for "Combining Plasma Extracellular Vesicle Let-7b-5p, miR-184 and Circulating miR-22-3p Levels for NSCLC Diagnosis and for Predicting Drug Resistance"

| | Dependent variable | Predictor | Predictor | Predictor | Predicted outcome | Predicted outcome | Agreement | Model parameter | | | |
|---|---|---|---|---|---|---|---|---|---|---|---|
| Model # | outcome | hsa.let.7b.5p | hsa.miR.184 | hsa.miR.22.3p | loocv - classification output | loocv - classification binary | Agreement: outcome vs LOOCV | intercept | hsa.let.7b.5p_coeff | hsa.miR.184_coeff | hsa.miR.22.3p_coeff |
| 1 | 0 | 127305 | 5319 | 758 | 0.334 | 0 | 1 | 0.313 | -9.98E-07 | 1.16E-05 | 1.15E-04 |
| 2 | 0 | 82749 | 1911 | 546 | 0.333 | 0 | 1 | 0.380 | -1.54E-06 | 1.07E-05 | 1.08E-04 |
| 3 | 0 | 25157 | 4717 | 3137 | 0.805 | 1 | 0 | 0.502 | -2.60E-06 | 9.04E-06 | 1.04E-04 |
| 4 | 0 | 107626 | 4283 | 2114 | 0.503 | 1 | 0 | 0.310 | -9.64E-07 | 1.12E-05 | 1.18E-04 |
| 5 | 0 | 131115 | 2986 | 3283 | 0.656 | 1 | 0 | 0.209 | -1.49E-07 | 1.17E-05 | 1.32E-04 |
| 6 | 0 | 141532 | 3180 | 1419 | 0.385 | 0 | 1 | 0.282 | -7.27E-07 | 1.16E-05 | 1.19E-04 |
| 7 | 0 | 108456 | 3342 | 924 | 0.350 | 0 | 1 | 0.338 | -1.20E-06 | 1.11E-05 | 1.13E-04 |
| 8 | 0 | 129569 | 8772 | 778 | 0.380 | 0 | 1 | 0.301 | -8.87E-07 | 1.19E-05 | 1.15E-04 |
| 9 | 0 | 85214 | 5498 | 407 | 0.353 | 0 | 1 | 0.376 | -1.50E-06 | 1.10E-05 | 1.08E-04 |
| 10 | 0 | 80505 | 7630 | 874 | 0.436 | 0 | 1 | 0.380 | -1.52E-06 | 1.11E-05 | 1.08E-04 |
| 11 | 0 | 96798 | 3448 | 318 | 0.298 | 0 | 1 | 0.359 | -1.38E-06 | 1.10E-05 | 1.10E-04 |
| 12 | 1 | 43127 | 8764 | 3132 | 0.713 | 1 | 1 | 0.274 | -9.77E-07 | 1.20E-05 | 1.20E-04 |
| 13 | 1 | 82362 | 15669 | 2239 | 0.640 | 1 | 1 | 0.312 | -1.33E-06 | 1.10E-05 | 1.19E-04 |
| 14 | 1 | 33742 | 9202 | 4866 | 0.946 | 1 | 1 | 0.311 | -1.22E-06 | 1.17E-05 | 1.17E-04 |
| 15 | 1 | 27027 | 20885 | 4365 | 1.038 | 1 | 1 | 0.316 | -1.28E-06 | 1.17E-05 | 1.17E-04 |
| 16 | 1 | 35250 | 24119 | 4699 | 1.112 | 1 | 1 | 0.311 | -1.27E-06 | 1.19E-05 | 1.19E-04 |
| 17 | 1 | 35957 | 12890 | 4614 | 0.958 | 1 | 1 | 0.312 | -1.23E-06 | 1.16E-05 | 1.17E-04 |
| 18 | 1 | 27112 | 23983 | 2843 | 0.879 | 1 | 1 | 0.299 | -1.13E-06 | 1.13E-05 | 1.19E-04 |
| 19 | 1 | 82557 | 5504 | 980 | 0.342 | 0 | 0 | 0.216 | -8.74E-07 | 1.26E-05 | 1.31E-04 |
| 20 | 1 | 52805 | 23762 | 3691 | 0.953 | 1 | 1 | 0.315 | -1.26E-06 | 1.14E-05 | 1.17E-04 |
| 21 | 1 | 47405 | 998 | 9406 | 1.628 | 1 | 1 | 0.176 | -5.65E-07 | 1.07E-05 | 1.56E-04 |
| 22 | 1 | 25157 | 19415 | 6642 | 1.337 | 1 | 1 | 0.287 | -1.23E-06 | 1.23E-05 | 1.27E-04 |
| 23 | 1 | 22758 | 15395 | 4554 | 0.999 | 1 | 1 | 0.314 | -1.25E-06 | 1.16E-05 | 1.17E-04 |
| 24 | 1 | 45776 | 5191 | 4232 | 0.802 | 1 | 1 | 0.297 | -1.13E-06 | 1.20E-05 | 1.17E-04 |
| 25 | 1 | 82298 | 4274 | 3017 | 0.600 | 1 | 1 | 0.302 | -1.29E-06 | 1.21E-05 | 1.17E-04 |
| 26 | 1 | 62430 | 1839 | 2647 | 0.539 | 1 | 1 | 0.248 | -9.01E-07 | 1.29E-05 | 1.22E-04 |
| 27 | 1 | 116371 | 4893 | 1448 | 0.354 | 0 | 0 | 0.316 | -1.67E-06 | 1.19E-05 | 1.20E-04 |
| 28 | 1 | 32350 | 9734 | 3374 | 0.765 | 1 | 1 | 0.277 | -9.75E-07 | 1.19E-05 | 1.20E-04 |
| 29 | 1 | 111065 | 4519 | 5398 | 0.828 | 1 | 1 | 0.353 | -1.57E-06 | 1.15E-05 | 1.11E-04 |
| 30 | 1 | 114855 | 3444 | 1934 | 0.400 | 0 | 0 | 0.325 | -1.68E-06 | 1.20E-05 | 1.17E-04 |
| 31 | 1 | 102374 | 6342 | 2819 | 0.569 | 1 | 1 | 0.334 | -1.57E-06 | 1.16E-05 | 1.14E-04 |
| 32 | 1 | 148442 | 330 | 2661 | 0.332 | 0 | 0 | 0.428 | -2.54E-06 | 1.17E-05 | 1.04E-04 |
| 33 | 1 | 35230 | 16260 | 4413 | 0.975 | 1 | 1 | 0.313 | -1.24E-06 | 1.16E-05 | 1.17E-04 |
| 34 | 1 | 80649 | 59723 | 1039 | 1.097 | 1 | 1 | 0.296 | -1.13E-06 | 1.29E-05 | 1.17E-04 |
| 35 | 1 | 20148 | 5302 | 3884 | 0.781 | 1 | 1 | 0.270 | -9.13E-07 | 1.22E-05 | 1.19E-04 |
| | | | | | | | **80.00%** | | | | |

Result

**R-code:**

```
library(boot)
library(ROCR)
library(LUR)
data <- read.delim(file.choose("211008-P71-Input-data.txt"))
#check the data
summary(data)
str(data)
typeof(data)
hist(data$hsa.let.7b.5p)
hist(data$hsa.miR.184)
hist(data$hsa.miR.22.3p)
##LOOCV using the LUR package
loocv.data <-
loocv(data,dependent="outcome",c("hsa.let.7b.5p","hsa.miR.184","hsa.miR.22.3p"),export_coefficients=TRUE)
write.table(loocv.data,"211012-P71-LOOCV-output.txt",sep=" ")
```

**Supplemental table 4. Leave-one-out cross validation analysis**

The three predictors three predictor microRNAs (*let-7b-5p, miR-184, miR-22-3p)* were examined for the binary outcome cancer-free versus confirmed cases. 35 regression models were generated, and each time one sample was left out to build the model. The sample that was left out was classified using the model generated from the remaining 34 samples. We then compared in how many instances the prediction agreed with the actual outcome and observed an 80% match between the prediction and the outcome. The related R-code is shown below the table
