## Supplementary information 1 for "Combining Plasma Extracellular Vesicle Let-7b-5p, miR-184 and Circulating miR-22-3p Levels for NSCLC Diagnosis and for Predicting Drug Resistance"

#### Supplementary information 1. Bioinformatics pipeline for survival analysis

#code adapted from: #https://www.jianshu.com/p/5e4285289137 #https://bioc.ism.ac.jp/packages/3.2/bioc/vignettes/TCGAbiolinks/inst/doc/tcgaBiolinks.html #https://www.rdocumentation.org/packages/survminer/versions/0.1.1/topics/ggsurvplot

#hsa-mir-184

```
samplesTP <- TCGAquery_SampleTypes(barcode = colnames(miR_matrix),typesample = c("TP"))
mir184_exp <- miR_matrix[c("hsa-mir-184"),samplesTP]
names(mir184_exp) <-  sapply(strsplit(names(mir184_exp),'-'),function(x) paste(x[1:3],collapse="-"))
clinical$GENE <- mir184_exp[clinical$submitter_id]

df1<-subset(clinical,select=c(submitter_id,vital_status,days_to_death,days_to_last_follow_up,GENE))
df1$years_death<- df1$days_to_death/365#convert days to years
df1$years_to_last_follow_up <- df1$days_to_last_follow_up/365
df1$os<-ifelse(df1$vital_status=='Alive',df1$years_to_last_follow_up,df1$years_death)#alive sample uses
df1 <- df1[!is.na(df1$GENE),]#Remove samples with 0 expression
df1[df1$vital_status=='Dead',]$vital_status <- 2
df1[df1$vital_status=='Alive',]$vital_status <- 1
df1$vital_status <- as.numeric(df1$vital_status)

#Determine the optimal cutpoint of variables
df1.cut <- surv_cutpoint(df1,
  time = "os",
  event = "vital_status",
  variables = c("GENE"))

summary(df1.cut)
```

```
##      cutpoint statistic
## GENE 13.91485  2.567961
```

```
#Categorize variables
df1.cat<-surv_categorize(df1.cut,variables = NULL, labels = c("low", "high"))
head(df1.cat)
```

```
##          os vital_status GENE
## 1 2.5945205            1  low
## 2 2.9616438            2  low
## 3 0.7342466            2  low
## 4 0.8410959            2  low
## 5 0.0000000            2  low
## 6 1.4931507            1  low
```

```
df1.cat
```

```
##              os vital_status GENE
## 1    2.59452055            1  low
## 2    2.96164384            2  low
## 3    0.73424658            2  low
## 4    0.84109589            2  low
## 5    0.00000000            2  low
## 6    1.49315068            1  low
## 7    1.47123288            1  low
## 8    0.06027397            2  low
## 9    8.14520548            1  low
## 10           NA            2  low
## 11   0.63013699            1  low
## 12   3.41369863            1  low
## 13   5.77808219            1  low
## 14   1.40547945            1  low
## 15   2.56986301            1  low
## 16   2.00000000            1  low
## 17   5.10684932            1 high
## 18   0.38082192            1  low
## 19   0.78630137            1  low
## 20   4.19452055            2  low
## 21   1.66301370            2 high
## 22   1.63835616            2  low
## 23   2.21369863            2  low
## 24   0.61643836            1  low
## 25   3.14520548            1  low
## 26   5.92054795            1  low
## 27   0.76986301            2  low
## 28   4.44109589            1  low
## 29   1.62465753            2  low
## 30   2.16712329            1  low
## 31   1.24657534            1  low
## 32   1.79726027            2  low
## 33   7.33150685            1  low
## 34   0.96712329            1  low
## 35   0.38630137            1  low
## 36   1.96986301            2  low
## 37   4.79452055            1  low
## 38   1.66301370            1  low
## 39   3.00547945            1  low
## 40   1.33424658            1  low
## 41   0.36712329            1  low
## 42   2.43287671            1  low
## 43   0.01095890            2  low
## 44   7.09589041            1  low
## 45   4.92602740            2  low
## 46   0.13150685            1  low
## 47   1.19178082            1 high
## 48   2.38082192            2  low
## 49   4.87123288            2  low
## 50   3.95068493            1  low
## 51   3.39452055            1  low
```

```
## 52   1.80000000            1  low
## 53  10.79452055            1  low
## 54   2.45479452            1  low
## 55   5.18630137            1  low
## 56   1.12054795            1  low
## 57   0.70410959            2  low
## 58   1.22739726            1  low
## 59   0.96164384            1  low
## 60   2.08219178            2  low
## 61   0.84383562            2  low
## 62   0.16986301            1  low
## 63   7.10958904            1  low
## 64   4.11232877            2  low
## 65   2.37260274            1  low
## 66   1.19178082            1  low
## 67   6.46575342            1  low
## 68   1.69041096            1  low
## 69   2.03013699            1  low
## 70   1.89041096            1  low
## 71   2.43561644            1  low
## 72   1.61917808            1  low
## 73   1.22739726            1  low
## 74   4.03835616            1  low
## 75   1.78356164            1  low
## 77   0.04109589            1  low
## 78   0.96986301            2  low
## 79   3.16986301            1  low
## 80   2.54794521            1  low
## 81   1.28219178            2  low
## 82   3.10958904            2  low
## 83   0.09041096            2  low
## 84   1.30958904            2  low
## 85   3.98356164            2  low
## 86   1.99452055            1  low
## 87   1.13424658            2  low
## 88   0.57534247            2  low
## 89   0.15890411            2  low
## 90   2.33424658            1 high
## 91   2.72054795            1  low
## 92   1.17534247            2  low
## 93   2.70684932            1  low
## 94   1.78356164            1  low
## 95   5.85479452            1  low
## 96   3.27945205            2  low
## 97   1.30684932            1  low
## 98   0.50410959            1  low
## 99   0.92054795            2  low
## 100  1.25205479            1  low
## 101  8.38082192            1  low
## 102  1.85479452            2  low
## 103  1.13424658            1  low
## 104  1.58356164            1 high
## 105  2.67397260            2  low
## 106  2.54520548            2  low
```

```
## 107  0.44109589            2  low
## 108  1.28493151            2  low
## 109  1.03013699            2  low
## 110  2.02739726            1  low
## 111  1.83561644            1  low
## 112  2.50136986            1 high
## 113  0.45205479            1  low
## 114  1.37808219            2  low
## 115  1.56986301            1 high
## 116  0.51232877            2  low
## 117  2.72602740            2  low
## 118  1.58356164            1  low
## 119  2.73150685            1  low
## 120  2.26301370            2  low
## 121  1.47671233            1 high
## 122  1.13698630            1 high
## 123  2.36712329            2  low
## 124  1.98082192            1  low
## 125  0.92876712            2  low
## 126  2.20547945            1 high
## 127  0.49041096            2  low
## 128  1.49863014            1  low
## 129  1.16712329            1  low
## 130  4.15342466            2 high
## 131  0.23835616            2  low
## 132  1.01369863            2  low
## 133  0.31780822            2  low
## 134  3.91506849            1  low
## 135  2.95616438            1 high
## 136  2.93424658            1  low
## 137  1.25205479            2  low
## 138  1.65753425            1  low
## 139  1.90136986            2  low
## 140  1.55616438            1  low
## 141  2.26575342            1  low
## 142  0.05205479            2  low
## 143  0.00000000            1  low
## 144  7.75890411            1  low
## 145  2.34246575            2  low
## 146  2.72602740            2  low
## 147  1.55616438            1  low
## 148  0.50958904            1  low
## 149  2.36438356            1  low
## 150  3.18630137            1  low
## 151  1.78904110            1  low
## 152  1.92054795            1 high
## 153  3.57534247            1  low
## 154  3.54246575            2  low
## 155          NA            1  low
## 156  2.47671233            1  low
## 157  3.08493151            1  low
## 158  0.49041096            1  low
## 159  0.66849315            2  low
## 160  1.96712329            1  low
```

```
## 161  1.51232877            1  low
## 162  1.78630137            1  low
## 163  0.06575342            1  low
## 164  8.68219178            2 high
## 165  0.32602740            1  low
## 166 19.34794521            1  low
## 167  5.29315068            1 high
## 168  1.67123288            1 high
## 169  4.65753425            1  low
## 170  6.70958904            1  low
## 171  1.64383562            1  low
## 172  3.38356164            2  low
## 173  3.77808219            2  low
## 174  0.70136986            1  low
## 175  1.16712329            1  low
## 176  4.43013699            1  low
## 177  1.20547945            1  low
## 178  2.01917808            2  low
## 179  2.25753425            1  low
## 180  1.87123288            1  low
## 181  1.63287671            1  low
## 182  0.04931507            2  low
## 183  1.26027397            2  low
## 184  7.16986301            2  low
## 185  0.83013699            2  low
## 186  3.37808219            1  low
## 188  0.09863014            1  low
## 189  2.93698630            1  low
## 190  1.33698630            2  low
## 191  2.60273973            2  low
## 192  3.08219178            1  low
## 193  1.71506849            1 high
## 194  1.92876712            1 high
## 195  2.08493151            1  low
## 196  1.63013699            1  low
## 197  5.40821918            1  low
## 198  2.40821918            2  low
## 199  0.70684932            2  low
## 200  0.68493151            2  low
## 201  4.38356164            2  low
## 202  1.67123288            1  low
## 203  1.01917808            1  low
## 204  1.65205479            1 high
## 205  1.22191781            1  low
## 206  0.21643836            1  low
## 207  1.41095890            1  low
## 208  9.95890411            1  low
## 210  1.82465753            2  low
## 211  0.03013699            1  low
## 212  1.96986301            1  low
## 213  3.25753425            1  low
## 214  0.51780822            2  low
## 215  0.13698630            1  low
## 216  0.45753425            2  low
```

```
## 217  2.31506849           1  low
## 218  7.73424658           1  low
## 220  2.58630137           1  low
## 221  5.65753425           1  low
## 222  4.10410959           2  low
## 223  0.41369863           1  low
## 224  1.16986301           1  low
## 225  2.70410959           2  low
## 226  1.47945205           1  low
## 227  0.42191781           2  low
## 228  1.80273973           1  low
## 229  0.33972603           2  low
## 230  0.84109589           1  low
## 231  0.66849315           2 high
## 232  0.87945205           2  low
## 233  2.60821918           2  low
## 234  1.55616438           1  low
## 235  2.29863014           1 high
## 236  3.75068493           1  low
## 237  7.34520548           2  low
## 238  1.15616438           1 high
## 239  1.36986301           2  low
## 240  6.81643836           1  low
## 241 18.44383562           1  low
## 242  4.44383562           2  low
## 243  0.26575342           2  low
## 244  1.44109589           1  low
## 245  3.22739726           1  low
## 246          NA           1  low
## 248  3.31232877           2  low
## 249  3.46575342           2 high
## 250  1.30958904           1  low
## 251  6.48767123           1  low
## 252  1.45479452           1  low
## 253  1.30410959           1  low
## 254  4.08767123           2  low
## 255  1.27671233           1  low
## 256  7.16712329           1  low
## 257  1.18904110           2  low
## 258          NA           2  low
## 259  3.33150685           1  low
## 260  3.83561644           1 high
## 261  1.64109589           1  low
## 262  6.55616438           2  low
## 263  1.92876712           1 high
## 264  8.93424658           1  low
## 265  7.38630137           1  low
## 266  9.05479452           1 high
## 267  1.90958904           2  low
## 268  2.84931507           1  low
## 269  0.33150685           2  low
## 270  2.81095890           2  low
## 271  3.09589041           1  low
## 272  1.21095890           2 high
```

```
## 273  3.20821918           2  low
## 274  0.79726027           2  low
## 275  3.36712329           2  low
## 276  1.62191781           1  low
## 277  0.23013699           1  low
## 278  2.60000000           1  low
## 279  1.71506849           2  low
## 280  1.13972603           1  low
## 281  0.50958904           1  low
## 282  3.19726027           2 high
## 284  3.06301370           1  low
## 285  2.16712329           1 high
## 286  0.59726027           1  low
## 287  3.11232877           2  low
## 288  1.80273973           1 high
## 289  2.16712329           1  low
## 290  1.59452055           2  low
## 291  5.06027397           1 high
## 292  0.48219178           2  low
## 293  1.32602740           1  low
## 294  3.47397260           2  low
## 295  6.15890411           1  low
## 296  2.85753425           2  low
## 297  2.41643836           1  low
## 298  0.93150685           2  low
## 299  3.27123288           2  low
## 300  0.88493151           1  low
## 301  2.60000000           2  low
## 302  2.47945205           2  low
## 303  1.64657534           1 high
## 304  1.00000000           1  low
## 306  1.42465753           1  low
## 307  2.36164384           1  low
## 308  4.10684932           2  low
## 309  0.93972603           2  low
## 310  1.72602740           1  low
## 311  6.02465753           1  low
## 312  1.30410959           1  low
## 313  0.77260274           2  low
## 314  1.11780822           1  low
## 315  0.38082192           2  low
## 316  1.13698630           1  low
## 317  2.25753425           1  low
## 318          NA           1  low
## 319  2.45479452           2  low
## 320  2.19178082           2 high
## 321  0.71232877           1  low
## 322  2.49863014           1  low
## 323  3.50684932           1  low
## 324  1.70958904           1 high
## 325  3.44657534           2  low
## 326  0.72328767           1  low
## 327          NA           1  low
## 328  1.31780822           1  low
```

```
## 329  1.62739726            2  low
## 330  0.16986301            2  low
## 331  3.48493151            1  low
## 332  0.27123288            2  low
## 333  2.45479452            2  low
## 334  0.07671233            1  low
## 335  4.18630137            2  low
## 336  0.36438356            1  low
## 337  2.93972603            2  low
## 338  2.00273973            2  low
## 339  3.17534247            1 high
## 340  2.52602740            2  low
## 341  1.68219178            1  low
## 342  1.98356164            1  low
## 343  4.17260274            1  low
## 344  1.02739726            2  low
## 345  4.27123288            1  low
## 346  1.41095890            1  low
## 347  0.82191781            2  low
## 348  1.72054795            2  low
## 349  1.52602740            2  low
## 350  1.65205479            1  low
## 351  1.57260274            2  low
## 352  3.74520548            1  low
## 353  2.16712329            1  low
## 354  0.66575342            2  low
## 355  4.47123288            2  low
## 356  2.90410959            1  low
## 357  0.87945205            2 high
## 358  5.55342466            2  low
## 359  1.34794521            1  low
## 360  1.65753425            1  low
## 361  1.20547945            2  low
## 362  2.21095890            2  low
## 363  1.12602740            1  low
## 364  4.90410959            2  low
## 365  1.93150685            1  low
## 366  0.96986301            1  low
## 367  5.66301370            1  low
## 368 10.29863014            1  low
## 369  2.20821918            1  low
## 370  1.30410959            1  low
## 371  2.43561644            1  low
## 372  1.12054795            2  low
## 373  1.85479452            1  low
## 374  2.30684932            1  low
## 375  9.20821918            2  low
## 376  0.24931507            2  low
## 377  1.18904110            2  low
## 378  2.02465753            1  low
## 379  1.26575342            1  low
## 380  0.03835616            1  low
## 381  2.11780822            1  low
## 382  3.70136986            1 high
```

```
## 383  1.92328767            2  low
## 384  0.55342466            1  low
## 385  1.53698630            2  low
## 386  2.22739726            1 high
## 387  1.49589041            1  low
## 388  1.70958904            2  low
## 389  1.21643836            2  low
## 390  1.89315068            1  low
## 391  1.50958904            1  low
## 392  2.41643836            1 high
## 393  4.61095890            1  low
## 394  5.01369863            2  low
## 395  1.78904110            2  low
## 396  3.92328767            1  low
## 397  8.47671233            1  low
## 398  1.78630137            1  low
## 399  2.08493151            2  low
## 400  1.66849315            1  low
## 401  3.89315068            2  low
## 402  1.65205479            1  low
## 403  1.54794521            1  low
## 404  1.51506849            1  low
## 405  0.78082192            1  low
## 406  3.06575342            1  low
## 407  3.71780822            2  low
## 408  1.84657534            1  low
## 409  0.47397260            2  low
## 410  0.75068493            2  low
## 411  2.12328767            1  low
## 412  0.84931507            1  low
## 413  2.36712329            1  low
## 414  0.92328767            1  low
## 415  0.46849315            2  low
## 416  0.75342466            2  low
## 417  0.61369863            1  low
## 418  0.12054795            1  low
## 419  1.43013699            1  low
## 420  3.05479452            2  low
## 421  1.55342466            1  low
## 423  1.05479452            1  low
## 424  6.19452055            1  low
## 425  0.23013699            1  low
## 426  0.20273973            2  low
## 427  0.44931507            2  low
## 428  0.92054795            2  low
## 429  1.71232877            2 high
## 430  3.53150685            1 high
## 431 13.67671233            1 high
## 432  1.83287671            1  low
## 433  1.16164384            1  low
## 434  3.65205479            1  low
## 435  1.66575342            1 high
## 436  1.73698630            1  low
## 437  2.49315068            1  low
```

```
## 438  1.05479452            1  low
## 439  1.21095890            1 high
## 440  0.03561644            1  low
## 441  2.04657534            1 high
## 442  3.08493151            1  low
## 443  6.89041096            1  low
## 444  1.60547945            2 high
## 445  0.64931507            2  low
## 446  0.12054795            1  low
## 447  5.95616438            2  low
## 448  1.33150685            1  low
## 449  1.50684932            2  low
## 450  0.35342466            1  low
## 451  1.03287671            1  low
## 452  1.83561644            1  low
## 453  4.73424658            1  low
## 454          NA            2  low
## 455  1.15890411            1 high
## 456  3.32876712            2  low
## 457  1.88767123            1  low
## 458  1.69041096            1  low
## 459  1.58356164            1 high
## 460  1.46301370            1  low
## 461 13.59178082            2  low
## 462  1.94794521            2  low
## 463  1.05479452            2  low
## 464  0.52876712            2  low
## 465 10.06575342            1 high
## 466  3.25753425            1  low
## 467  0.16438356            1  low
## 468  1.81917808            1  low
## 469  1.19178082            1  low
## 470  4.52876712            2  low
## 471  5.12328767            1  low
## 472  4.05205479            1  low
## 473  7.17808219            2  low
## 474  3.21917808            1  low
## 475  5.38356164            1  low
## 476  3.92054795            1  low
## 477  0.09589041            1  low
## 478  1.17260274            2  low
## 479 13.05479452            1  low
## 480  0.71232877            2  low
## 481 19.85753425            1  low
## 482  2.08493151            1  low
## 483  1.93150685            1 high
## 484  2.67671233            1  low
## 485  2.27123288            1  low
## 486  3.52876712            2  low
## 487  2.12876712            2  low
## 488  1.28219178            2  low
## 490  1.38356164            1 high
## 491  2.83835616            1  low
## 492  3.62739726            1  low
```

```
## 493  3.52054795           1  low
## 494  1.92054795           2  low
## 495  1.14520548           1  low
## 496  2.77534247           1 high
## 497          NA           2  low
## 498  6.09315068           1  low
## 499  1.46849315           1  low
## 500  1.36712329           1  low
## 501  6.35068493           2  low
## 502  2.38904110           1  low
## 503  1.72328767           1  low
## 504  1.21643836           2  low
## 505  0.47671233           1  low
## 506  0.49863014           1  low
## 507  4.72602740           2  low
## 508  1.27123288           2  low
## 509  0.00000000           1  low
## 510  2.27397260           1  low
## 511  3.14246575           2  low
## 512  0.32602740           2  low
## 513  1.67123288           1  low
## 514  2.73698630           2  low
## 515  1.71506849           1  low
## 516  2.86575342           2  low
## 517  1.54520548           1  low
## 518  1.14246575           1  low
## 519  0.00000000           1  low
## 520          NA           1 high
## 521  3.56438356           1  low
## 522  1.48219178           1  low
```

```
#Determine the level of miRNA expression
fit <- survfit(Surv(os, vital_status)~GENE, data=df1.cat) #Modeling based on expression
ggsurvplot(fit,data = df1.cat,title    = "hsa-mir-184", size = 1 ,xlim = c(0, 5) ,ylim=c(1,100), risk.ta
   font.subtitle = c(20, "bold.plain", "black"),
   font.caption = c(25, "plain", "black"),
   font.x = c(25, "plain", "black"),
   font.y = c(25, "plain", "black"),
   font.tickslab = c(24, "plain", "black"), fun = "pct", break.x.by =1)
```

```
## Warning: Vectorized input to `element_text()` is not officially supported.
## Results may be unexpected or may change in future versions of ggplot2.
```

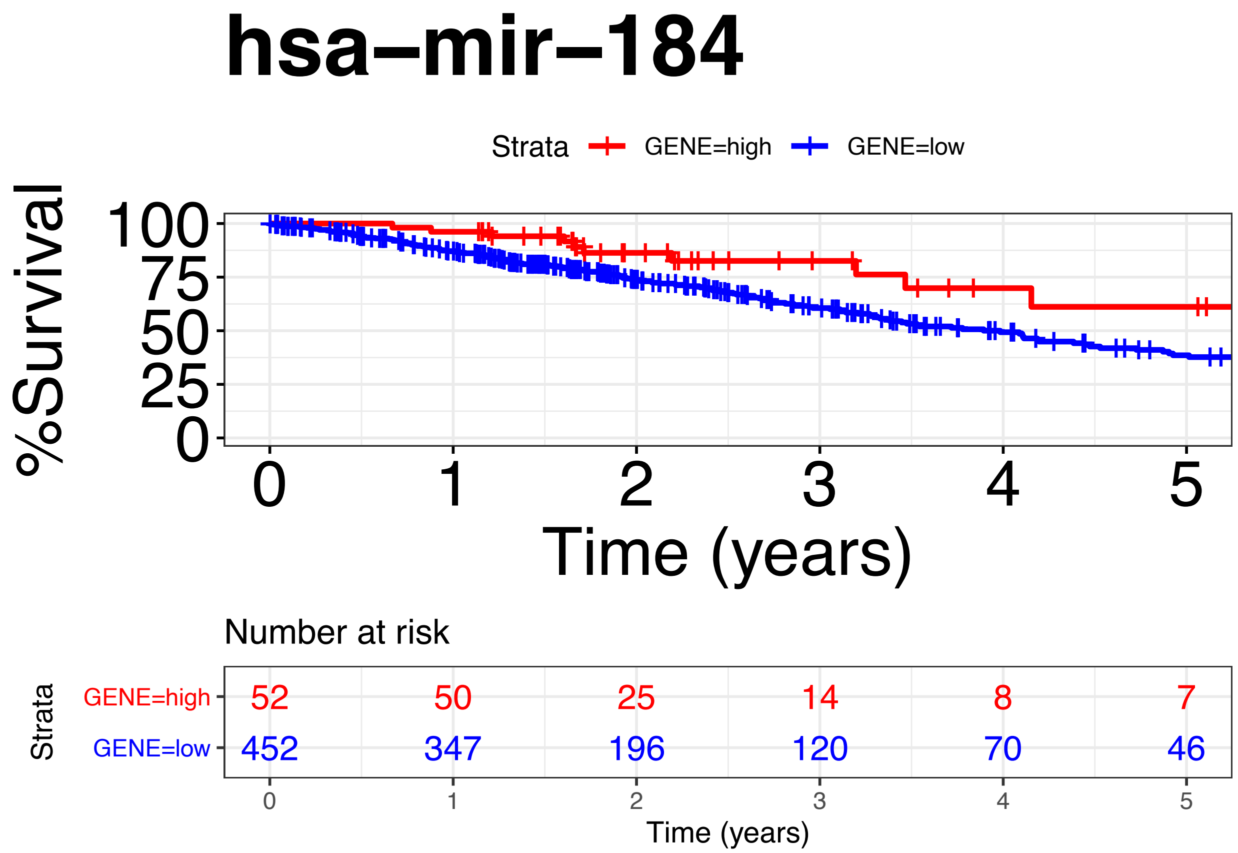


```
ggsurvplot(fit,data = df1.cat,title    = "hsa-mir-184", size = 1,xlim = c(0, 5),ylim=c(1,100),xlab = "Ti
   font.subtitle = c(20, "bold.plain", "black"),
   font.caption = c(25, "plain", "black"),
   font.x = c(25, "plain", "black"),
   font.y = c(25, "plain", "black"),
   font.tickslab = c(24, "plain", "black"), fun = "pct", break.x.by =1)
```

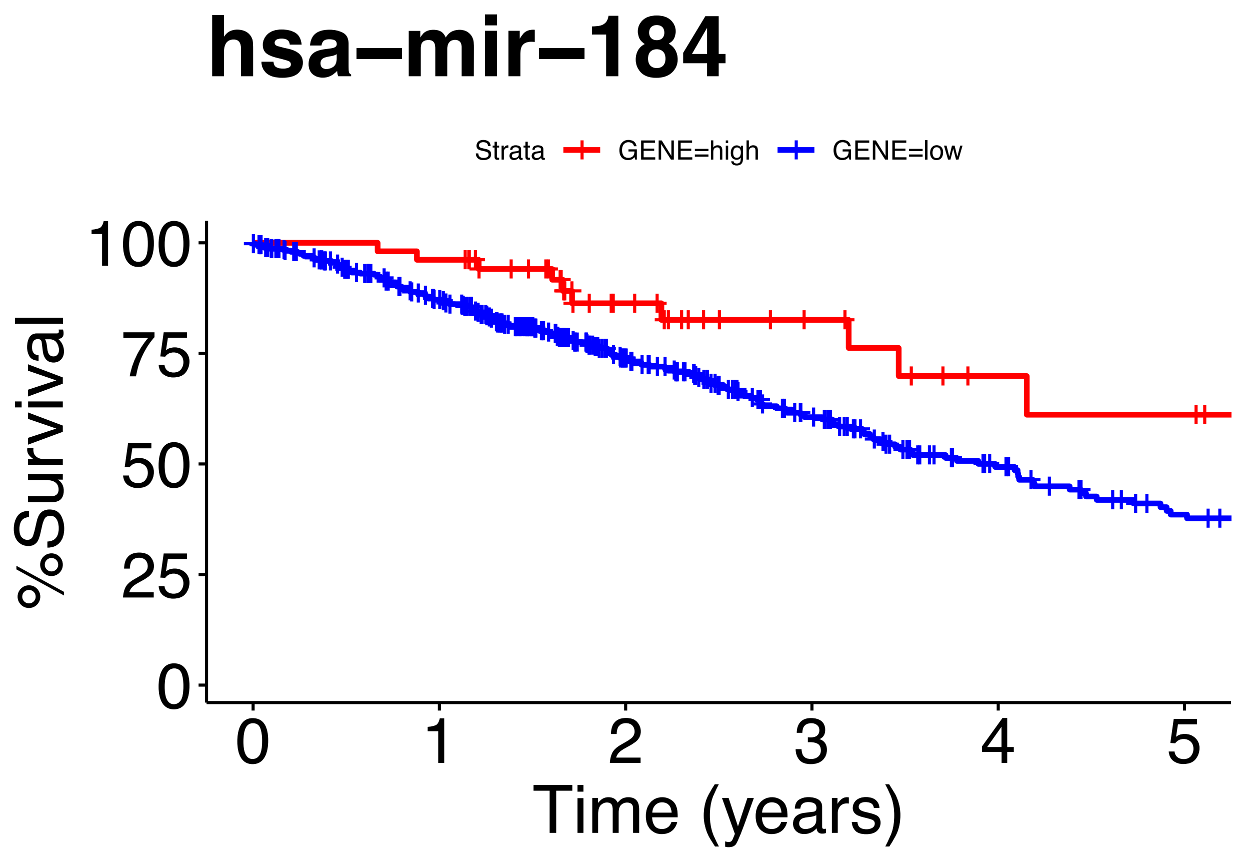


```
ggsurvplot(fit,data = df1.cat,title    = "hsa-mir-184", size = 1, xlim = c(0, 10)  ,ylim=c(1,100), end.
   font.subtitle = c(20, "bold,plain", "black"),
   font.caption = c(25, "plain", "black"),
   font.x = c(25, "plain", "black"),
   font.y = c(25, "plain", "black"),
   font.tickslab = c(24, "plain", "black"), fun = "pct", break.x.by =1)
```

```
## Warning: Vectorized input to `element_text()` is not officially supported.
## Results may be unexpected or may change in future versions of ggplot2.
```

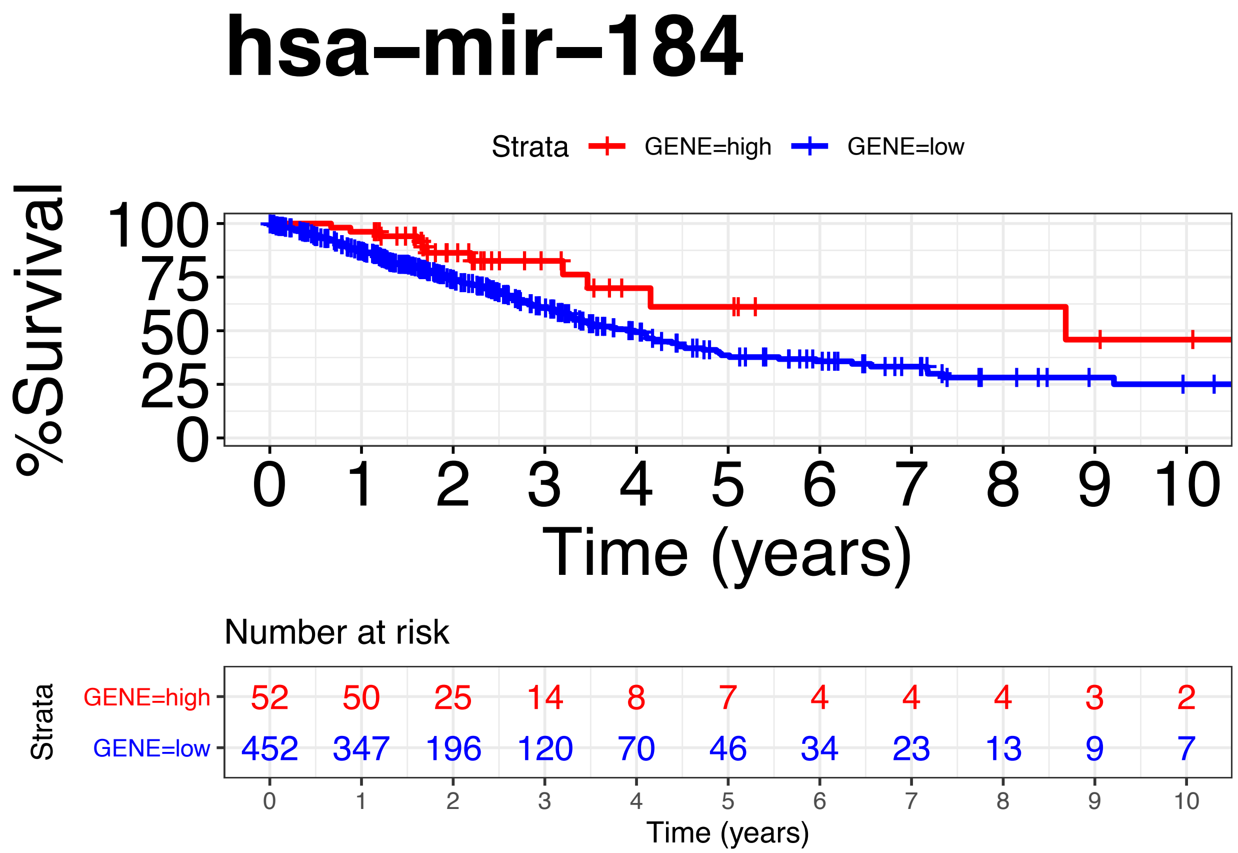


```
ggsurvplot(fit,data = df1.cat,title     = "hsa-mir-184", size = 1,xlim = c(0, 10), ylim = c(0, 100),xlab
   font.subtitle = c(20, "bold.plain", "black"),
   font.caption = c(25, "plain", "black"),
   font.x = c(25, "plain", "black"),
   font.y = c(25, "plain", "black"),
   font.tickslab = c(24, "plain", "black"), fun = "pct", break.x.by =1)
```

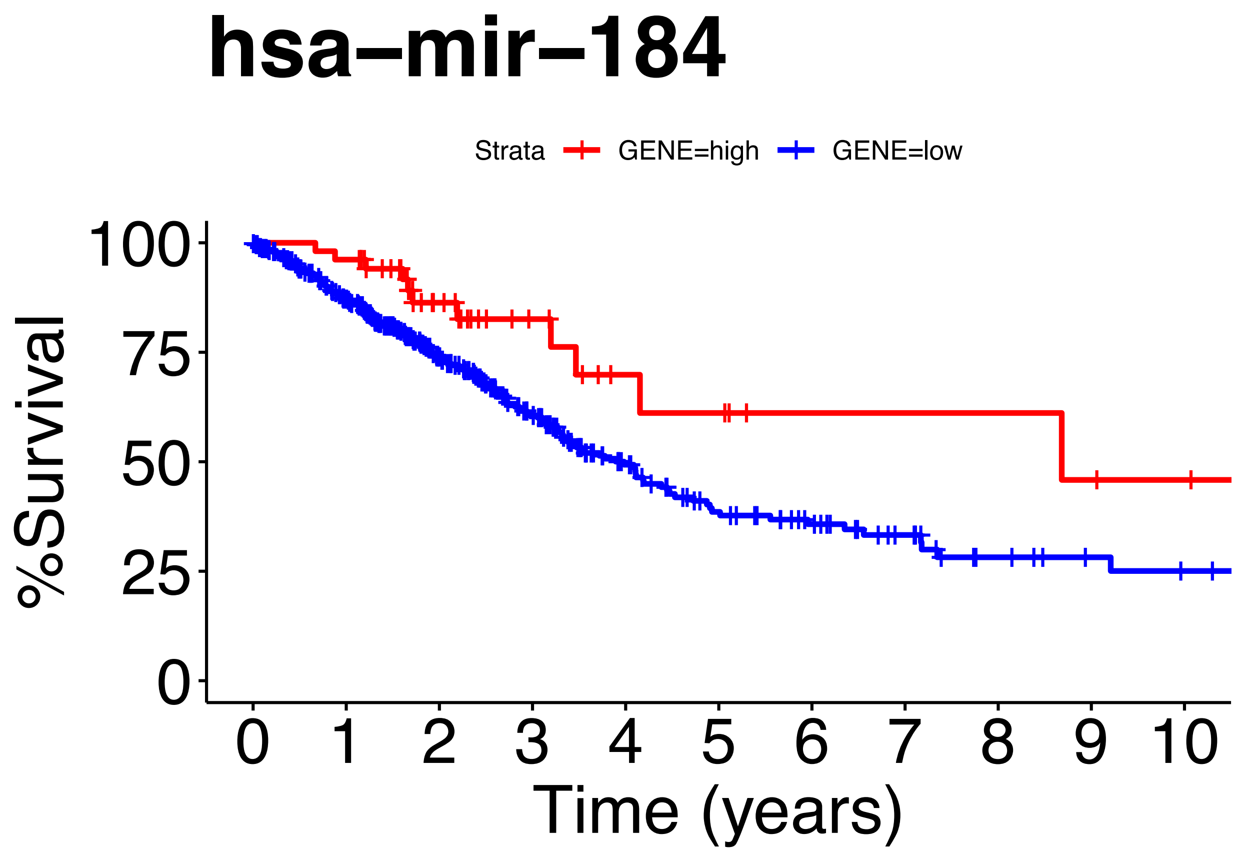


```r
#Cox_Regression
fit.coxph1<- coxph(Surv(os, vital_status) ~ GENE, data=df1.cat)
ggforest(fit.coxph1,data = df1.cat)
```

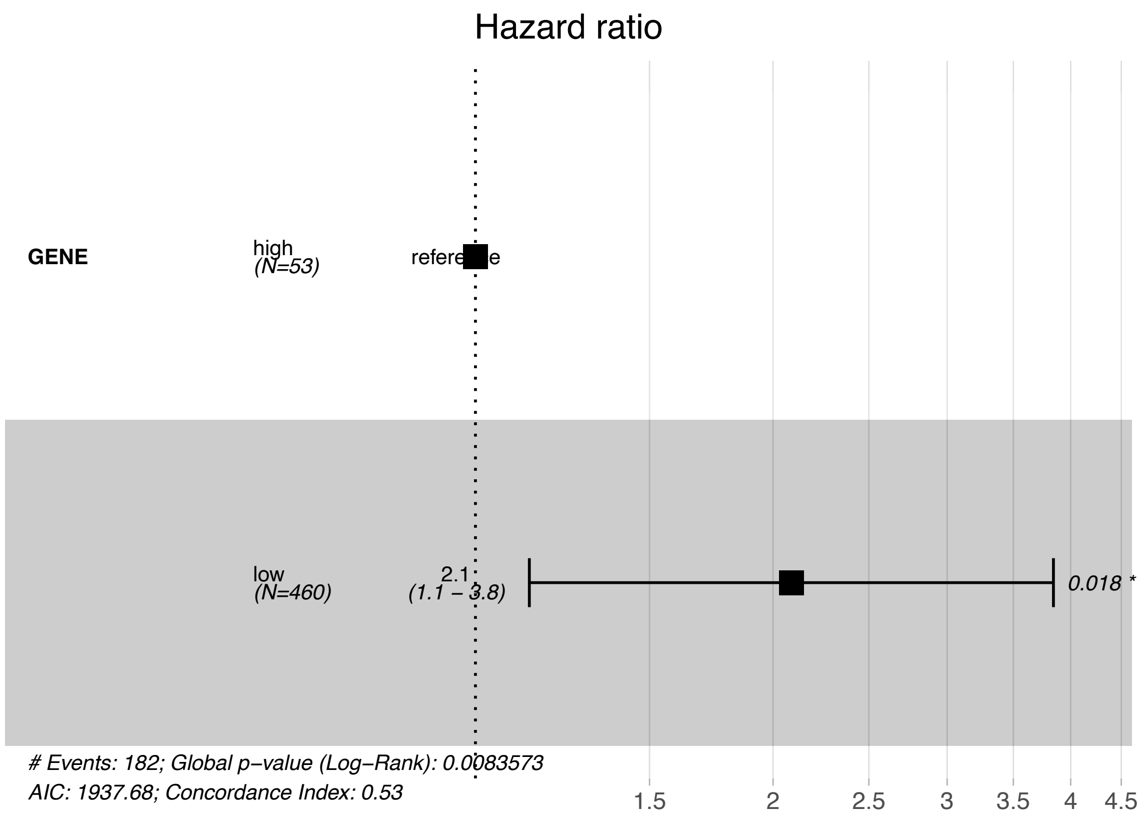


```r
print(fit)
```

```
## Call: survfit(formula = Surv(os, vital_status) ~ GENE, data = df1.cat)
##
##    9 observations deleted due to missingness
##             n events median 0.95LCL 0.95UCL
## GENE=high  52     11   8.68    4.15      NA
## GENE=low  452    171   3.98    3.33    4.53
```

```r
print(fit.coxph1)
```

```
## Call:
## coxph(formula = Surv(os, vital_status) ~ GENE, data = df1.cat)
##
##           coef exp(coef) se(coef)     z      p
## GENElow 0.7359    2.0873   0.3114 2.363 0.0181
##
## Likelihood ratio test=6.96  on 1 df, p=0.008357
## n= 504, number of events= 182
##    (9 observations deleted due to missingness)
```

```r
coxph(Surv(os, vital_status)~ GENE, data=df1.cat) %>%
gtsummary::tbl_regression(exp = TRUE)
```

```
## Table printed with ‘knitr::kable()‘, not {gt}. Learn why at
## http://www.danieldsjoberg.com/gtsummary/articles/rmarkdown.html
## To suppress this message, include ‘message = FALSE‘ in code chunk header.
```

| Characteristic | HR | 95% CI | p-value |
|---|---|---|---|
| GENE | | | |
| high | | | |
| low | 2.09 | 1.13, 3.84 | 0.018 |

```
summary(fit)$table
```

```
##           records n.max n.start events    *rmean *se(rmean)   median  0.95LCL
## GENE=high      52    52      52     11 11.483411   2.204069 8.682192 4.153425
## GENE=low      452   452     452    171  7.023464   0.694736 3.983562 3.328767
##            0.95UCL
## GENE=high       NA
## GENE=low  4.528767
```

#hsa-mir-22

```
#Get mirna data
#Screen the tumor sample barcode from the results: TP (primary solid tumor)

samplesTP <- TCGAquery_SampleTypes(barcode = colnames(miR_matrix),typesample = c("TP"))
mir22_exp <- miR_matrix[c("hsa-mir-22"),samplesTP]
names(mir22_exp) <-  sapply(strsplit(names(mir22_exp),'-'),function(x) paste(x[1:3],collapse="-"))
clinical$GENE <- mir22_exp[clinical$submitter_id]

#Integrate vital status,deaths , last follow up visit.
df2<-subset(clinical,select=c(submitter_id,vital_status,days_to_death,days_to_last_follow_up,GENE))
df2$years_death<- df2$days_to_death/365#convert days to years
df2$years_to_last_follow_up <- df2$days_to_last_follow_up/365
df2$os<-ifelse(df2$vital_status=='Alive',df2$years_to_last_follow_up,df2$years_death)#calculate it as d
df2 <- df2[!is.na(df2$GENE),]#Remove samples with 0 expression
df2[df2$vital_status=='Dead',]$vital_status <- 2
df2[df2$vital_status=='Alive',]$vital_status <- 1
df2$vital_status <- as.numeric(df2$vital_status)

#Determine the optimal cutpoint of variables
df2.cut <- surv_cutpoint(df2,
  time = "os",
  event = "vital_status",
  variables = c("GENE"))
summary(df2.cut)
```

```
##      cutpoint statistic
## GENE 81157.96  1.112411
```

```r
#Categorize variables
df2.cat<-surv_categorize(df2.cut,variables = NULL, labels = c("low", "high"))
head(df2.cat)
```

```
##          os vital_status GENE
## 1 2.5945205            1  low
## 2 2.9616438            2 high
## 3 0.7342466            2  low
## 4 0.8410959            2  low
## 5 0.0000000            2  low
## 6 1.4931507            1 high
```

```r
#get/plot the level of miRNA expression
fit2 <- survfit(Surv(os, vital_status)~GENE, data=df2.cat) #Modeling based on expression.

#plotKM
ggsurvplot(fit2,data = df2.cat,title = "hsa-mir-22", size = 1, xlim = c(0, 5),ylim=c(1,100), risk.table
   font.subtitle = c(20, "bold.plain", "black"),
   font.caption = c(25, "plain", "black"),
   font.x = c(25, "plain", "black"),
   font.y = c(25, "plain", "black"),
   font.tickslab = c(24, "plain", "black"), fun = "pct", break.x.by =1)
```

```
## Warning: Vectorized input to `element_text()` is not officially supported.
## Results may be unexpected or may change in future versions of ggplot2.
```

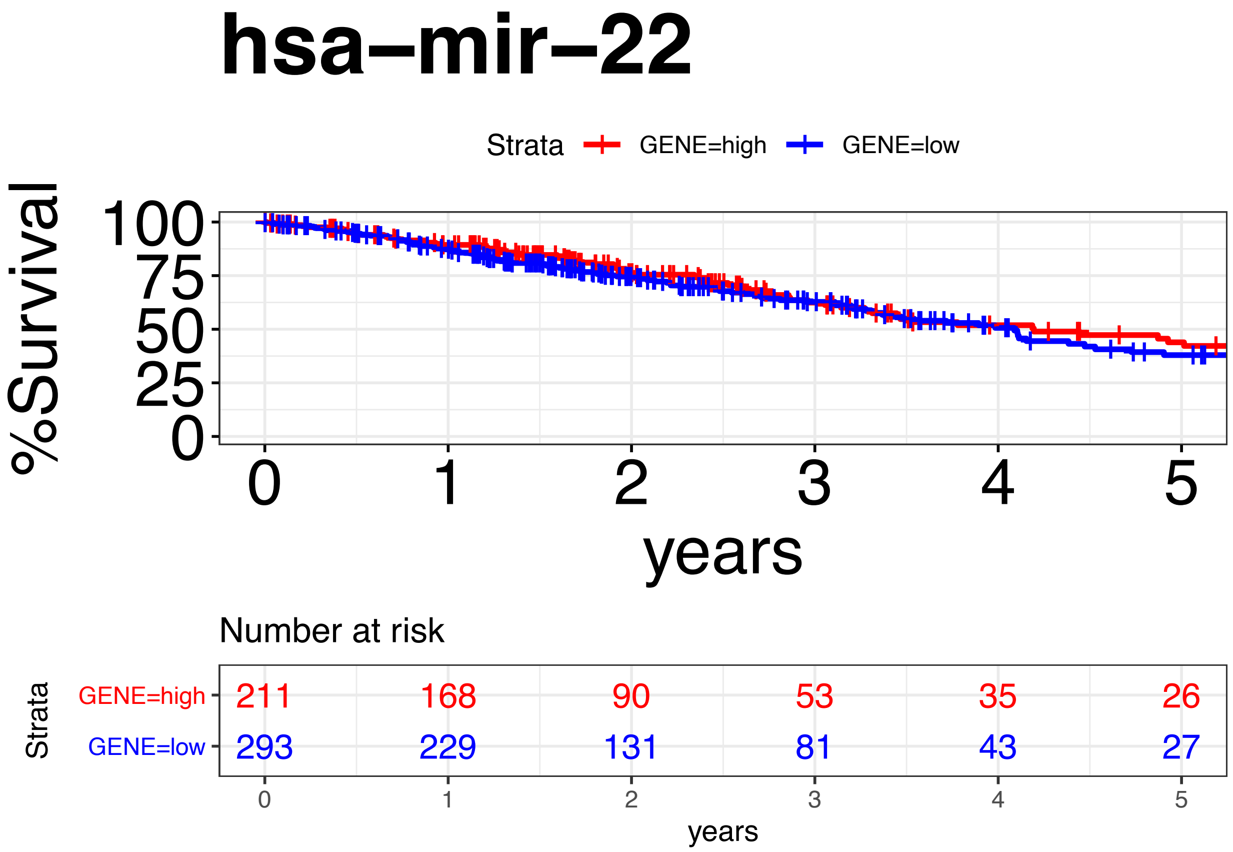

```r
ggsurvplot(fit2,data = df2.cat,title = "hsa-mir-22", size = 1, xlim = c(0, 5),ylim=c(1,100),xlab = "yea
   font.subtitle = c(20, "bold.italic", "black"),
   font.caption = c(25, "plain", "black"),
   font.x = c(25, "plain", "black"),
   font.y = c(25, "plain", "black"),
   font.tickslab = c(24, "plain", "black"), fun = "pct", break.x.by =1)
```

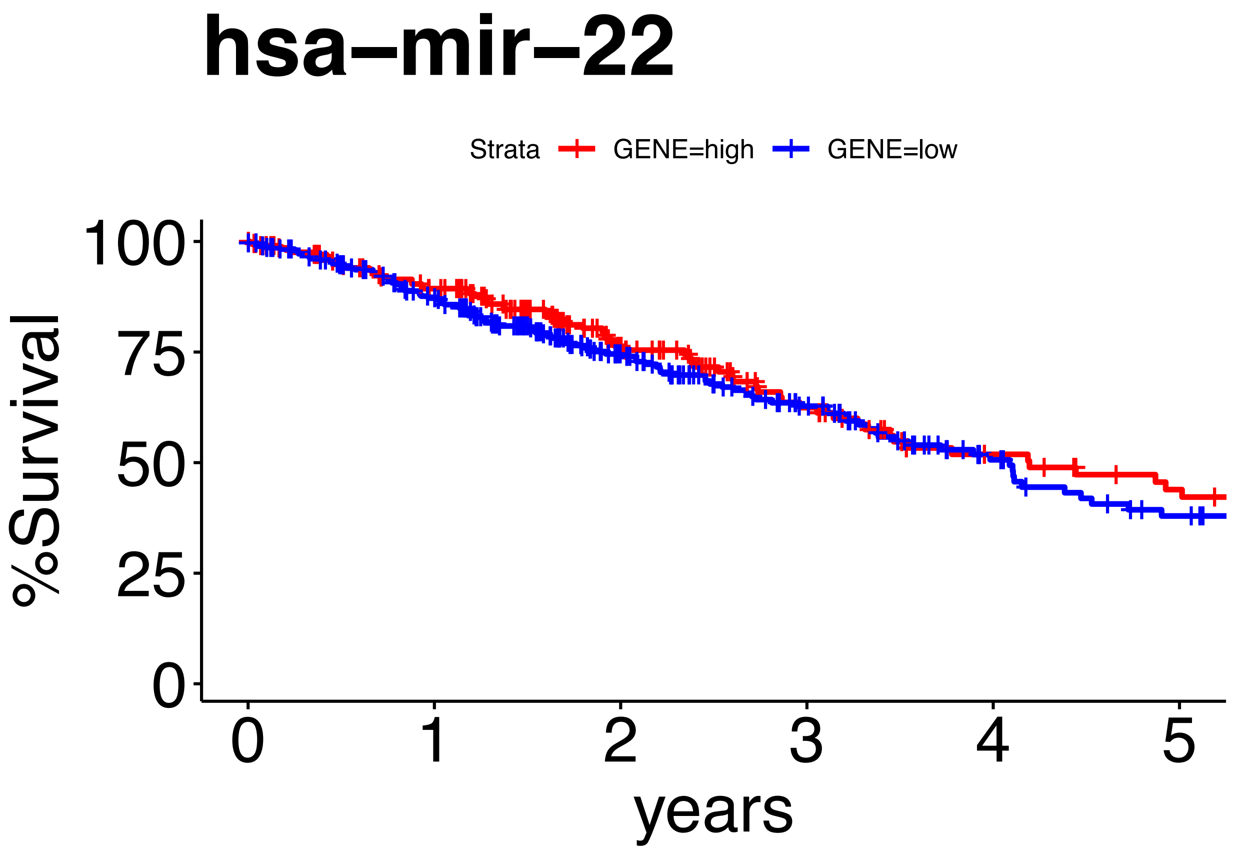


```r
ggsurvplot(fit2,data = df2.cat,title = "hsa-mir-22", size = 1, xlim = c(0, 10),ylim=c(1,100), risk.tabl
   font.subtitle = c(20, "bold.plain", "black"),
   font.caption = c(25, "plain", "black"),
   font.x = c(25, "plain", "black"),
   font.y = c(25, "italic", "black"),
   font.tickslab = c(24, "plain", "black"), fun = "pct", break.x.by =1)
```

```
## Warning: Vectorized input to ‘element_text()‘ is not officially supported.
## Results may be unexpected or may change in future versions of ggplot2.
```

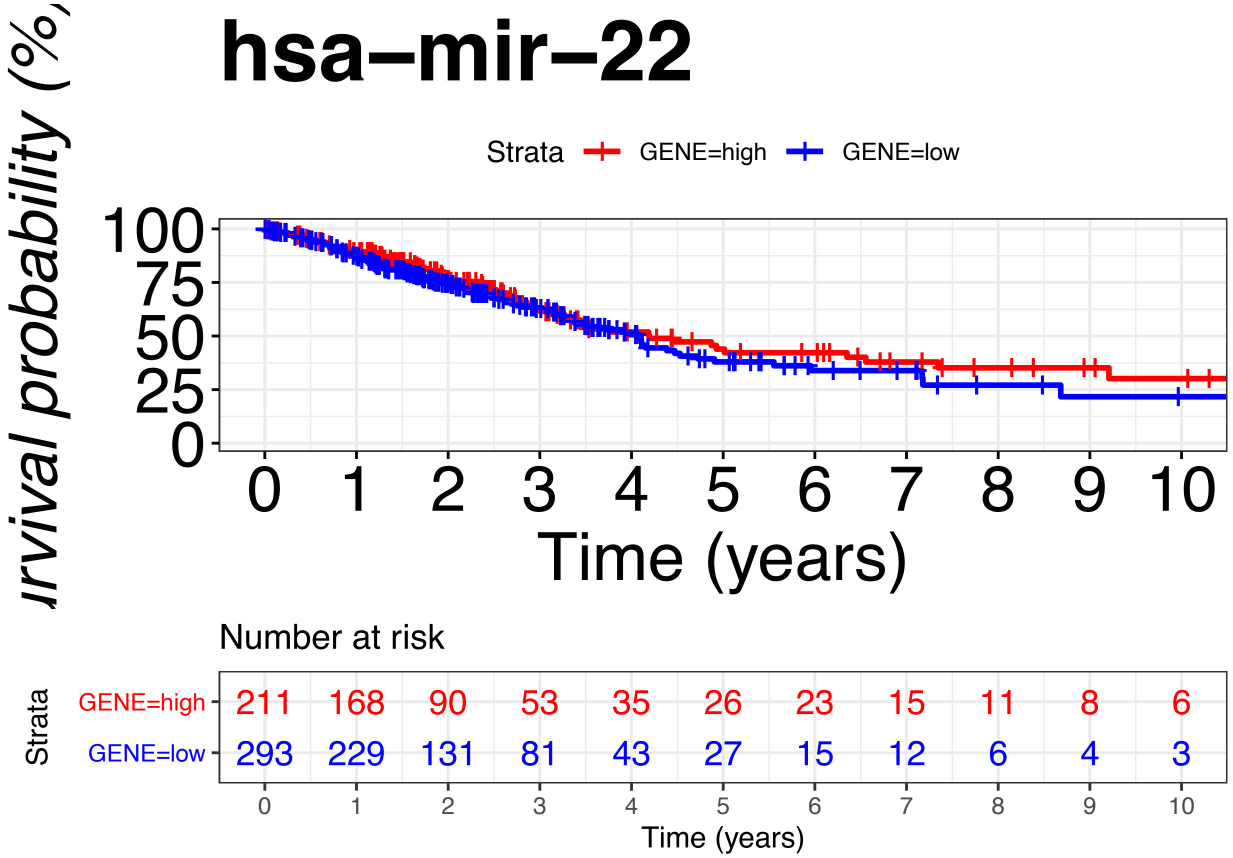


```
ggsurvplot(fit2,data = df2.cat,title = "hsa-mir-22", size = 1, xlim = c(0, 10),ylim=c(1,100),pval=TRUE,
   font.subtitle = c(20, "bold.plain", "black"),
   font.caption = c(25, "plain", "black"),
   font.x = c(25, "plain", "black"),
   font.y = c(25, "plain", "black"),
   font.tickslab = c(24, "plain", "black"), fun = "pct", break.x.by =1)
```

```
## Warning: Removed 1 rows containing missing values (geom_text).

## Warning: Removed 1 rows containing missing values (geom_text).
```

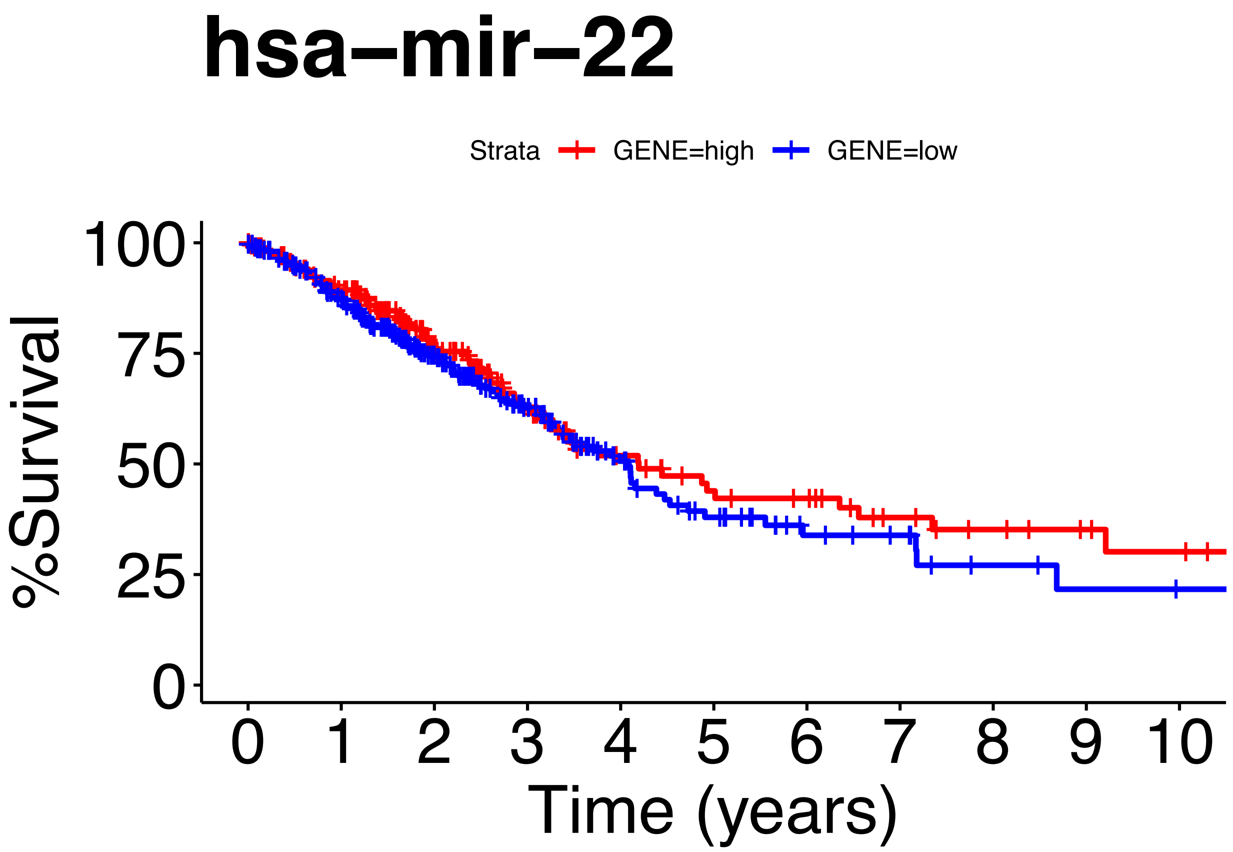


```
#Cox_Regression
fit.coxph2<- coxph(Surv(os, vital_status)~ GENE, data=df2.cat)
ggforest(fit.coxph2,data = df2.cat)
```

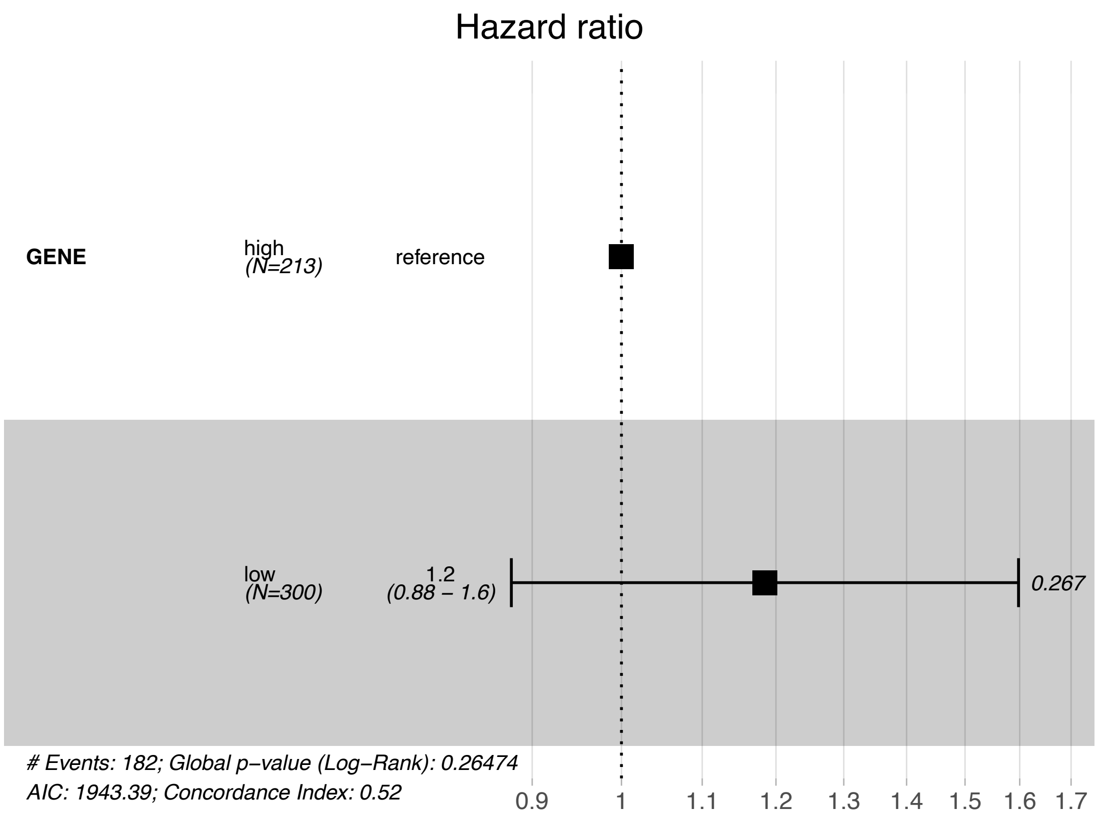


```r
print(fit2)
```

```
## Call: survfit(formula = Surv(os, vital_status) ~ GENE, data = df2.cat)
##
##    9 observations deleted due to missingness
##              n events median 0.95LCL 0.95UCL
## GENE=high 211     71   4.19    3.31    7.35
## GENE=low  293    111   4.09    3.37    4.73
```

```r
print(fit.coxph2)
```

```
## Call:
## coxph(formula = Surv(os, vital_status) ~ GENE, data = df2.cat)
##
##           coef exp(coef) se(coef)     z     p
## GENElow 0.1693    1.1845   0.1527 1.109 0.267
##
## Likelihood ratio test=1.24  on 1 df, p=0.2647
## n= 504, number of events= 182
##    (9 observations deleted due to missingness)
```

```r
coxph(Surv(os, vital_status)~GENE, data=df2.cat) %>%
gtsummary::tbl_regression(exp = TRUE)
```

```
## Table printed with `knitr::kable()`, not {gt}. Learn why at
## http://www.danieldsjoberg.com/gtsummary/articles/rmarkdown.html
## To suppress this message, include `message = FALSE` in code chunk header.
```

| **Characteristic** | **HR** | **95% CI** | **p-value** |
|---|---|---|---|
| GENE | | | |
| high | | | |
| low | 1.18 | 0.88, 1.60 | 0.3 |

```
summary(fit2)$table
```

```
##           records n.max n.start events   *rmean *se(rmean)   median  0.95LCL
## GENE=high     211   211     211     71 8.357805  0.9818366 4.194521 3.312329
## GENE=low      293   293     293    111 6.332918  0.8630399 4.087671 3.367123
##            0.95UCL
## GENE=high 7.345205
## GENE=low  4.726027
```

#hsa-mir-22+184

```
merged6 <- merge(df1.cat,df2.cat)
merged6
```

```
##              os vital_status GENE
## 1    0.00000000            1  low
## 2    0.00000000            1  low
## 3    0.00000000            1  low
## 4    0.00000000            1  low
## 5    0.00000000            1  low
## 6    0.00000000            1  low
## 7    0.00000000            2  low
## 8    0.04109589            1  low
## 9    0.04931507            2  low
## 10   0.06027397            2  low
## 11   0.07671233            1  low
## 12   0.09041096            2  low
## 13   0.09863014            1  low
## 14   0.12054795            1  low
## 15   0.12054795            1  low
## 16   0.13150685            1  low
## 17   0.16986301            1  low
## 18   0.16986301            2  low
## 19   0.21643836            1  low
## 20   0.23013699            1  low
## 21   0.23013699            1  low
## 22   0.23013699            1  low
## 23   0.23013699            1  low
## 24   0.24931507            2  low
## 25   0.26575342            2  low
## 26   0.27123288            2  low
```

```
## 27   0.31780822            2  low
## 28   0.32602740            1  low
## 29   0.32602740            2  low
## 30   0.33150685            2  low
## 31   0.38082192            2  low
## 32   0.38630137            1  low
## 33   0.41369863            1  low
## 34   0.46849315            2  low
## 35   0.47397260            2  low
## 36   0.47671233            1  low
## 37   0.49041096            1  low
## 38   0.49041096            2  low
## 39   0.49863014            1  low
## 40   0.50410959            1  low
## 41   0.50958904            1  low
## 42   0.50958904            1  low
## 43   0.51232877            2  low
## 44   0.52876712            2  low
## 45   0.55342466            1  low
## 46   0.57534247            2  low
## 47   0.61643836            1  low
## 48   0.63013699            1  low
## 49   0.66849315            2  low
## 50   0.66849315            2  low
## 51   0.70410959            2  low
## 52   0.70684932            2  low
## 53   0.72328767            1  low
## 54   0.73424658            2  low
## 55   0.75342466            2  low
## 56   0.76986301            2  low
## 57   0.77260274            2  low
## 58   0.78082192            1  low
## 59   0.78630137            1  low
## 60   0.79726027            2  low
## 61   0.82191781            2  low
## 62   0.83013699            2  low
## 63   0.84109589            1  low
## 64   0.84109589            2  low
## 65   0.84383562            2  low
## 66   0.84931507            1  low
## 67   0.87945205            2 high
## 68   0.87945205            2 high
## 69   0.88493151            1  low
## 70   0.92054795            2  low
## 71   0.92054795            2  low
## 72   0.92054795            2  low
## 73   0.92054795            2  low
## 74   0.92876712            2  low
## 75   0.96164384            1  low
## 76   0.96986301            2  low
## 77   1.00000000            1  low
## 78   1.01369863            2  low
## 79   1.01917808            1  low
## 80   1.02739726            2  low
```

```
## 81   1.03013699            2  low
## 82   1.05479452            1  low
## 83   1.05479452            1  low
## 84   1.05479452            2  low
## 85   1.12054795            2  low
## 86   1.13424658            1  low
## 87   1.13424658            2  low
## 88   1.13698630            1 high
## 89   1.13698630            1 high
## 90   1.14246575            1  low
## 91   1.16164384            1  low
## 92   1.16986301            1  low
## 93   1.17260274            2  low
## 94   1.18904110            2  low
## 95   1.18904110            2  low
## 96   1.19178082            1 high
## 97   1.19178082            1 high
## 98   1.19178082            1  low
## 99   1.19178082            1  low
## 100  1.20547945            2  low
## 101  1.21643836            2  low
## 102  1.21643836            2  low
## 103  1.22191781            1  low
## 104  1.22739726            1  low
## 105  1.22739726            1  low
## 106  1.22739726            1  low
## 107  1.22739726            1  low
## 108  1.24657534            1  low
## 109  1.25205479            2  low
## 110  1.26027397            2  low
## 111  1.28219178            2  low
## 112  1.28219178            2  low
## 113  1.30410959            1  low
## 114  1.30410959            1  low
## 115  1.30410959            1  low
## 116  1.30410959            1  low
## 117  1.30410959            1  low
## 118  1.30410959            1  low
## 119  1.30410959            1  low
## 120  1.30410959            1  low
## 121  1.30410959            1  low
## 122  1.30958904            1  low
## 123  1.30958904            2  low
## 124  1.31780822            1  low
## 125  1.32602740            1  low
## 126  1.33150685            1  low
## 127  1.33424658            1  low
## 128  1.33698630            2  low
## 129  1.34794521            1  low
## 130  1.38356164            1 high
## 131  1.42465753            1  low
## 132  1.44109589            1  low
## 133  1.45479452            1  low
## 134  1.47123288            1  low
```

```
## 135  1.47671233           1 high
## 136  1.48219178           1  low
## 137  1.49863014           1  low
## 138  1.50684932           2  low
## 139  1.51506849           1  low
## 140  1.52602740           2  low
## 141  1.53698630           2  low
## 142  1.54520548           1  low
## 143  1.54794521           1  low
## 144  1.55342466           1  low
## 145  1.55616438           1  low
## 146  1.55616438           1  low
## 147  1.55616438           1  low
## 148  1.55616438           1  low
## 149  1.55616438           1  low
## 150  1.55616438           1  low
## 151  1.55616438           1  low
## 152  1.55616438           1  low
## 153  1.55616438           1  low
## 154  1.57260274           2  low
## 155  1.58356164           1 high
## 156  1.58356164           1 high
## 157  1.58356164           1 high
## 158  1.58356164           1 high
## 159  1.58356164           1  low
## 160  1.61917808           1  low
## 161  1.62191781           1  low
## 162  1.62465753           2  low
## 163  1.65205479           1 high
## 164  1.65205479           1 high
## 165  1.65205479           1 high
## 166  1.65753425           1  low
## 167  1.65753425           1  low
## 168  1.66301370           2 high
## 169  1.66849315           1  low
## 170  1.67123288           1 high
## 171  1.67123288           1  low
## 172  1.67123288           1  low
## 173  1.67123288           1  low
## 174  1.67123288           1  low
## 175  1.69041096           1  low
## 176  1.69041096           1  low
## 177  1.70958904           1 high
## 178  1.71506849           1 high
## 179  1.71506849           1  low
## 180  1.71506849           2  low
## 181  1.72054795           2  low
## 182  1.72602740           1  low
## 183  1.73698630           1  low
## 184  1.78356164           1  low
## 185  1.78356164           1  low
## 186  1.78356164           1  low
## 187  1.78356164           1  low
## 188  1.78630137           1  low
```

```
## 189  1.78630137           1  low
## 190  1.78630137           1  low
## 191  1.78630137           1  low
## 192  1.78904110           1  low
## 193  1.78904110           2  low
## 194  1.80273973           1 high
## 195  1.80273973           1 high
## 196  1.81917808           1  low
## 197  1.82465753           2  low
## 198  1.83287671           1  low
## 199  1.83561644           1  low
## 200  1.83561644           1  low
## 201  1.83561644           1  low
## 202  1.83561644           1  low
## 203  1.85479452           1  low
## 204  1.85479452           2  low
## 205  1.88767123           1  low
## 206  1.89041096           1  low
## 207  1.89315068           1  low
## 208  1.92054795           1 high
## 209  1.92054795           2  low
## 210  1.92876712           1 high
## 211  1.92876712           1 high
## 212  1.92876712           1 high
## 213  1.92876712           1 high
## 214  1.93150685           1 high
## 215  1.93150685           1  low
## 216  1.96712329           1  low
## 217  1.96986301           1  low
## 218  1.98082192           1  low
## 219  1.99452055           1  low
## 220 10.06575342           1 high
## 221 10.79452055           1  low
## 222 13.59178082           2  low
## 223 13.67671233           1 high
## 224 19.85753425           1  low
## 225  2.00273973           2  low
## 226  2.03013699           1  low
## 227  2.08219178           2  low
## 228  2.08493151           1  low
## 229  2.08493151           1  low
## 230  2.08493151           2  low
## 231  2.11780822           1  low
## 232  2.12328767           1  low
## 233  2.12876712           2  low
## 234  2.16712329           1 high
## 235  2.16712329           1  low
## 236  2.16712329           1  low
## 237  2.16712329           1  low
## 238  2.16712329           1  low
## 239  2.16712329           1  low
## 240  2.16712329           1  low
## 241  2.16712329           1  low
## 242  2.16712329           1  low
```

```
## 243  2.16712329            1  low
## 244  2.20547945            1 high
## 245  2.21095890            2  low
## 246  2.21369863            2  low
## 247  2.22739726            1 high
## 248  2.25753425            1  low
## 249  2.25753425            1  low
## 250  2.25753425            1  low
## 251  2.25753425            1  low
## 252  2.26301370            2  low
## 253  2.26575342            1  low
## 254  2.27123288            1  low
## 255  2.27397260            1  low
## 256  2.29863014            1 high
## 257  2.30684932            1  low
## 258  2.31506849            1  low
## 259  2.36438356            1  low
## 260  2.36712329            1  low
## 261  2.38904110            1  low
## 262  2.41643836            1 high
## 263  2.41643836            1  low
## 264  2.45479452            2  low
## 265  2.45479452            2  low
## 266  2.45479452            2  low
## 267  2.45479452            2  low
## 268  2.47945205            2  low
## 269  2.49315068            1  low
## 270  2.49863014            1  low
## 271  2.50136986            1 high
## 272  2.54520548            2  low
## 273  2.54794521            1  low
## 274  2.59452055            1  low
## 275  2.60821918            2  low
## 276  2.67397260            2  low
## 277  2.70410959            2  low
## 278  2.70684932            1  low
## 279  2.72602740            2  low
## 280  2.72602740            2  low
## 281  2.81095890            2  low
## 282  2.83835616            1  low
## 283  2.84931507            1  low
## 284  2.90410959            1  low
## 285  2.93424658            1  low
## 286  2.93698630            1  low
## 287  2.93972603            2  low
## 288  3.00547945            1  low
## 289  3.08219178            1  low
## 290  3.08493151            1  low
## 291  3.08493151            1  low
## 292  3.08493151            1  low
## 293  3.08493151            1  low
## 294  3.10958904            2  low
## 295  3.11232877            2  low
## 296  3.14520548            1  low
```

```
## 297  3.16986301            1  low
## 298  3.20821918            2  low
## 299  3.21917808            1  low
## 300  3.22739726            1  low
## 301  3.25753425            1  low
## 302  3.25753425            1  low
## 303  3.25753425            1  low
## 304  3.25753425            1  low
## 305  3.27945205            2  low
## 306  3.32876712            2  low
## 307  3.36712329            2  low
## 308  3.37808219            1  low
## 309  3.38356164            2  low
## 310  3.46575342            2 high
## 311  3.47397260            2  low
## 312  3.48493151            1  low
## 313  3.52054795            1  low
## 314  3.53150685            1 high
## 315  3.54246575            2  low
## 316  3.56438356            1  low
## 317  3.57534247            1  low
## 318  3.62739726            1  low
## 319  3.65205479            1  low
## 320  3.71780822            2  low
## 321  3.74520548            1  low
## 322  3.75068493            1  low
## 323  3.89315068            2  low
## 324  3.91506849            1  low
## 325  3.92054795            1  low
## 326  3.92328767            1  low
## 327  3.98356164            2  low
## 328  4.03835616            1  low
## 329  4.05205479            1  low
## 330  4.08767123            2  low
## 331  4.10410959            2  low
## 332  4.10684932            2  low
## 333  4.11232877            2  low
## 334  4.17260274            1  low
## 335  4.38356164            2  low
## 336  4.47123288            2  low
## 337  4.52876712            2  low
## 338  4.61095890            1  low
## 339  4.72602740            2  low
## 340  4.73424658            1  low
## 341  4.79452055            1  low
## 342  4.90410959            2  low
## 343  5.12328767            1  low
## 344  5.38356164            1  low
## 345  5.40821918            1  low
## 346  5.55342466            2  low
## 347  5.65753425            1  low
## 348  5.66301370            1  low
## 349  5.77808219            1  low
## 350  5.92054795            1  low
```

```
## 351  5.95616438           2  low
## 352  6.19452055           1  low
## 353  6.48767123           1  low
## 354  6.89041096           1  low
## 355  7.09589041           1  low
## 356  7.10958904           1  low
## 357  7.16986301           2  low
## 358  7.17808219           2  low
## 359  7.33150685           1  low
## 360  7.75890411           1  low
## 361  8.47671233           1  low
## 362  9.05479452           1 high
## 363  9.95890411           1  low
## 364          NA           1 high
## 365          NA           1 high
## 366          NA           1  low
## 367          NA           1  low
## 368          NA           1  low
## 369          NA           1  low
## 370          NA           1  low
## 371          NA           1  low
## 372          NA           1  low
## 373          NA           1  low
## 374          NA           1  low
## 375          NA           1  low
## 376          NA           1  low
## 377          NA           1  low
## 378          NA           2  low
## 379          NA           2  low
## 380          NA           2  low
## 381          NA           2  low
## 382          NA           2  low
## 383          NA           2  low
## 384          NA           2  low
## 385          NA           2  low
## 386          NA           2  low
## 387          NA           2  low
## 388          NA           2  low
## 389          NA           2  low
## 390          NA           2  low
## 391          NA           2  low
## 392          NA           2  low
## 393          NA           2  low
```

```
fit8<- survfit(Surv(os, vital_status)~ GENE, data=merged6)

#Plot_KM
ggsurvplot(fit8,data = merged6,title    = "hsa-mir-22+184", size = 1 ,xlim = c(0, 5), risk.table = TRUE
   font.subtitle = c(20, "bold.plain", "black"),
   font.caption = c(25, "plain", "black"),
   font.x = c(25, "plain", "black"),
   font.y = c(25, "plain", "black"),
   font.tickslab = c(24, "plain", "black"), fun = "pct", break.x.by =1, ylim=c(0,100))
```

```
## Warning: Vectorized input to 'element_text()' is not officially supported.
## Results may be unexpected or may change in future versions of ggplot2.
```

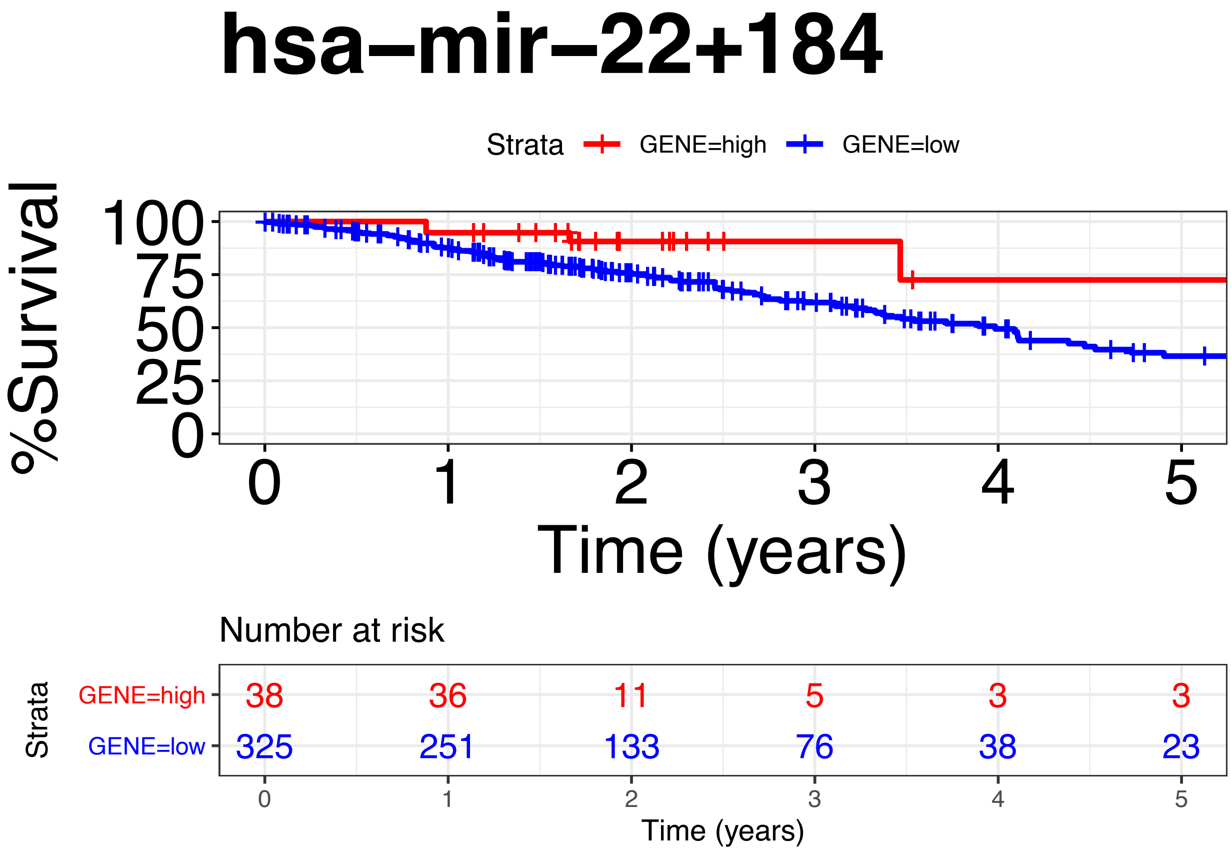


```
ggsurvplot(fit8,data = merged6,title    = "hsa-mir-22+184", size = 1,xlim = c(0, 5),xlab = "Time (years
   font.subtitle = c(20, "bold.plain", "black"),
   font.caption = c(25, "plain", "black"),
   font.x = c(25, "plain", "black"),
   font.y = c(25, "plain", "black"),
   font.tickslab = c(24, "plain", "black"), fun = "pct", break.x.by =1, ylim=c(0,100))
```

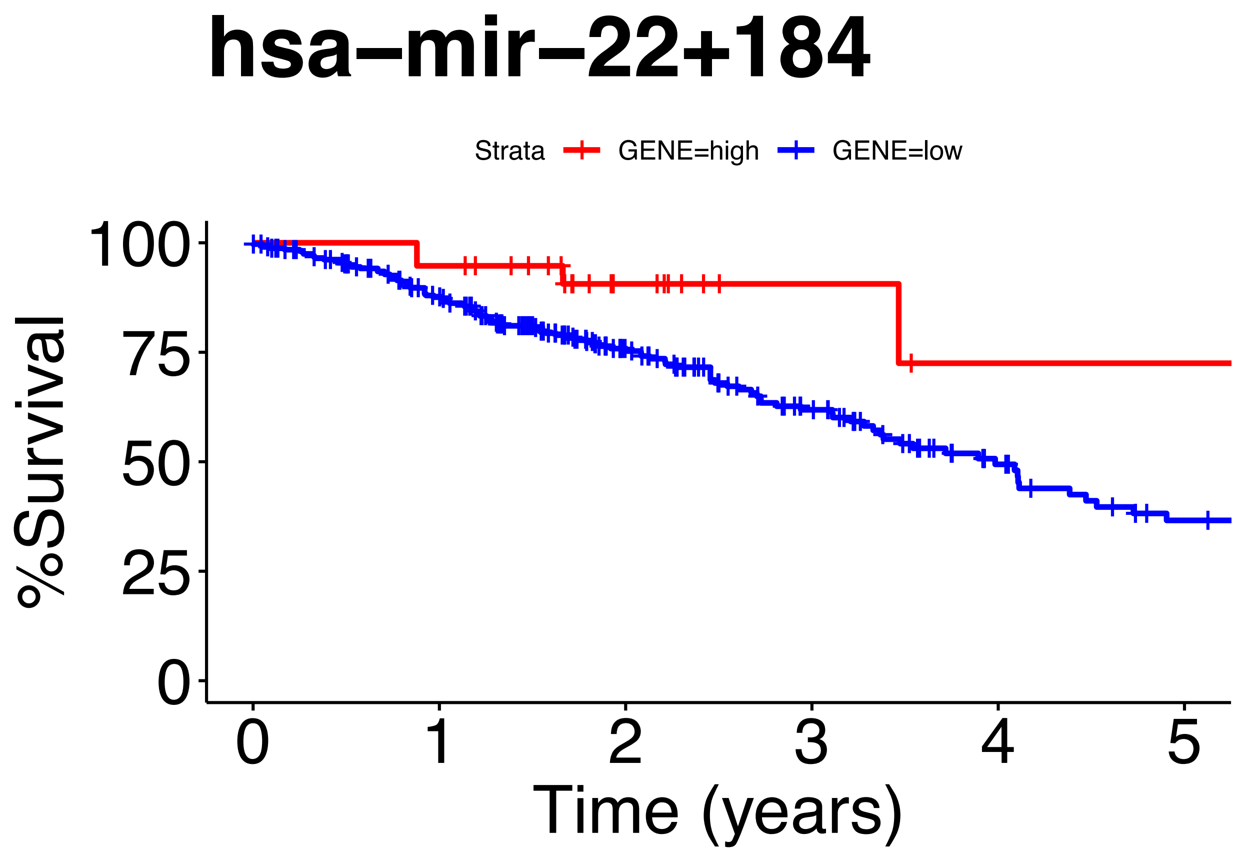


```
ggsurvplot(fit8,data = merged6,title    = "hsa-mir-22+184", size = 1 ,xlim = c(0, 10), risk.table = TRUI
   font.subtitle = c(20, "bold.plain", "black"),
   font.caption = c(25, "plain", "black"),
   font.x = c(25, "plain", "black"),
   font.y = c(25, "plain", "black"),
   font.tickslab = c(24, "plain", "black"), fun = "pct", break.x.by =1, ylim=c(0,100))
```

```
## Warning: Vectorized input to `element_text()` is not officially supported.
## Results may be unexpected or may change in future versions of ggplot2.
```

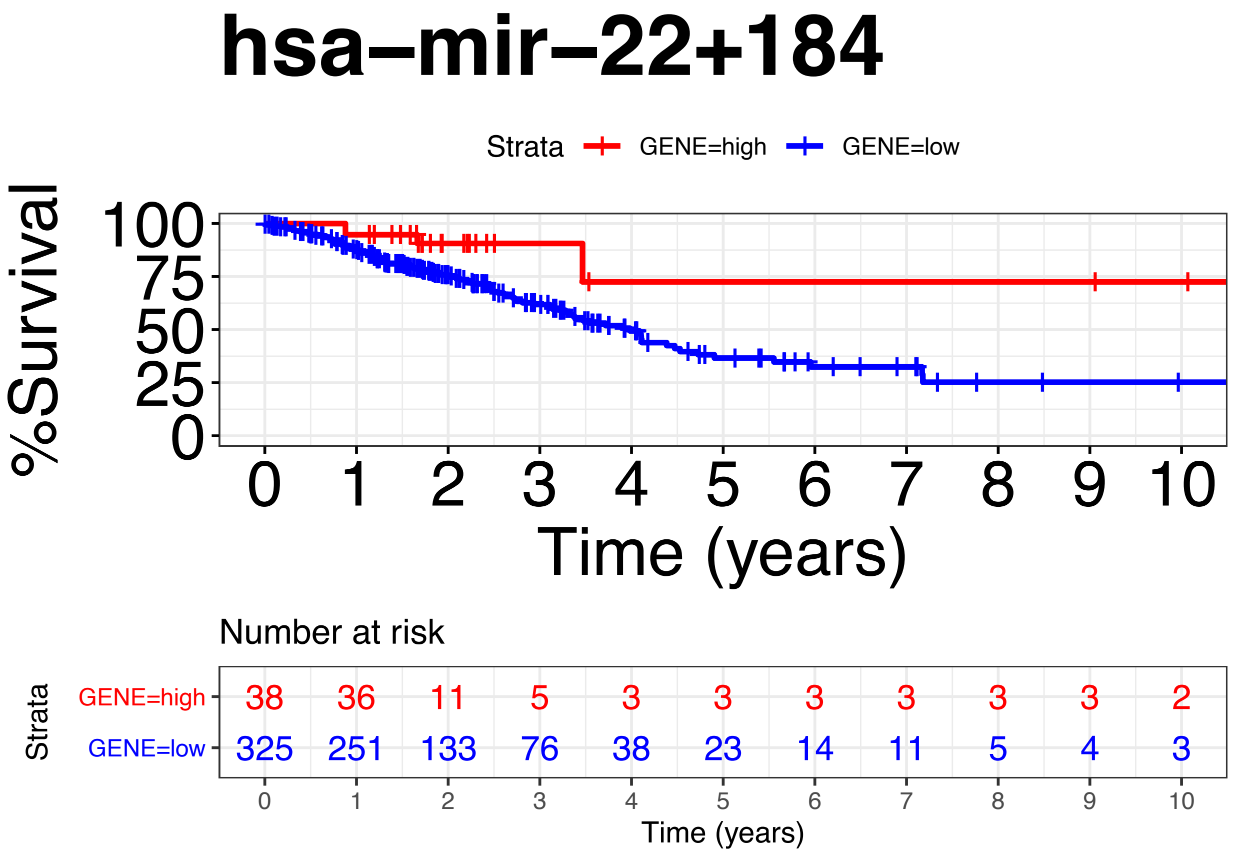


```
ggsurvplot(fit8,data = merged6,title    = "hsa-mir-22+184", size = 1,xlim = c(0, 10), xlab = "Time (yea
   font.subtitle = c(20, "bold.plain", "black"),
   font.caption = c(25, "plain", "black"),
   font.x = c(25, "plain", "black"),
   font.y = c(25, "plain", "black"),
   font.tickslab = c(24, "plain", "black"), fun = "pct", break.x.by =1, ylim=c(0,100))
```

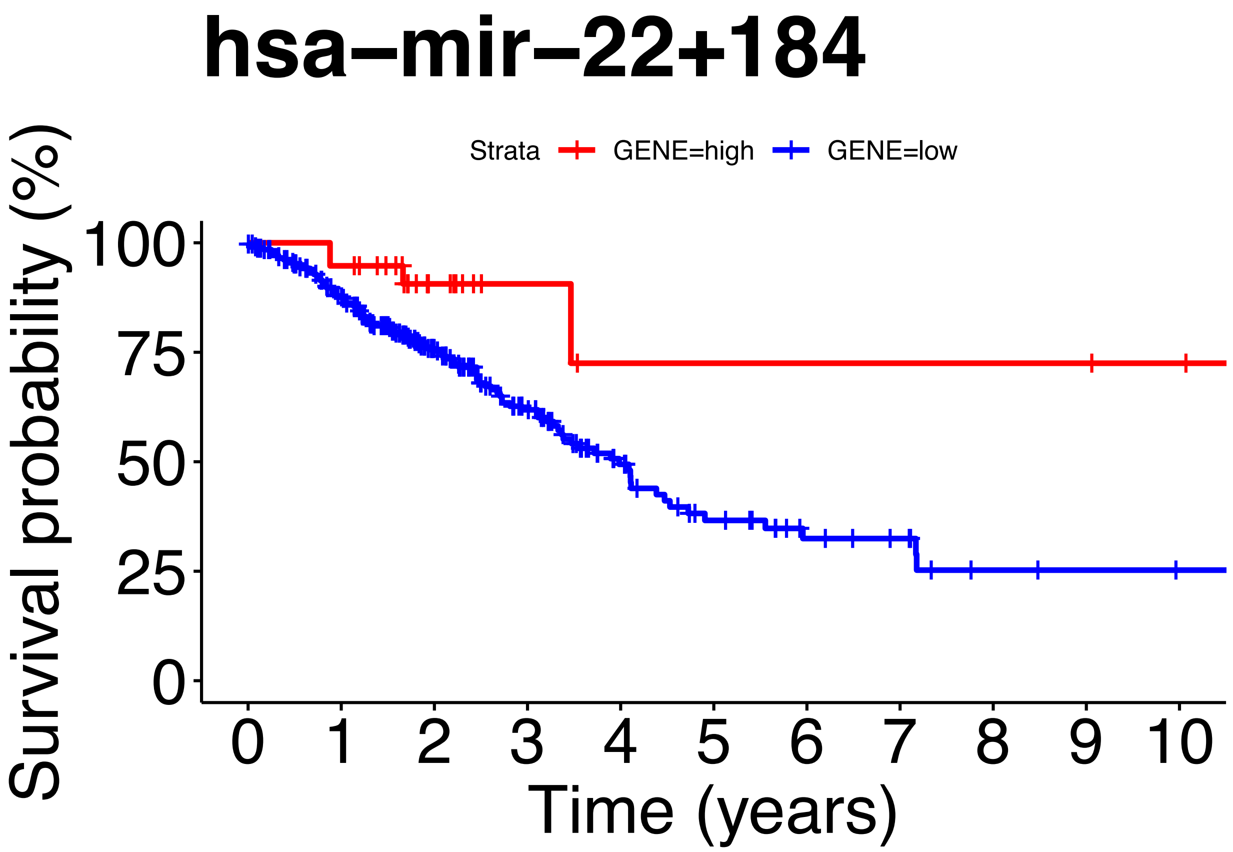


```r
#Cox_Regression
fit.coxph1<- coxph(Surv(os, vital_status)~ GENE, data=merged6)
ggforest(fit.coxph1,data = merged6)
```

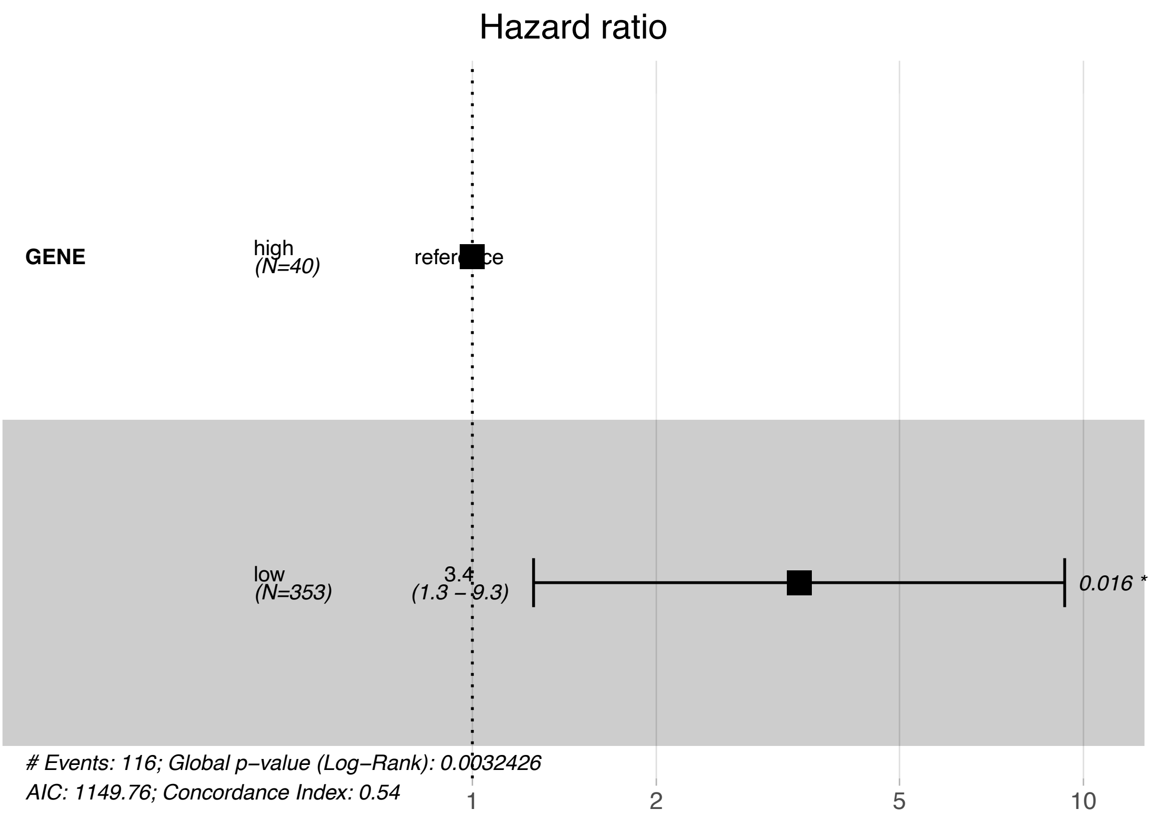


```r
print(fit8)
```

```
## Call: survfit(formula = Surv(os, vital_status) ~ GENE, data = merged6)
##
##    30 observations deleted due to missingness
##              n events median 0.95LCL 0.95UCL
## GENE=high  38      4     NA    3.47      NA
## GENE=low  325    112   3.98    3.37    4.73
```

```r
print(fit.coxph1)
```

```
## Call:
## coxph(formula = Surv(os, vital_status) ~ GENE, data = merged6)
##
##           coef exp(coef) se(coef)     z      p
## GENElow 1.2313    3.4255   0.5105 2.412 0.0159
##
## Likelihood ratio test=8.67  on 1 df, p=0.003243
## n= 363, number of events= 116
##    (30 observations deleted due to missingness)
```

```r
coxph(Surv(os, vital_status)~ GENE, data=merged6) %>%
gtsummary::tbl_regression(exp = TRUE)
```

```
## Table printed with `knitr::kable()`, not {gt}. Learn why at
## http://www.danieldsjoberg.com/gtsummary/articles/rmarkdown.html
## To suppress this message, include `message = FALSE` in code chunk header.
```

| Characteristic | HR | 95% CI | p-value |
|---|---|---|---|
| GENE | | | |
| high | | | |
| low | 3.43 | 1.26, 9.32 | 0.016 |

```r
summary(fit8)$table
```

```
##           records n.max n.start events    *rmean *se(rmean)   median  0.95LCL
## GENE=high      38    38      38      4 15.138481  2.7782600       NA 3.465753
## GENE=low      325   325     325    112  6.559784  0.9027472 3.983562 3.367123
##            0.95UCL
## GENE=high       NA
## GENE=low  4.726027
```

**Combining Plasma Extracellular Vesicle Let-7b-5p, miR-184 and Circulating miR-22-3p Levels for NSCLC Diagnosis and for Predicting Drug Resistance**

G. P. Vadla[1], B. Daghat[1], A. Garcia[1*], G. Perez[1*], V. Ahmad[1], N. Patterson[1], Y. Manjunath[2], J.T. Kaifi[2,3], G. Li[2], and C.Y. Chabu[1,2,3‡]

### Supplementary information 2. R session information for covariate analyses

```
## – Session info ————————————————————————————
##  setting  value
##  version  R version 4.1.0 (2021-05-18)
##  os       Ubuntu 20.04.2 LTS
##  system   x86_64, linux-gnu
##  ui       X11
##  language (EN)
##  collate  en_US.UTF-8
##  ctype    en_US.UTF-8
##  tz       Etc/UTC
##  date     2021-08-05

## – Packages ————————————————————————————
package        * version date       lib source
##  abind          1.4-5   2016-07-21 [1] RSPM (R 4.0.3)
##  assertthat     0.2.1   2019-03-21 [1] RSPM (R 4.1.0)
##  backports      1.2.1   2020-12-09 [1] RSPM (R 4.1.0)
##  Biobase      * 2.52.0  2021-05-19 [1] Bioconductor
##  BiocGenerics * 0.38.0  2021-05-19 [1] Bioconductor
##  broom          0.7.9   2021-07-27 [1] RSPM (R 4.0.5)
##  bslib          0.2.5.1 2021-05-18 [1] RSPM (R 4.1.0)
##  cachem         1.0.5   2021-05-15 [1] RSPM (R 4.1.0)
##  callr          3.7.0   2021-04-20 [1] RSPM (R 4.1.0)
##  car            3.0-11  2021-06-27 [1] RSPM (R 4.0.5)
##  carData        3.0-4   2020-05-22 [1] RSPM (R 4.0.3)
##  cellranger     1.1.0   2016-07-27 [1] RSPM (R 4.1.0)
##  cli            3.0.1   2021-07-17 [1] RSPM (R 4.0.5)
##  colorspace     2.0-2   2021-06-24 [1] RSPM (R 4.1.0)
##  crayon         1.4.1   2021-02-08 [1] RSPM (R 4.1.0)
##  crosstalk      1.1.1   2021-01-12 [1] RSPM (R 4.0.3)
##  curl           4.3.2   2021-06-23 [1] RSPM (R 4.1.0)
##  data.table     1.14.0  2021-02-21 [1] RSPM (R 4.1.0)
##  DBI            1.1.1   2021-01-15 [1] RSPM (R 4.1.0)
##  desc           1.3.0   2021-03-05 [1] RSPM (R 4.1.0)
##  devtools       2.4.2   2021-06-07 [1] RSPM (R 4.1.0)
##  digest         0.6.27  2020-10-24 [1] RSPM (R 4.1.0)
##  dplyr        * 1.0.7   2021-06-18 [1] RSPM (R 4.1.0)
##  DT           * 0.18    2021-04-14 [1] RSPM (R 4.0.4)
```

```
## edgeR        * 3.34.0  2021-05-19 [1] Bioconductor
## ellipsis       0.3.2   2021-04-29 [1] RSPM (R 4.1.0)
## evaluate       0.14    2019-05-28 [1] RSPM (R 4.1.0)
## fansi          0.5.0   2021-05-25 [1] RSPM (R 4.1.0)
## farver         2.1.0   2021-02-28 [1] RSPM (R 4.1.0)
## fastmap        1.1.0   2021-01-25 [1] RSPM (R 4.1.0)
## forcats        0.5.1   2021-01-27 [1] RSPM (R 4.1.0)
## foreign        0.8-81  2020-12-22 [2] CRAN (R 4.1.0)
## fs             1.5.0   2020-07-31 [1] RSPM (R 4.1.0)
## generics       0.1.0   2020-10-31 [1] RSPM (R 4.1.0)
## GGally       * 2.1.2   2021-06-21 [1] RSPM (R 4.0.5)
## ggplot2      * 3.3.5   2021-06-25 [1] RSPM (R 4.1.0)
## ggpubr       * 0.4.0   2020-06-27 [1] RSPM (R 4.0.3)
## ggrepel      * 0.9.1   2021-01-15 [1] RSPM (R 4.0.5)
## ggsignif       0.6.2   2021-06-14 [1] RSPM (R 4.0.5)
## glue           1.4.2   2020-08-27 [1] RSPM (R 4.1.0)
## gtable         0.3.0   2019-03-25 [1] RSPM (R 4.1.0)
## haven          2.4.1   2021-04-23 [1] RSPM (R 4.1.0)
## highr          0.9     2021-04-16 [1] RSPM (R 4.1.0)
## hms            1.1.0   2021-05-17 [1] RSPM (R 4.1.0)
## htmltools      0.5.1.1 2021-01-22 [1] RSPM (R 4.1.0)
## htmlwidgets    1.5.3   2020-12-10 [1] RSPM (R 4.1.0)
## httr           1.4.2   2020-07-20 [1] RSPM (R 4.1.0)
## jquerylib      0.1.4   2021-04-26 [1] RSPM (R 4.1.0)
## jsonlite       1.7.2   2020-12-09 [1] RSPM (R 4.1.0)
## kableExtra   * 1.3.4   2021-02-20 [1] RSPM (R 4.0.3)
## knitr          1.33    2021-04-24 [1] RSPM (R 4.1.0)
## labeling       0.4.2   2020-10-20 [1] RSPM (R 4.1.0)
## lattice        0.20-44 2021-05-02 [2] CRAN (R 4.1.0)
## lazyeval       0.2.2   2019-03-15 [1] RSPM (R 4.0.3)
## lifecycle      1.0.0   2021-02-15 [1] RSPM (R 4.1.0)
## limma        * 3.48.1  2021-06-24 [1] Bioconductor
## locfit         1.5-9.4 2020-03-25 [1] RSPM (R 4.0.3)
## magrittr     * 2.0.1   2020-11-17 [1] RSPM (R 4.1.0)
## Matrix         1.3-3   2021-05-04 [2] CRAN (R 4.1.0)
## memoise        2.0.0   2021-01-26 [1] RSPM (R 4.1.0)
## mgcv           1.8-35  2021-04-18 [2] CRAN (R 4.1.0)
## mime           0.11    2021-06-23 [1] RSPM (R 4.1.0)
## munsell        0.5.0   2018-06-12 [1] RSPM (R 4.1.0)
## nlme           3.1-152 2021-02-04 [2] CRAN (R 4.1.0)
## openxlsx       4.2.4   2021-06-16 [1] RSPM (R 4.0.5)
## pcaMethods   * 1.84.0  2021-05-19 [1] Bioconductor
## pheatmap     * 1.0.12  2019-01-04 [1] RSPM (R 4.0.3)
```

```
## pillar        1.6.1   2021-05-16 [1] RSPM (R 4.1.0)
## pkgbuild      1.2.0   2020-12-15 [1] RSPM (R 4.1.0)
## pkgconfig     2.0.3   2019-09-22 [1] RSPM (R 4.1.0)
## pkgload       1.2.1   2021-04-06 [1] RSPM (R 4.1.0)
## plotly      * 4.9.4.1 2021-06-18 [1] RSPM (R 4.0.5)
## plyr        * 1.8.6   2020-03-03 [1] RSPM (R 4.0.5)
## prettyunits   1.1.1   2020-01-24 [1] RSPM (R 4.1.0)
## processx      3.5.2   2021-04-30 [1] RSPM (R 4.1.0)
## ps            1.6.0   2021-02-28 [1] RSPM (R 4.1.0)
## purrr         0.3.4   2020-04-17 [1] RSPM (R 4.1.0)
## R6            2.5.0   2020-10-28 [1] RSPM (R 4.1.0)
## RColorBrewer * 1.1-2   2014-12-07 [1] RSPM (R 4.1.0)
## Rcpp          1.0.7   2021-07-07 [1] RSPM (R 4.1.0)
## readr       * 2.0.0   2021-07-20 [1] RSPM (R 4.0.5)
## readxl      * 1.3.1   2019-03-13 [1] RSPM (R 4.1.0)
## remotes       2.4.0   2021-06-02 [1] RSPM (R 4.1.0)
## reshape       0.8.8   2018-10-23 [1] RSPM (R 4.0.3)
## rio           0.5.27  2021-06-21 [1] RSPM (R 4.0.5)
## rlang         0.4.11  2021-04-30 [1] RSPM (R 4.1.0)
## rmarkdown     2.9     2021-06-15 [1] RSPM (R 4.1.0)
## ROCR        * 1.0-11  2020-05-02 [1] RSPM (R 4.0.0)
## rprojroot     2.0.2   2020-11-15 [1] RSPM (R 4.1.0)
## rstatix       0.7.0   2021-02-13 [1] RSPM (R 4.0.3)
## rstudioapi    0.13    2020-11-12 [1] RSPM (R 4.1.0)
## Rtsne       * 0.15    2018-11-10 [1] RSPM (R 4.0.5)
## rvest         1.0.1   2021-07-26 [1] RSPM (R 4.0.5)
## sass          0.4.0   2021-05-12 [1] RSPM (R 4.1.0)
## scales        1.1.1   2020-05-11 [1] RSPM (R 4.1.0)
## sessioninfo   1.1.1   2018-11-05 [1] RSPM (R 4.1.0)
## stringi     * 1.7.3   2021-07-16 [1] RSPM (R 4.0.5)
## stringr     * 1.4.0   2019-02-10 [1] RSPM (R 4.1.0)
## svglite       2.0.0   2021-02-20 [1] RSPM (R 4.0.5)
## systemfonts   1.0.2   2021-05-11 [1] RSPM (R 4.0.5)
## testthat      3.0.4   2021-07-01 [1] RSPM (R 4.1.0)
## tibble      * 3.1.3   2021-07-23 [1] RSPM (R 4.0.5)
## tidyr       * 1.1.3   2021-03-03 [1] RSPM (R 4.1.0)
## tidyselect    1.1.1   2021-04-30 [1] RSPM (R 4.1.0)
## tzdb          0.1.2   2021-07-20 [1] RSPM (R 4.0.5)
## usethis       2.0.1   2021-02-10 [1] RSPM (R 4.1.0)
## utf8          1.2.2   2021-07-24 [1] RSPM (R 4.0.5)
## vctrs         0.3.8   2021-04-29 [1] RSPM (R 4.1.0)
## viridisLite   0.4.0   2021-04-13 [1] RSPM (R 4.1.0)
## webshot       0.5.2   2019-11-22 [1] RSPM (R 4.1.0)
```

```
##  withr          2.4.2   2021-04-18 [1] RSPM (R 4.1.0)
##  WriteXLS     * 6.3.0   2021-04-01 [1] RSPM (R 4.0.4)
##  xfun           0.24    2021-06-15 [1] RSPM (R 4.1.0)
##  xml2           1.3.2   2020-04-23 [1] RSPM (R 4.1.0)
##  yaml           2.2.1   2020-02-01 [1] RSPM (R 4.1.0)
##  zip            2.2.0   2021-05-31 [1] RSPM (R 4.1.0)
##
## [1] /usr/local/lib/R/site-library
## [2] /usr/local/lib/R/library
```
